# Frequency-dependent coupling in response to oscillatory inputs in networks of electrically coupled nodes: Gap junction networks and spatially extended neurons - Full Version

**DOI:** 10.1101/2025.09.12.675827

**Authors:** Andrea Bel, Ulises Chialva, Horacio G. Rotstein

**Affiliations:** Departamento de Matemática, Universidad Nacional del Sur (UNS) and CONICET, Bahía Blanca, Argentina; Federated Department of Biological Sciences, New Jersey Institute of Technology and Rutgers University, Newark, NJ, USA

## Abstract

In electrically coupled networks, the coupling coefficient (CC) quantifies the strength of the connectivity or the degree to which two participating nodes are coupled in response to an external input to one of them. The CC is measured by computing the relative responses of the indirectly activated (post-J) and the directly activated (pre-J) nodes. In response to time-dependent inputs, the CC is frequency-dependent and has two components capturing the contributions of the amplitude and phase frequency profiles of the participating nodes (quotient of the amplitudes and phase-difference, respectively). The properties and mechanisms of generation of the frequency-dependent CCs (FD-CCs) are largely unknown beyond electrically coupled passive cells and their electrical circuit equivalents. Being linear and 1D, the FD-CCs for passive cells are relatively simple, consisting of low-pass filters (amplitude) and positive and monotonically increasing phase-difference profiles. In linear systems, the FD-CCs depend on the properties of the pre-J cell and the connectivity and are independent of the properties of the post-J cell and the input amplitude. There is a gap in our understanding of the FD-CCs are shaped by (i) how the presence of intrinsic cellular positive and negative feedback currents and the resulting amplification and resonance phenomena, and (ii) the presence of cellular nonlinearities that incorporates the dependence of the FD-CC on the post-J node in addition to the pre-J one. In this paper we address these issues by using biophysically plausible (conductance-based) mathematical modeling, numerical simulations, analytical calculations and dynamical systems tools. We conduct a systematic analysis of the properties of the FD-CC in networks of two electrically connected nodes receiving oscillatory inputs, which is the minimal network architecture that allows for a systematic study of the biophysical and dynamic mechanisms that shape the FD-CC profiles. The participating neurons are either passive cells (low-pass filters) or resonators (band-pass filter) and exhibit lagging or mixed leading-lagging phase-shift responses as the input frequency increases. The formalism and tools we develop and use in this paper can be extended to larger networks with an arbitrary number of nodes, to spatially extended multicompartment neuronal models, and to neurons having a variety of ionic currents. The principles that emerge from our study are directly applicable to these scenarios. Our results make experimentally testable predictions and have implications for the understanding of spike transmission, synchronized firing and coincidence detection in electrically coupled networks in the presence of oscillatory inputs. For clarity, the paper includes an extensive supplementary material section.

## 1 Introduction

Neuronal circuits are impacted by external inputs that often have oscillatory components spanning a range of frequencies (*f*). The frequency-dependent output is shaped by the complex interaction between the input and the cellular and synaptic properties. Neuronal filters describe the information processing building blocks where specific frequency components of the output are enhanced over others [1–6]. Therefore neuronal filters play a crucial role in shaping neuronal communication and how the brain process information, generates rhythmic activity and performs complex computations [3, 4, 7–18].

In this paper we investigate the mechanism of interaction of cellular neuronal filters in electrically coupled networks. We use biophysically plausible (conductance-based) mathematical modeling, numerical simulations, analytical calculations and dynamical systems tools. We focus on gap junction connected cells and compartmental models of spatially extended neurons. While, biologically, these are different objects, they belong to the same family of mathematical models of electrically coupled nodes that express neuronal amplitude and phase filters and are subject to inputs with frequency components within certain range, and therefore can be investigated within the same framework.

Two of the most common types of neuronal filters are low-pass (LPFs; e.g., Figs. 1-A1, -B1) and band-pass (BPFs; e.g., Figs. 1-A2, -B2) filters. BPFs are closely linked to the concept of neuronal resonance [1, 4, 19], defined as the ability of a neuronal system to exhibit a maximal response (e.g., subthreshold membrane potential) to periodic inputs at a preferred (resonant), non-zero frequency (or frequency band). In contrast, LPFs describe a preference for inputs in the lowest frequency band. The associated frequency-dependent phase (phase-shift) profiles can be entirely positive (e.g., Figs. 1-C1), indicating a lagging response for all input frequencies, or transition from negative to positive as the input frequency increases (e.g., Figs. 1-C2), indicating a mixed leading-lagging response and exhibiting phasonance at the (nonzero) frequency at which the phase vanishes. In both cases, the phase monotonically increases for frequencies above the phasonant frequency. Resonance and phasonance are related phenomena, and when they coexist, the resonant and phasonant frequency do not necessarily coincide [20, 21].

**Figure 1:**
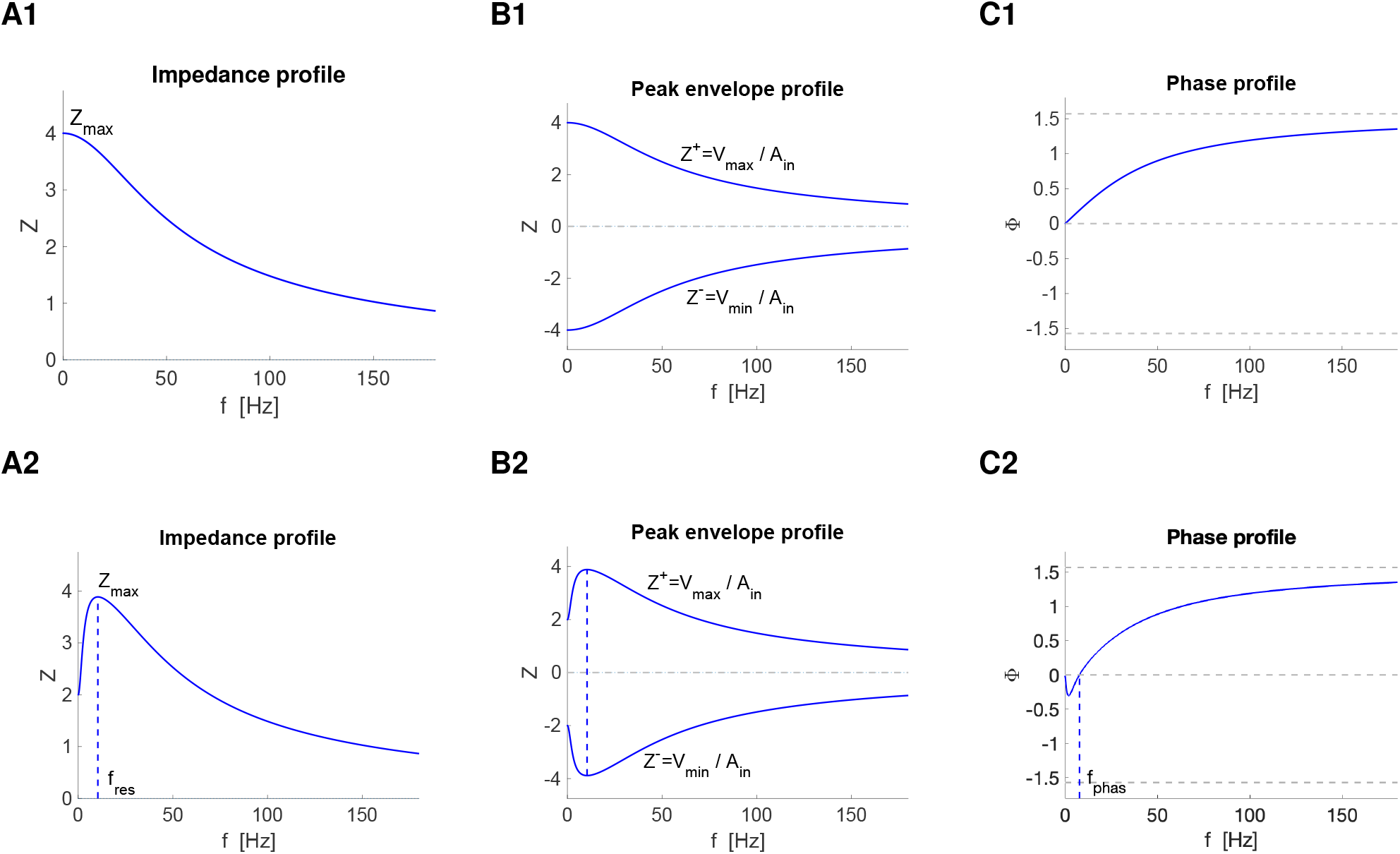
Schematic diagrams of the impedance (A), peak envelope (B) and phase (C) profiles for a passive cell (low-pass filter, LPF, and lagging response; top) and a cell exhibiting resonance (band-pass filter; BPF) and phasonance (leading-lagging response) (bottom). **A. A1.** *Z*_*max*_ = *Z*(0). **A2.** Resonance refers to the ability of a cell to exhibit peak in *Z*(*f*) at a non-zero (resonant) frequency *f*_*res*_. **B1 - B2** *V*_*max*_/*V*_*min*_ are the steady state peak /trough (maximum/minimum) voltage response values to the oscillatory inputs. *Z*^+^ and *Z*^−^ are the upper and lower *Z*-envelope profiles, respectively. **C1.** Phasonance refers to the ability of a cell to exhibit a zero phase (phase-shift, Φ = 0) response at a non-zero (phasonant) frequency *f*_*phas*_. The voltage response lags for *f > f*_*phas*_ and leads for *f < f*_*phas*_.

Electrical synapses mediated by gap junctions are ubiquitous in the nervous system and coexist with chemical (excitatory and inhibitory) synapses [22–34]. The metric usually used to characterize the strength of electrical synapses is the coupling coefficient (CC) *K* = Δ*V*_2_*/*Δ*V*_1_ [23–26, 28, 35–44, 44]. It is measured by injecting a step current into one cell and computing the ratio of the steady state voltage deflections of the indirectly (Δ*V*_2_; post-J) and directly (Δ*V*_1_; pre-J) stimulated cells. The larger *K*, the stronger the degree to which the pre-J and post-J cells are electrically coupled. Note that in this context (Fig. 2), we used the terms pre-J and post-J cells although electrical transmission mediated by gap junctions is reciprocal [40].

**Figure 2:**
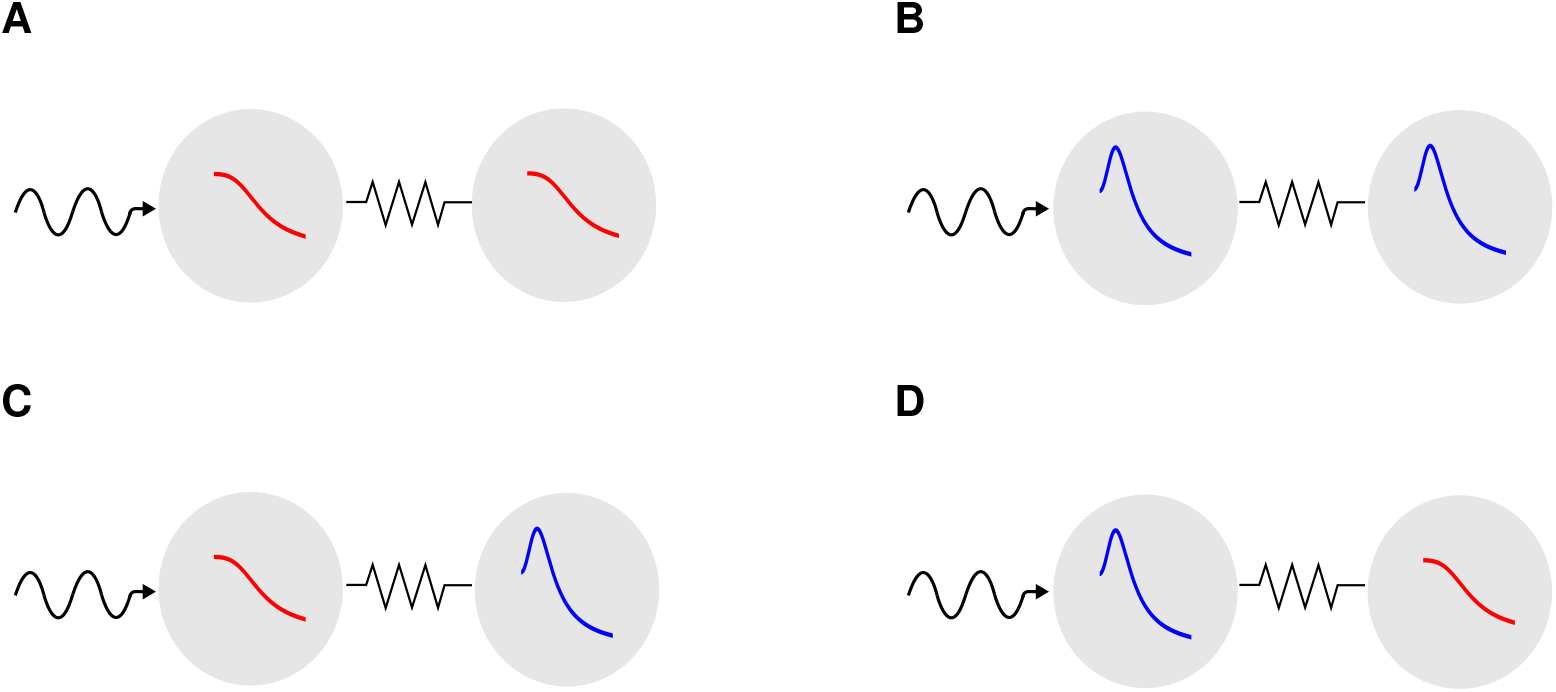
Schematic diagrams of electrically coupled cells receiving oscillatory inputs. The individual cells (disconnected) can have one-dimensional (1D, red) or two-dimensional (2D, blue) dynamics and exhibit either a low-pass filter (LPF; e.g., passive cells, red, see Fig. 1-A1) or a band-pass filter (BPF; e.g., resonators, blue, see Fig. 1-A2). The oscillatory inputs arrives to only one of the cells and are propagated to the second cell. We refer to the cell that receives the input as the pre-J cell and to the other cell as the post-J cell. In our models we use the indices *k* = 1 (pre-J) and *k* = 2 (post-J) cell. The same diagrams represent two-compartment models receiving oscillatory input to only one compartment. The filtering properties of the two connected cells may qualitatively differ from these of the individual (disconnected cells).

The magnitude of the CC is determined by the electrical connectivity and the cellular properties and can be altered by neuromodulators and additional synaptic inputs that affect the components of the CC [43, 45]. To the linear level of description, *K* varies between 0 to 1, increases monotonically with the gap junction conductance (*g*_*c*_) and de-creases monotonically with the post-J membrane (*g*_*L*_) and ionic conductances (*g*_2_) (e.g., in passive cells supplemented with a positive or negative feedback terms). These dependences occur in a balanced, homeostatic manner on the combined parameters *g*_*L*_*/g*_*c*_ and *g*_2_*/g*_*c*_. This characterization determines the efficiency of electrical transmission for heterogeneous network and establishes a path in the flow of information in gap junction connected networks. Electrical transmission is more efficient from the higher to the lower conductances as compared to the reverse direction.

The analysis of more realistic scenarios involving time-dependent inputs and the effects of the intrinsic cellular time constants requires the extending the notion of CC to include the response of gap junction networks (electrically coupled networks mediated by gap junctions) to oscillatory inputs [24,32,40,43]. This leads to the frequency-dependent (amplitude) coupling and phase-difference coefficient profiles defined as *K*(*f)* = *A*_2_(*f)/A*_1_(*f)* and the ΔΦ(*f)* = Φ_2_(*f)* − Φ_1_(*f)*, respectively, where *A*_1*/*2_(*f)* and Φ_1*/*2_(*f)* are the amplitude and phase profiles of the pre-J and post-J cells (described in detail in Section 2.6.1). For more complex signals, this definition can be further extended to be the quotient of the (complex) Fourier transforms of the post-J and pre-J responses. For passive cells, the *K*(*f)* profile are LPFs [23–29, 36, 40, 41, 46–50], indicating that the signal transmission is more efficient and the coupling is stronger for the slower than for the faster membrane potential fluctuations. The ΔΦ profile are positive and monotonically increasing. These two profiles depend on a combination of the intrinsic cellular properties and the gap junction connectivity.

Theoretical studies have mainly focused on models based on linear circuit theory. For linear systems, *K*(*f)* and ΔΦ(*f)* depend on the biophysical properties of the post-J cell and the gap junction conductance, and are independent of the biophysical properties of the pre-J and the input amplitude [43] (revisited in Section 3.1.1). Furthermore, modeling studies have largely used passive cell models (but see [32, 43, 49, 51–53]). Being linear and one-dimensional, the *K*(*f)* and ΔΦ(*f)* profiles for electrically coupled passive cells are a modulated version of the corresponding response profiles for the post-J cell and therefore remain LPFs(*K*) and positive (ΔΦ), respectively.

However, neurons possess a variety of ionic currents that are inherently nonlinear and their dynamics involve a multiplicity of time scales. It is largely unknown how the *K*(*f)* and ΔΦ(*f)* profiles are shaped by the participating building blocks when the above mentioned assumptions are relaxed (but see [43]). It is unclear how the properties of the pre-J cell and the input amplitude affect the *K*(*f)* and ΔΦ(*f)* profiles (in addition to the properties of the post-J cell and the gap junction conductance) when the neurons operate away from the linear regime.

The generation of cellular resonance and phasonance in individual cells requires the interplay of negative and positive feedback effects mediated by the gating variables associated to the intrinsic slow resonant currents (e.g., hyperpolarization-activated mixed-cation *I*_*h*_, M-type slow potassium *I*_*M*_ and T-type calcium *I*_*CaT*_ inactivation) and fast amplifying currents (e.g., persistent sodium *I*_*Nap*_ and *I*_*Ca,T*_ activation), which oppose and favor changes in voltage, respectively [1, 20, 21]. The presence of amplifying and resonant ionic currents regulate the frequency-dependent coupling by increasing the *K*(*f)* profile [32, 49, 51] or by causing *K*(*f*) profile to be a BPF [32, 52], indicating the existence of an intermediate frequency band at which the transmission of information is more efficient than for other frequencies. However, a systematic study of how presence cellular resonance and phasonance in the participating cells shape the *K* and ΔΦ profiles is lacking.

Multicompartment models of spatially extended neurons have been used to investigate the neuronal resonant properties distributed along the somato-dendritic axis and how the neuronal intrinsic ionic currents affect the response to, and propagation of spatially localized inputs [5, 54–62]. In an intriguing set of results [54, 55], it was experimentally demonstrated that two different types of subthreshold theta resonances [54, 55, 63–65] coexist along the somato-dendritic axis n hippocampal CA1 pyramidal cells (*in vitro*): (i) a perisomatic resonance generated by a slow-potassium M-type current (*I*_*M*_) and amplified by a persistent sodium current (*I*_*Nap*_), and (ii) a dendritic resonance generated by a hyperpolarization-activated mixed-cation (sodium and potassium) current or h-current (*I*_*h*_). The mechanisms that control the interaction between these two types of resonances are not well understood.

We address these issues in this paper. We consider a network of two electrically connected neurons mediated by gap junctions receiving external inputs (Fig. 2). This is the minimal network architecture that allows for a systematic study of the biophysical and dynamic mechanisms that shape the *K*(*f)* and ΔΦ(*f)* profiles. The participating neurons are either passive cells (LPFs; 1D; Fig. 1-A1) or resonators (BPFs; 2D; Fig. 1-A2) and exhibit positive (Fig. 1-C1) or mixed negative-positive (Fig. 1-C2; phasonators) phase responses. We begin our study by using linear models to understand how the participating building blocks (e.g., conductances, negative and positive feedback effects) operating at the single node level control the *K*(*f)* and ΔΦ(*f)* profiles. We then turn our attention to linear models as linearized models of nonlinear of neurons with biophysically realistic ionic currents and investigate (to the linear level of approximation) how the presence of cellular ionic currents and cellular resonance and phasonance contribute to shaping the *K*(*f)* and ΔΦ(*f)* profiles. We use *I*_*Nap*_, *I*_*h*_ and *I*_*Ks*_ motivated by the experimental results mentioned above and previous studies [21, 66, 67]. We then extend our investigation to include the model nonlinearities to understand how the *K*(*f)* and ΔΦ(*f)* profiles are shaped by the interaction between the pre-J and post-J cells, and not just the post-J cells as for linear models. For gap junction connected networks, the connectivity is typically symmetric, while for compartmental models is asymmetric as the result of the different geometries of the participating compartments [68, 69]. We investigate the effects of asymmetric connectivity to understand how the compartmental geometry interacts with the neuronal biophysical properties to shape the *K*(*f)* and ΔΦ(*f)* profiles.

The formalism and tools we develop and use in this paper can be extended to larger networks with an arbitrary number of nodes, to spatially extended multicompartment neuronal models and to models having ionic currents with regenerative and restorative processes such as the (*I*_*Ca,T*_) activation and inactivation.

Our results make experimentally testable predictions and have implications for the understanding of spike transmission, synchronic firing and coincidence detection in electrically coupled networks in the presence of oscillatory inputs.

Table 1 summarizes the notation used throughout the paper.

**Table 1:**
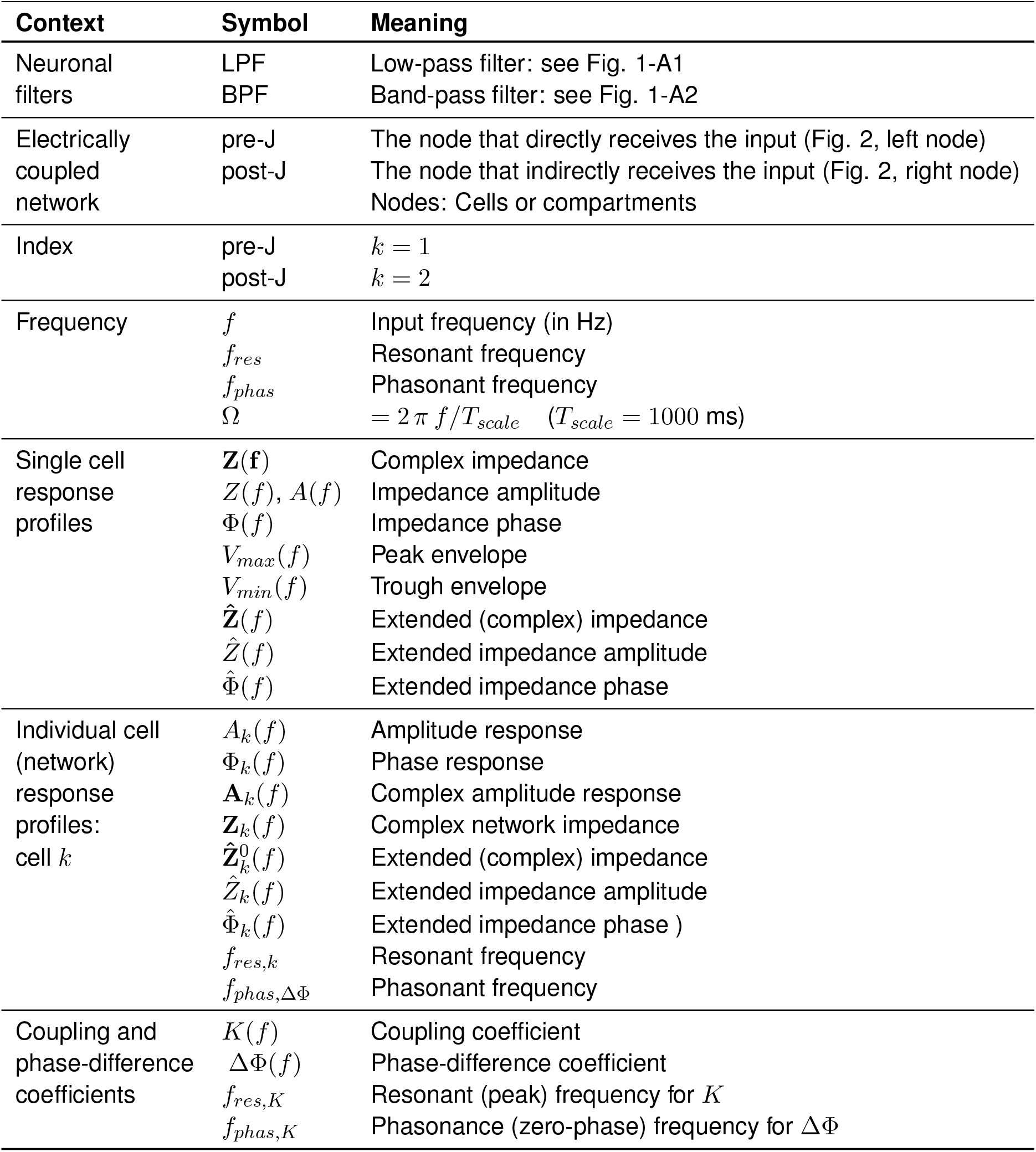
Glossary.

## 2 Methods

We consider a network of two electrically connected neurons mediated by gap junctions receiving external inputs (Fig. 2). With minimal modifications, the formalism we use describes two-compartment neurons receiving external inputs. The dynamics of the individual neurons are described by reduced, linearized [20, 21] or quadratized [70–72] biophysical (conductance-based) models of Hodgkin-Huxley (HH) type [69, 73]. The models include non-spiking ionic currents such as the persistent sodium current (*I*_*Nap*_), the hyperpolarization-activated mixed-cation (*I*_*h*_) current, the M-type slow-potassium current (*I*_*Ks*_). *I*_*Nap*_ is regenerative (amplifying, favors changes in voltage, provides positive feedback effects) and has fast dynamics, and it is assumed to be slaved to voltage, while *I*_*h*_ and *I*_*Ks*_ are restorative (resonant, oppose changes in voltage, provide negative feedback effects and have slower dynamics [66, 67, 71, 74, 75].

We refer the reader to S1 (Supplementary Material) for a description of the HH models and for the details on the linearization and quadratization processes as well as for the formulas linking the biophysical parameters with the parameters of the reduced (linearized or quadratized) models.

We use the following units for all the models in this paper: mV for voltage, ms for time, *µ*F/cm^2^ for capacitance, *µ*A/cm^2^ for current, mS/cm^2^ for the maximal conductances and Hz for frequency.

### 2.1 Networks of electrically coupled linearized cells

The linearized network model is described by (see Section S1.3)

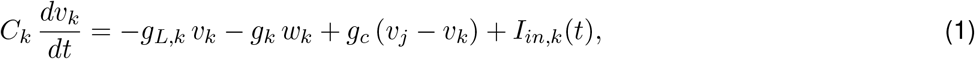

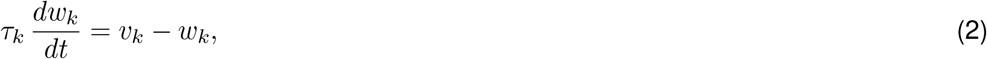

for *k, j* = 1, 2. In eqs. (1)-(2), *t* is time, *v*_*k*_ represent the membrane potential, *w*_*k*_ represent the gating variable, *C*_*k*_ are the capacitances, *g*_*L,k*_ are the linearized leak maximal conductances, *g*_*k*_ are the ionic current linearized conductances, *τ*_*k*_ are the linearized time constants for the linearized gating variables *w*_*k*_, *g*_*c*_ is the gap junction coupling coefficient, and *I*_*in,k*_(*t*) are time-dependent input currents. The linearized component of the ionic currents with instantaneously fast dynamics (e.g., *I*_*Nap*_) are incorporated in *g*_*L,k*_. For two-compartment models with different connectivity coefficients, we use the notation *g*_*c*_ = *g*_*c,kj*_.

Note that the heterogeneity due to different values of the biophysical parameters in the original conductance-based model is translated into the linearized model both explicitly and implicitly through the equilibria (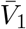 and 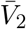) of the original system. Unless stated otherwise, we use *C*_1_ = *C*_2_ = 1 for all models in this paper.

### 2.2 Networks of electrically coupled quadratized cells

The quadratized network model is given by (see Section S1.4)

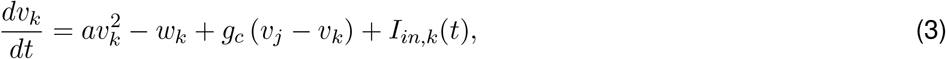

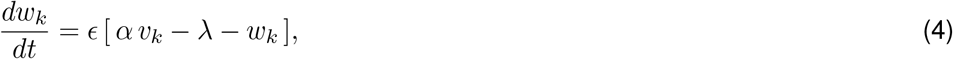

for *k, j* = 1, 2, *k* ≠ *j*, where *a >* 0 captures the effects of an amplifying current, *g*_*c*_ represents the gap junction coupling coefficient, *ϵ* is the inverse of the time constant of the resonant gating variable, *α* ≥ 0 represents the strength of the negative feedback (resonant) current, and *λ* is a combination of parameters including, and having the same sign as *I*_*app*_ in the original biophysical model. Increasing (decreasing) values of *I*_*app*_ cause *λ* to increase (decrease) and displaces the *w*-nullcline to the right (left), thus favoring (opposing) the generation of subthreshold oscillations and spikes (Fig. 3). Geometrically, *a* controles the curvature of the parabolic *v*-nullcline, *α* controls the slope of the *w*-nullcline, *ϵ* represents the time scale separation between the two variables, and *λ* controls the displacement of the *w*-nullcline with respect to the *v*-nullcline. As for the linear models, for two-compartment models with different connectivity coefficients, we use the notation *g*_*c*_ = *g*_*c,kj*_.

**Figure 3:**
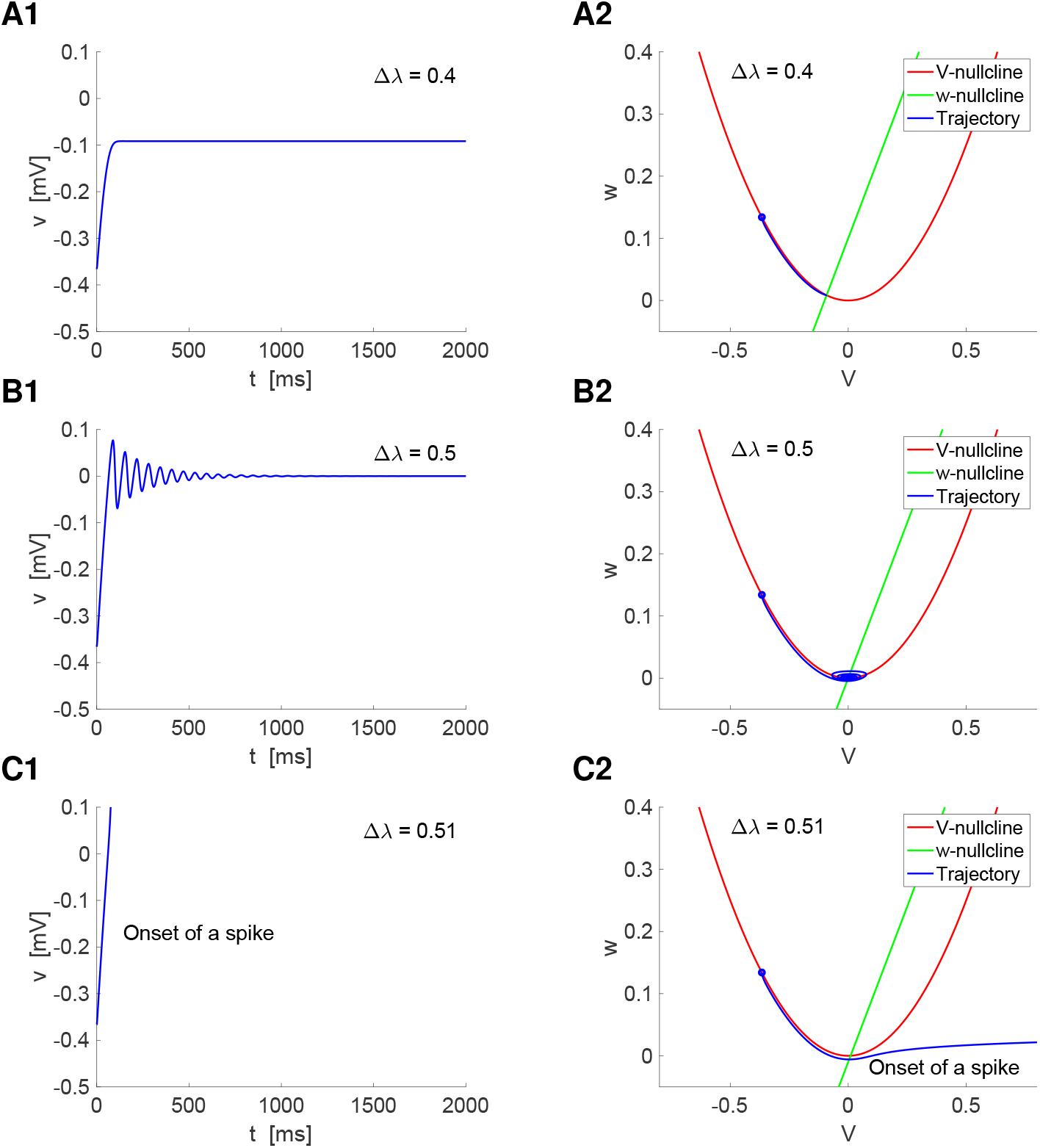
Response of a single cell quadratic model to constant inputs: Voltage traces and phase-plane diagrams. We used the model (3)-(4) for a single cell (*g*_*c*_ = 0) with *a* = 1, *α* = 1 and *ϵ* = 0.01. The initial conditions (blue dots) were determined as the steady-state response to a baseline value of *λ*: *λ*_*base*_ = *−*0.5 and the values of *λ* were determined as *λ* = *λ*_*base*_ + Δ*λ*. **Left Column.** Voltage traces. **Right column.** Phase-plane diagrams. The *V* -nullclines (red) are the curves satisfying *w* = *a v*^2^ and the *w*-nullclines (green) are the lines satisfying *w* = *α v − λ*. The fixed-point is determined by the interSection between the two nullclines. The trajectories (blue) are initially located at the blue dot and evolve towards the fixed-point. **A.** Δ*λ* = 0.4 (*λ* = *−*0.1). The model exhibits an almost monotonic increase to the steady state value of *v* (there is an imperceptible overshoot at low values of *t*). **B.** Δ*λ* = 0.5 (*λ* = 0). The model exhibits subthreshold oscillations around the steady-state value of *v*. **C.** Δ*λ* = 0.51 (*λ* = 0.1). The model produces the onset of spikes when the trajectory increases unboundedly and escapes the subthreshold voltage regime. The model ceases to be a good approximation to any biophysical model. A voltage reset mechanism is needed to bring the trajectory back to the subthreshold regime.

Nonlinearities of parabolic type in the subthreshold voltage regime are generated in the presence of regenerative/amplifying currents (e.g., *I*_*Nap*_). The resulting voltage nullclines of parabolic type are shaped by the interplay of these and the other participating ionic currents (e.g., *I*_*h*_, *I*_*Ks*_).

Quadratization of biophysically plausible models of HH type extends the notion of linearization to include the parabolic-like properties of the *V* -nullcline in the subthreshold regime and therefore capture more realistic aspects of the dynamics of these models, which can be missed by the corresponding linearization [70–72, 74, 76, 77]. In contrast to linearization, which is carried out around the model’s fixed point (determined by the interSection of the model’s nullclines), quadratization is done around the extremum (minimum or maximum) of the *V* -nullcline of quadratic type and the resulting quadratized nullcline is always concave up.

The process of quadratization [70, 71] consists of expanding the right-side of the (original) biophysical model’s differential equations into Taylor series around the minimum/maximum (*V*_*e*_, *x*_1,*e*_) of the parabolic-like *V* -nullcline in the subthreshold regime, neglecting all the terms with power bigger than two in the equation for *V* and bigger than one in the equation for *x*_1_, and translating the minimum/maximum of the *V* -nullcline to the origin. The description of the quadratization process as well as the definition of the quadratized parameters below in terms of the parameters of biophysically realistic (conductance-based) models of Hodgkin-Huxley type are presented in the Supplementary material S1.4. Additional details are provided in [70–72] for 2D and 3D models. A Comparison between the original (biophysical) *I*_*Nap*_ + *I*_*h*_ and *I*_*Nap*_ + *I*_*Ks*_ models (see below) and their quadratization is shown in Fig. S6-A3 and -B3 [72]. A detailed description of the process for 3D models of HH type including time-dependent external currents and synaptic inputs is presented in [70] and also discussed in [72] in the context of reduced neuronal models.

While the assumption that the *V* -nullcline is parabolic-like in the subthreshold regime (e.g., Fig. 2 in [70] and Fig. 7 in [72]) is a rather general property of the type of models of HH described above, in certain, also rather general parameter regimes, these models can have *V* -nullclines of cubic-type [66, 67] around which (not around the parabolic component of the cubic-like nullcline) relevant subthreshold behavior such as subthreshold oscillations are generated.

### 2.3 The I_Nap_ + I_h_ and I_Nap_ + I_Ks_ conductance-based models

The *I*_*Nap*_ + *I*_*h*_ and *I*_*Nap*_ + *I*_*Ks*_ models [66, 71] are conductance-based models of Hodgkin-Huxley type [73] in the subthreshold voltage regime that involve the interaction of the two active ionic currents that define them. Both models include two variables: the voltage *V* and a gating variable associated to the slower *I*_*h*_ or *I*_*Ks*_ currents. The dynamics of *I*_*Nap*_ is assumed to be instantaneously fast (slaved to voltage). The mathematical formulation is presented in Section S1.1 together with the parameter values we use in this paper. The two models are adapted versions of the models presented in [75, 78] (see also [66, 67, 70–72]).

Both the *I*_*Nap*_ + *I*_*h*_ and *I*_*Nap*_ + *I*_*Ks*_ models involve the interaction of a restorative current, *I*_*h*_ and *I*_*Ks*_, having a resonant gating variable, and the same regenerative current, *I*_*Nap*_, having an amplifying gating variable. Because all these currents have a single gating variable, we refer to them as resonant and amplifying currents, correspondingly. They provide a positive (*I*_*Nap*_) and slower negative (*I*_*h*_ and *I*_*Ks*_) feedback effects, favoring and opposing changes in voltage, respectively.

The two models have *V* -nullclines of quadratic type or are quasi-linear in the subthreshold regime for the parameter sets considered in this paper (Figs. S4, and S6-A1 and -B1) (see also [70], Fig. 2, and [72], Fig. 7). Note that while the two models describe the interplay of a resonant (*I*_*h*_ or *I*_*Ks*_) and an amplifying current (*I*_*Nap*_), the nullclines are qualitatively a mirror image of each other: the *V* -nullcline is concave down for the *I*_*Nap*_ + *I*_*h*_ model (Fig. S4) and concave up for the *I*_*Nap*_ + *I*_*Ks*_ model (Figs. S1.2.4). The process of quadratization (see Section S1.4) uncovers the qualitative similarity of the phase-plane diagrams for both models (Fig. S6) (except for the concavity differences between the nullclines for the slow gating variable).

Figs. S4 and S1.2.4 illustrate the dynamics of the *I*_*Nap*_ + *I*_*h*_ and *I*_*Nap*_ + *I*_*Ks*_ models, respectively. In the subthreshold regime, the models have a stable fixed-point, towards which trajectories evolve, and sometimes an additional, unstable fixed-point. The system transitions from quasi-linear (in a vicinity of the stable fixed-point) to parabolic-like as some control parameter (e.g, *I*_*app*_, *G*_*p*_, *G*_*h*_) and the dynamics are modified accordingly. Increasing values of certain parameters (e.g., *I*_*app*_, *G*_*h*_, *G*_*p*_) cause a Hopf bifurcation near the knee of the *V* -nullcline where the fixed-point looses stability and moves to the “right branch” of the parabolic-like nullcline. Trajectories leaving the subthreshold regime is interpreted as the onset of a spike. When the models are supplemented with a return mechanism to the subthreshold regime, they are referred to as models of quadratic integrate-and-fire type with adaptation [72, 74, 76].

The *I*_*Nap*_ model is a 1D reduced version of the 2D models discussed above, which does not include any slow ionic current. It is described in Section S1.2 in the Supplementary Material by eqs. (S1) with *I*_1_ = 0 and *I*_2_ = *I*_*Nap*_ (eq. S3). Analogously to the other two models, it has parabolic-like or quasi-linear speed (*dV/dt* vs. *V)* curve in the subthreshold regime. The dynamics of the *I*_*Nap*_ model are illustrated in Fig. S2 in the Supplementary Material. For low enough values of *G*_*p*_, the model develops nonlinearities of parabolic type, but remains quasi-linear in a vicinity of the stable fixed-point (blue dot at the interSection of the speed curve). The trajectories (not shown) move along the *V* axis and converge to these fixed points. As *G*_*p*_ increases, the nonlinearities become more pronounced in a vicinity of the stable fixed-point, For the saddle-node bifurcation value of *G*_*p*_, the fixed-points cease to exist and trajectories leave the subthreshold regime. This is interpreted as the onset of a spike and if the model is endowed with a return mechanism to the subthreshold regime, it becomes a model of quadratic integrate-and-fire type [72] (the well-known quadratic integrate-and-fire model [79, 80] has a purely parabolic speed curve)

### 2.4 Single cell voltage response to oscillatory inputs: low- and band-pass filters

The voltage response of a neuron receiving an oscillatory input current (with frequency *f)* is typically measured in terms of the impedance **Z**(*f)*, a complex quantity with amplitude *Z*(*f)* = *|***Z**(*f)|* and phase Φ(*f)*. Following others, we refer to *Z*(*f)* simply as the impedance. Passive cells are low-pass filters (LPFs; e.g., Figs. 1-A1) and exhibit a lagged response (e.g., Figs. 1-C1). More complex cells can be band-pass filters (BPFs; e.g., Figs. 1-A2) and exhibit a lagged/leading response (e.g., Figs. 1-C2). A system exhibits *resonance* if *Z*(*f)* peaks at a non-zero (resonant) frequency *f*_*res*_ (e.g., Figs. 1-A2) and *phasonance* if Φ(*f)* vanishes at a non-zero (phasonant) frequency *f*_*phas*_ (e.g., Figs. 1-C2).

The generation of cellular resonance requires the interplay of negative and positive feedback effects mediated by the gating variables associated to the intrinsic, resonant (e.g., *I*_*h*_, *I*_*Ks*_) and amplifying (*I*_*Nap*_) ionic currents, respectively.

The resonant properties of 2D linear and nonlinear neuronal systems, including their relationship between the cellular intrinsic biophysical properties and the dynamic mechanism of generation of resonance have been investigated extensively by us and other authors [20, 21, 71, 81]. In the Supplementary Material S2 we present the analytical expressions of *Z*(*f)* and Φ(*f)* for linear (1D and 2D) neuronal models receiving a sinusoidal input current of the form

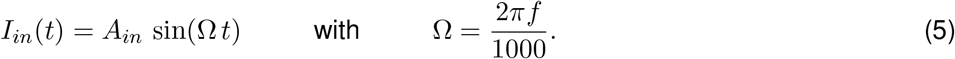

For nonlinear systems we define the peak envelope impedance profile (Fig. 1-B) [81, 82] as

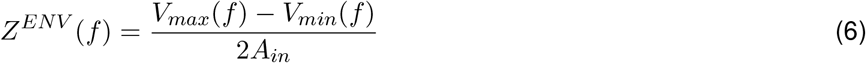

where *V*_*max*_(*f*) and *V*_*min*_(*f*) are the peak and trough envelope profiles, respectively, defined as the maximum and minimum values, respectively, of the steady-state oscillatory voltage *V*_*out*_(*f*) as a function of the input frequency *f*. For linear systems, *Z*^*ENV*^ (*f*) = *Z*(*f*). Similarly, the phase Φ is computed as the distance between the peaks of the output and closest input normalized by period, and *f*_*phas*_ is the non-zero frequency where Φ vanishes. These quantities extend the concept of impedance and phase to nonlinear systems under certain conditions (the input and output frequencies coincide and the output amplitude is uniform across cycles for a given input with constant amplitude).

### 2.5 Network voltage response to oscillatory inputs

We now consider a two-cell network (Fig. 2) receiving sinusoidal inputs of the form (5) *I*_*in,k*_(*t*) = *A*_*in,k*_ sin(Ω *t*) (*k* = 1, 2). In this paper, we consider inputs to only one cell (cell 1); i.e., *I*_*in,1*_(*t*) = *A*_*in*_ sin(Ω *t*) and *I*_*in,2*_(*t*) = 0. For generality, in the Supplementary Material S3 we provide the calculations using linear networks of *N* cells, each having 2D linear dynamics and sinusoidal inputs to all cells in the network.

#### 2.5.1 Networks of linear nodes

For networks having linear nodes and linear connectivity, the voltage response of each cell has the form *V*_*out,k*_(*t*; *f*) = *A*_*k*_(*f*) sin(Ω *t −* Φ_*k*_(*f*)). The amplitude *A*_*k*_(Ω) and phase Φ_*k*_(Ω) are the component of the complex network voltage amplitude response **A**_*k*_(*f*) derived in Section S3

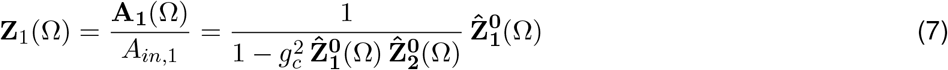

and

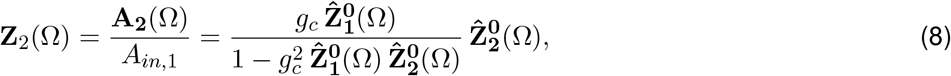

where 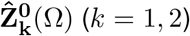 are the extended (complex) impedances of the individual (disconnected) nodes in the network obtained from eq. (S32) in the Supplementary Material S2 by substituting the parameters *g*_*L,k*_ by *g*_*L,k*_ + *g*_*c*_

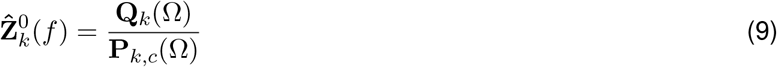

where

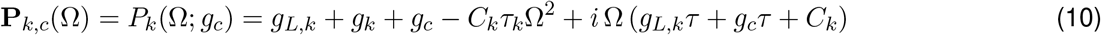

and

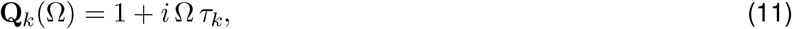

for *k* = 1, 2. This captures the fact that the gap junction current includes a “self-connectivity” term, which is incorporated into the autonomous part of the corresponding nodes *k*. The extended amplitude impedance and phase profiles are given by eqs. (S53)-(S54) in Section S3.1.

#### 2.5.2 Networks of nonlinear nodes: network impedance and peak envelope profiles

For nonlinear networks we adapt the tools defined in the previous Section by adding the appropriate indices and referring to the corresponding quantities for the individual (disconnected) cells by adding a superscript “0”. In contrast to linear systems, the voltage response for nonlinear systems is not symmetric. Therefore, while the impedance amplitude captures the frequency content of the voltage response signal, it does not necessarily describe the frequency-dependent prope rties of the peak and trough envelope profiles. Because neuronal signal occurs via voltage threshold-like mechanisms, we will look at the peak envelope profiles *V*_*max,k*_(*f*) for *k* = 1, 2

### 2.6 Coupling coefficient: Frequency-dependent profiles and preferred communication frequencies

#### 2.6.1 Amplitude and phase

We extend the notion of the (steady state) coupling coefficient in response to constant (DC) inputs [23–25, 28, 35] to include the frequency-dependent effects in response to oscillatory inputs by defining the amplitude coupling coefficient

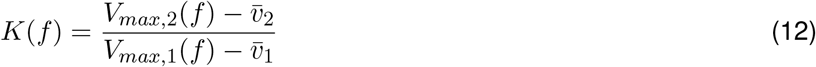

where 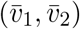 is the fixed point of system (1)-(2), when unperturbed and the phase-difference coupling coefficient

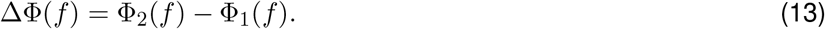

For linear systems 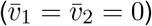, *K*(*f)* and ΔΦ(*f)* are the components of the following complex quantity

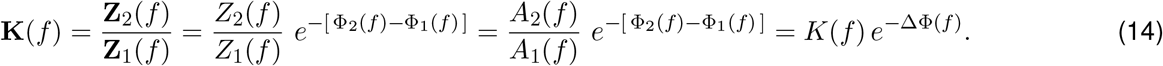

From (7)-(8),

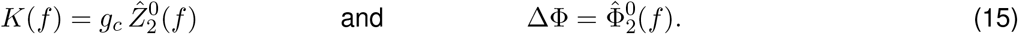

#### 2.6.2 *K*-resonance and ΔΦ-phasonance

We define *K*-resonance as the ability of the coupling coefficient *K*(*f*) to exhibit a peak at a non-zero input frequency *f*_*res,K*_, and ΔΦ-phasonance as the ability of the frequency-dependent coupling phase ΔΦ(*f*) to vanish for a nonzero input frequency *f*_*phas,ΔΦ*_.

From eq. (12), a necessary condition for the existence of *K*-resonance (or *K*-antiresonance) is given by

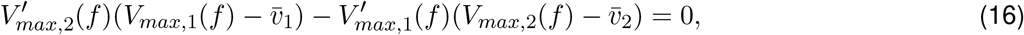

for some value *f >* 0.

We characterize the *K*(*f*) profile by using two attributes: (i) the resonant frequency *f*_*res,K*_, defined as the peak frequency of *K*(*f*), and (ii) the resonance amplitude *Q*_*K*_ = *K*_*max*_ − *K*(0). If the *K* profile does not exhibit resonance, then *f*_*res,K*_ = *Q*_*K*_ = 0. We characterize the ΔΦ(*f*) profile by using the phasonant frequency *f*_*phas,ΔΦ*_.

### 2.7 Electrically coupled neuronal compartments

The mathematical formulation of two-compartment models of spatially extended cells (e.g., dendritic, somato-dendritic) having linear dynamics is as described in Sections 2.1 and 2.2 with *g*_*c*_ substituted by *g*_*c*_/σ_*k*_ (*k* = 1, 2) with *σ*_1_ + *σ*_2_ = 1, where *σ*_*k*_ represents the fraction of the total cell area taken up by the corresponding compartment [83–85]. For linearized models, because of the possible asymmetry between the two compartments, there is a constant membrane potential effect due to the differences in the resting potential of the two compartments.

For two-compartment models of linear nodes where the input arrives only to one of the compartments (compartment 1), the complex network voltage amplitude response presented in Section 2.5.1 extends to (see also Section S3)

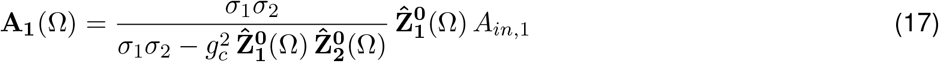

and

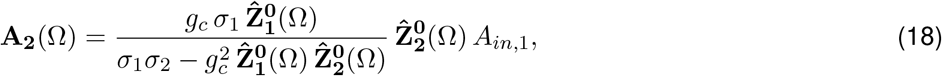

where 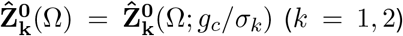 are the extended (complex) impedances of the individual (disconnected) nodes in the network obtained from eq. (S32) in the Supplementary Material S2 by substituting the parameters *g*_*L,k*_ by *g*_*L,k*_ + *g*_*c*_/σ_*k*_. This can be straightforwardly extended to more detailed formulations involving the geometric (length, area) and biophysical properties of neuronal compartments and a larger number of compartments [68, 69].

It follows that

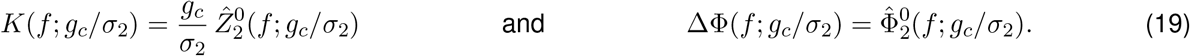

### 2.8 Numerical simulations

To compute the numerical solutions we used a Runge-Kutta method of order 4 [86] with a time step Δ*t* = 0.01 ms. Smaller values of Δ*t* have been used to check the accuracy of the results. All neural models and metrics, including phase-plane analysis, were implemented in MATLAB (The Mathworks, Natick, MA). The codes are available at https://github.com/BioDatanamics-Lab/Coupling_Coefficient-25_05.

## 3 Results

### 3.1 Steady-state coupling coefficient *K* in response to constant inputs

#### 3.1.1 Linear models: revisited

The steady-state response of the electrically coupled linear cells (1)-(2) to a constant (DC) inputs *I*_1_ (*I*_2_ = 0) is given by

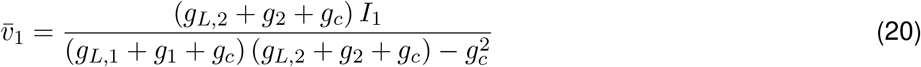

and

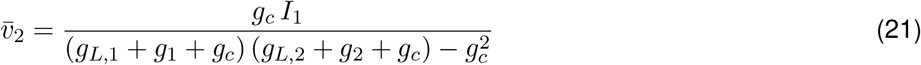

for the pre-J and post-J cells, respectively.

The coupling coefficient is therefore given by

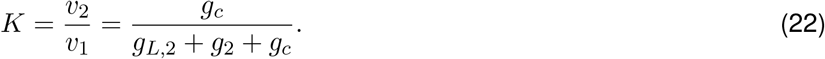

As reported before for networks of passive cells (*g*_2_ = 0) [43, 44], *K* is independent of the input current and the intrinsic properties of the pre-J cell, and varies between 0 and 1. For networks of 2D linear cells, *K* is affected by *g*_2_ in addition to*g*_*L,2*_ and it remains independent of the intrinsic properties of the pre-J cell.

Note that if the connectivity coefficients are different, as it is usually the case for two-compartment models, then

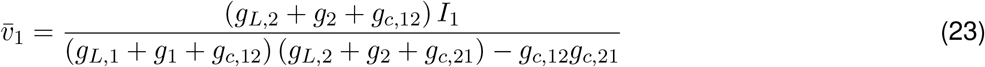

and

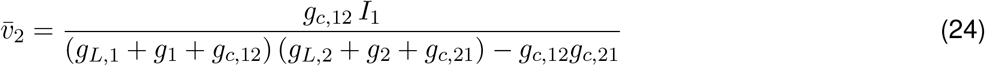

for the pre-J and post-J cells, respectively, and

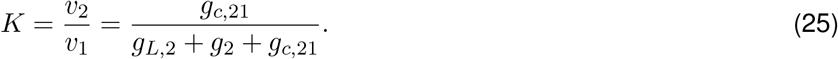

#### 3.1.2 Quadratic models

Our analytical results for networks of weakly nonlinear cells (Section S4) show that as the result of the presence of nonlinearities, *K* is no longer independent of *I*_1_ and is (weakly) affected by the intrinsic properties of both the pre- and post-J cells (Section S4.1). Here, we test these ideas for the quadratic model (3)-(4).

The dynamics of the quadratic model is illustrated in Fig. 3. Changes in the values of the constant (DC) input current are monotonically reflected in the parameter *λ*. In our simulations, we first determined a baseline value of *λ, λ*_*base*_ and computed the stationary voltage response *v*_*ss,base*_ of the individual cells to this value. We then computed *λ* as *λ* = *λ*_*base*_ + Δ*λ* and used *v*_*ss,base*_ as the initial condition in our simulations (blue dots in Fig. (right column).

Fig. 4-A illustrates the response of the pre-J (*v*_1_) and post-J (*v*_2_) cells for values of *g*_*c*_ that increase from panel A1 to panel A2. Fig. 4-B shows curves of *K* as a function of Δ*λ* for representative scenarios. In contrast to linear models, there is a dependence of *K* on *λ*, which is more pronounced for the larger values of *g*_*c*_. Comparison between panels B1 and B2 illustrate that the dependence of *K* on *λ* is independent on *ϵ*. Comparison between panels B1 and B3 illustrate that the dependence of *K* on *λ* is more pronounced and the values of *K* are larger the smaller *α* (all other parameters fixed). Finally, comparison between panels B1 and B4 illustrate that the dependence of *K* on *λ* is slightly more pronounced and the values of *K* are larger the smaller *a* (all other parameters fixed).

**Figure 4:**
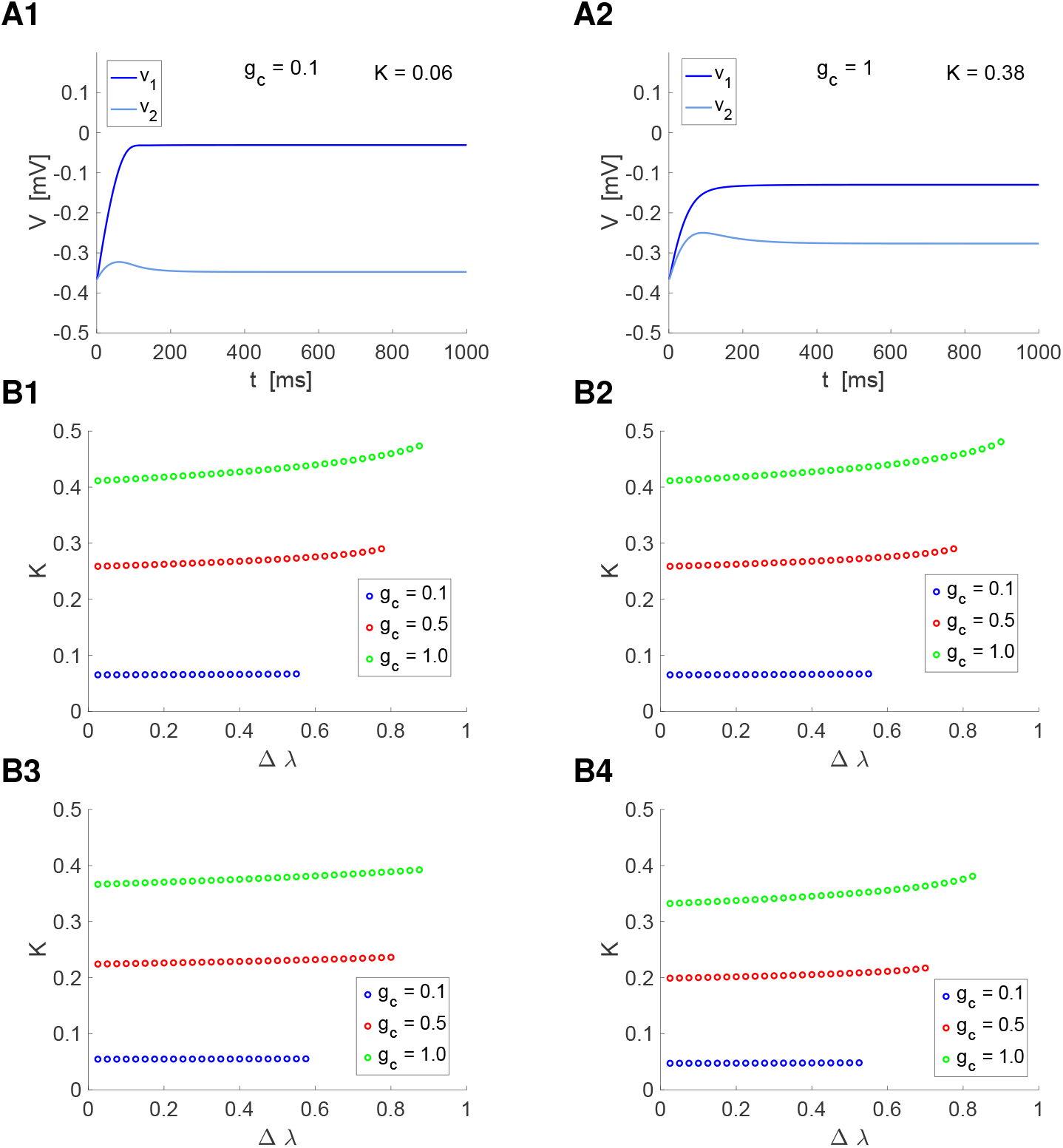
Response of a single cell quadratic model to constant inputs: Coupling coefficient. We used the model (3)-(4) with *a* = 1, *α* = 1 and *ϵ* = 0.01. The initial conditions were determined as the steady-state response to a baseline value of *λ*: *λ*_*base*_ = *−*0.5 and the values of *λ* were determined as *λ* = *λ*_*base*_ +Δ*λ*. **A.** Voltage traces for representative examples and *λ* = 0.5. **A1.** *g*_*c*_ = 0.1. **A2.** *g*_*c*_ = 1. **B.** Dependence of *K* on *λ* = *λ*_*base*_ + Δ*λ* for representative values of *g*_*c*_, *ϵ, α* and *a*. **B1.** *a* = 1, *α* = 0.25, *ϵ* = 0.01. **B2.** *a* = 1, *α* = 0.25, *ϵ* = 0.1. **B3.** *a* = 1, *α* = 1, *ϵ* = 0.01. **B4.** *a* = 2, *α* = 0.25, *ϵ* = 0.01.

### 3.2 Response to oscillatory inputs: Electrically coupled linear cells

Linear models of single neurons exhibit a variety of amplitude and phase profiles in response to oscillatory inputs (e.g., Fig. 1) [1,20,21,87–89] (see Section S2). Neuronal linearized models encode information about the inherently nonlinear biophysical properties of neurons and networks. Therefore, linearized models can be used to analyze the nonlinear dependence of the *K*(*f*) and ΔΦ(*f*) filtering properties on the resonant and amplifying mechanisms operating at the single cell level by tracking the changes in the (linearized) model parameters as the result of the changes in the levels of the interacting ionic currents. While the response for each set of parameter values is linear, the variation of the responses as the parameters are recalculated for each biophysical scenario capture nonlinear effects such as the nonlinear amplification that occurs near threshold.

Here we analyze the response of electrically coupled linear cells to oscillatory inputs. We use these results in the next Section to analyze the response of biophysical models to oscillatory inputs. The expressions for the linearized conductances *g*_*L*_ and *g* in terms of the biophysical parameters of the original (biophysical) model are presented in Section S1.3 by eqs. (S14) and (S15).

For purely linear networks (linear nodes and linear connectivity) of the form (1)-(2), the network complex amplitude response to oscillatory inputs for the pre-J (**A**_1_) and post-J (**A**_2_) cells are given by eqs. (7) and (8), respectively (see Section S3 for details on the formulas and their derivation). The numerators describe the response of the directly activated pre-J cell and the indirectly activated post-J cell, respectively, modulated by the presence of the gap junction via de extended impedances (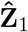 and 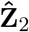). The denominators, describe the interaction between the two cells in response to the oscillatory input.

From eqs. (15), the CC *K*(*f*) for linear systems is proportional to the extended impedance of the post-J cell and the phase-difference coefficient ΔΦ(*f*) is equal to the extended phase of the post-J cell. Both quantities are independent of the input amplitude (*A*_*in*_) to the pre-J cell and of the intrinsic properties of the pre-J cell. While the formulas are valid for both cells, here we focus on *g*_*L,2*_ and *g*_2_ for the post-J cell since the *K*(*f*)- and ΔΦ(*f*)-profiles depend on these parameters.

#### 3.2.1 Passive cells

For two electrically coupled passive cells (1)-(2) (*g*_*k*_ = 0, *k* = 1, 2) (Fig. 2-A), the coupling and phase-difference coefficient profiles are given, respectively, by

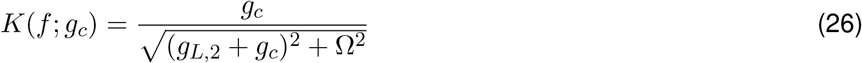

and

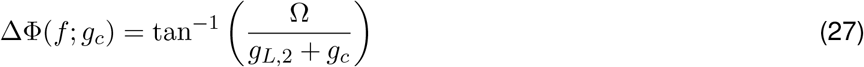

The calculations are presented in the Supplementary material S2, eqs. (S38) and (S39).

The *K*(*f*)-profiles are LPFs (Fig. 5, left) and the ΔΦ(*f*) profile are monotonically increasing (Fig. 5, right), indicating that the lag difference between the two cells is monotonically increasing. Therefore, the results from this Section remain true when the pre-J cells is a resonator (Fig. 2-D).

**Figure 5:**
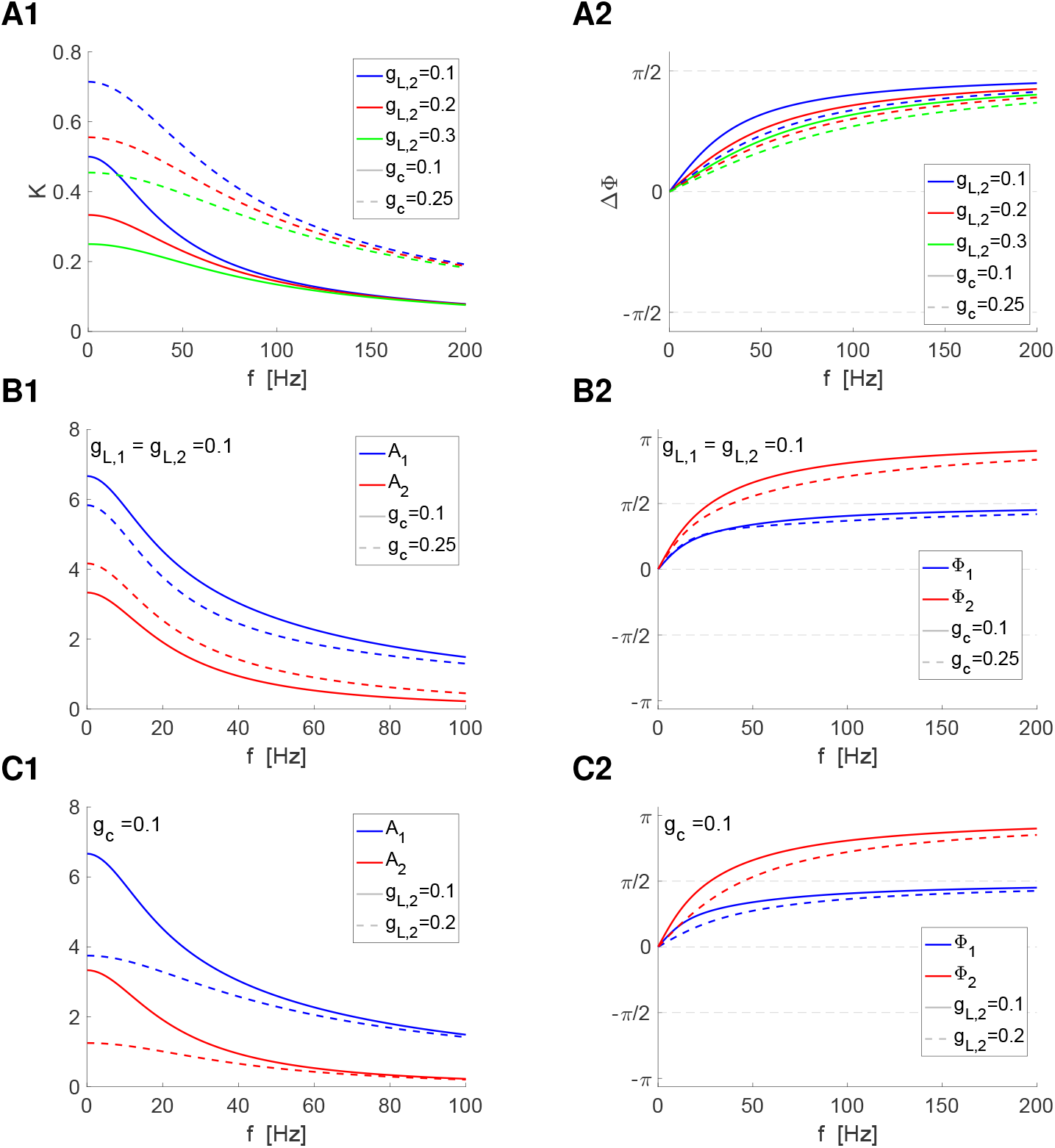
Response of two electrically coupled passive cells to oscillatory inputs for representative parameter values. The linear model is given by eqs. (1)-(2) with *g*_1_ = *g*_2_ = 0. Cells 1 and 2 are the pre-J and post-J cells, respectively. **A.** Coupling and phase-difference coefficients. **A1.** Coupling coefficient *K* given by eq. (26). **A2.** Phase-difference coefficient ΔΦ given by eq. (27). **B.** Amplitude and phase responses for the pre-J (blue) and post-J (red) cells for representative values of *g*_*L,1*_, *g*_*L,2*_ and *g*_*c*_. The blue curves and red curves correspond to the solid- and dashed-blue curves in panels A, respectively. These quantities were computed using eqs. (7)-(8) with *g*_1_ = *g*_2_ = 0. **B1.** Amplitude responses. **B2.** Phase responses. **C.** Amplitude and phase responses for the pre-J (blue) and post-J (red) cells for representative values of *g*_*L,1*_, *g*_*L,2*_ and *g*_*c*_. The blue curves and red curves correspond to the solid- and dashed-blue curves in panels A, respectively. These quantities were computed using eqs. (7)-(8) with *g*_1_ = *g*_2_ = 0. **C1.** Amplitude responses. **C2.** Phase responses.

Fig. 5-A illustrates the dependence of the *K*(*f*)- and ΔΦ(*f*)-profiles on *g*_*c*_ and *g*_*L,2*_. The network *A*-profiles (individual cells) are LPFs (Fig. 5-A1). Increasing values of *g*_*L,2*_ (for fixed values of *g*_*c*_) attenuate the *A*-profiles and make them shallower. Increasing values of *g*_*c*_ (for fixed values of *g*_*L,2*_) also make the *K*(*f)*-profiles shallower, but amplify them, as expected. In the first case, the recruitment of the post-J cell is stronger, which in turn weakens the response of the pre-J cell (Fig. 5-B1), while in the second case, the response of the two cells is weaker as *g*_*L,2*_ increases (Fig. 5-C1).

The network Φ-profiles are monotonically increasing (Fig. 5-A2). Increasing values of *g*_*L,2*_ (for fixed values of *g*_*c*_) and increasing values of *g*_*c*_ (for fixed values of *g*_*L,2*_) reduce the rate of increase of the ΔΦ(*f)* profiles, indicating that the response oscillations’ peaks for the two cells are closer the larger the input frequency. The stronger recruitment of the post-J cell as *g*_*c*_ increases (Fig. 5-B1) is accompanied by a reduction in both the network phases and phase-difference for each input frequency *f* (Fig. 5-B2). A similar effect is produced by increasing values of *g*_*L,2*_.

#### 3.2.2 Linear (2D) resonators: Interplay of positive and negative feedback effects

Resonators refer to cells that exhibit resonance [1]. For simplicity, here, we include in this category cells that exhibit resonance both in the absence or presence of damped subthreshold oscillations [20, 21] where the fixed-points are nodes and foci, respectively (i.e., we do not distinguish between resonators and damped oscillators). Resonators may also show phasonance [20, 21]. Resonance, phasonance and oscillations require the interplay of positive and slower negative feedback effects that favor and oppose changes of voltage, respectively. In the linearized models (1)-(2), the negative feedback is represented by *g*_*k*_ *>* 0 (*k* = 1, 2) and the positive feedback is embedded in the parameters *g*_*L,k*_.

For two electrically coupled 2D linear cells (1)-(2) (*g*_*k*_ *>* 0, *k* = 1, 2) (Fig. 2-B), the coupling and phase-difference coefficient profiles are given, respectively, by

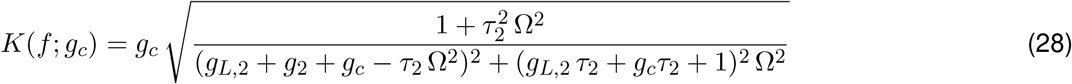

and

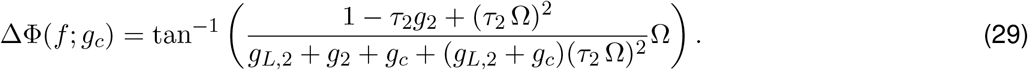

The calculations are presented in the Supplementary material S2, eqs. (S33) and (S34) with *C* = 1. These results in this Section remain true when the pre-J cells is passive (Fig. 2-C).

Fig. 6 illustrates the *K*(*f*) and ΔΦ(*f*) profiles for representative scenarios. We focus on individual cells that exhibit resonance (e.g., Fig. 6-D1, dashed-blue) and phasonance (e.g., Fig. 6-D2, dashed-blue). The *K*(*f*) profiles are BPFs(f (Fig. 6-A1 to -C1), exhibiting *K*-resonance, and the ΔΦ(*f*) profiles show ΔΦ-phasonance (e.g., Fig. 6-A2 to C2) in most cases considered. As mentioned above, these profiles are determined by the parameter of the post-J cell and *g*_*c*_, and are independent of the input amplitude and the properties of the post-J cell.

**Figure 6:**
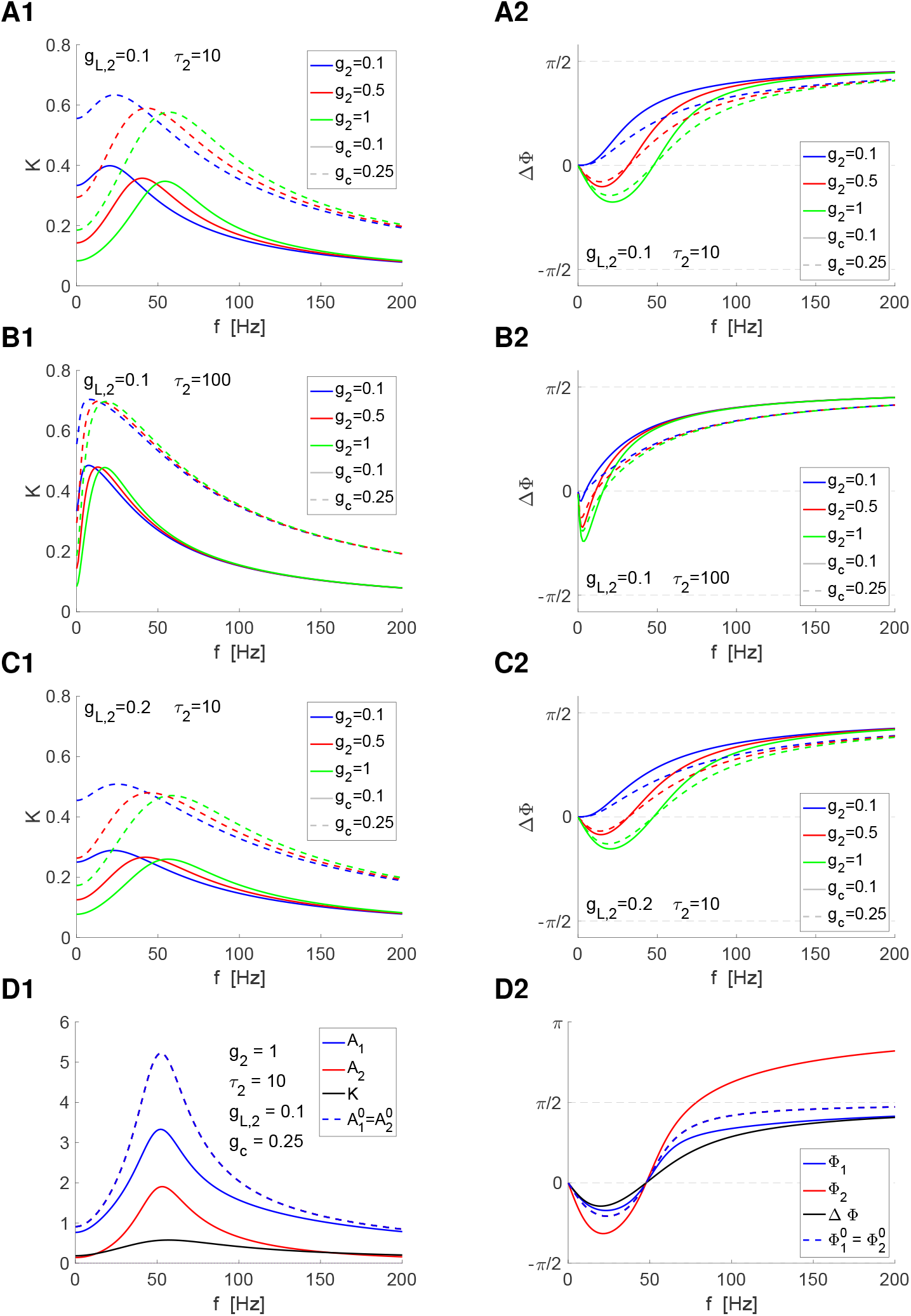
Response of two electrically coupled resonators to oscillatory inputs for representative parameter values. The linear model is given by eqs. (1)-(2). The pre-J and post-J cells (1 and 2, respectively) are identical. **A1, B1, C1.** Coupling coefficients *K*(*f*) given by eq. (28). **A2, B2, C2.** Phase-difference coefficient ΔΦ(*f*) given by eq. (29). **D.** Network amplitude (*A*_1_ and *A*_2_, panel D1) and phase (Φ_1_ Φ_2_, panel D2) responses computed from the complex responses **A_1_** and **A_2_** given by eqs. (7)-(8) for the parameter values used for the solid-green curves in panels A. Superimposed to these profiles are the amplitudes of the individual cells (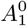 and 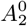, dashed-blue in panel D1) and the coupling coefficient *K* (panel D1), and the phase of the individual cells (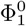 and 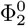, dashed-blue in panel D2) and the phase-difference coefficient ΔΦ (panel D2).

**Figure 7:**
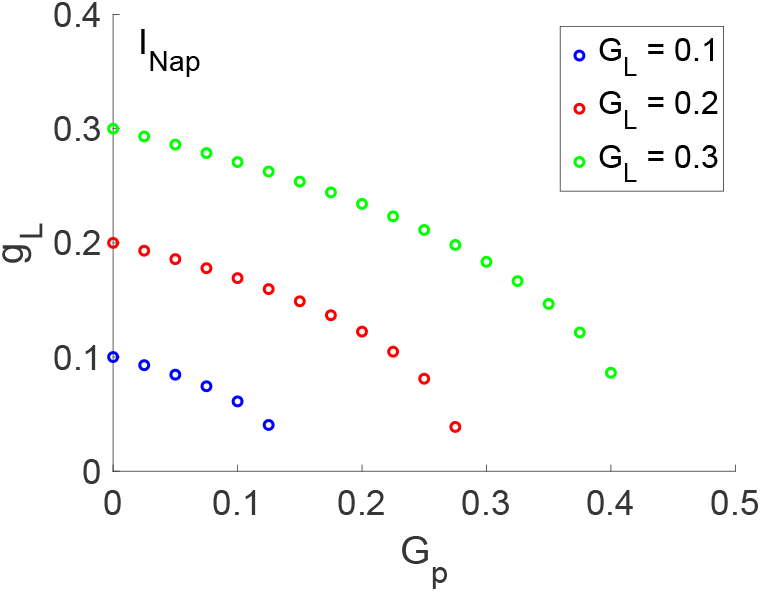
I_Nap_ model: Dependence of the linearized conductance g_L_ on the biophysical conductances G_p_ and G_L_. We used eq. (30) with the parameter values presented in Section S1.2 and *I*_*app*_ = 0.

Figs. 6-A1 to -C1 show the dependence of the *K*(*f*)-profiles on the resonant linearized conductance *g*_2_, the time constant *τ*_2_ of the resonant gating variable, the linearized leak conductance *g*_*L,2*_ and the gap junction coefficient *g*_*c*_. Increasing values of *g*_2_ attenuate the *K*(*f*) profiles and increase the *K*-resonant frequency *f*_*res,K*_ (Fig. 6-A1 to -C1). Increasing values of *g*_*c*_ attenuate the *K*(*f*) profile with no significant change in *f*_*res,K*_ (Fig. 6-A1 to -C1). Increasing values of *τ*_2_ shift *f*_*res,K*_ to lower values, amplify the *K*(*f*) profiles and make the peakier (compare (Figs. 6-A1 and B1). Increasing values of *g*_*L,2*_ attenuate the *K*(*f*) profiles and make them shallower, with no significant differences in *f*_*res,K*_ (compare (Figs. 6-A1 and C1).

Figs. 6-B1 to -D1 show the dependence of the ΔΦ-profiles on *g*_2_, *τ*_2_, *g*_*L,2*_ and *g*_*c*_. Increasing values of *g*_2_, shift the phasonant frequency *f*_*phas*_ to the right and make the changes in the ΔΦ-profiles more pronounced, particularly for the lower frequencies (Fig. 6-A2 to -C2). For the parameter values in Fig. 6-A and *g*_2_ = 0.1, the system exhibits *K*-resonance, but not ΔΦ-phasonance (solid blue). Increasing values of *g*_*c*_ decreases the rate of change of the ΔΦ profiles. Increasing values of *τ*_2_ shift *f*_*phas*_ to lower values (compare (Figs. 6-A2 and B2). Increasing values of *g*_*L,2*_ has a similar effect as increasing the values of *g*_*c*_ (compare (Figs. 6-A2 and C2).

### 3.3 Response to oscillatory inputs: Electrically coupled linearized I_Nap_ + I_h_ and I_Nap_ + I_Ks_ cells

We now turn to the analysis of the dependence of the *K*(*f*) and ΔΦ(*f*) profiles of biophysically plausible electrically coupled (nonlinear) cells on their biophysical constitutive properties by using the results for the electrically coupled linear resonators discussed above. We primarily focus on the *K*-resonant and ΔΦ-phasonant properties. The *I*_*Nap*_ + *I*_*h*_ and *I*_*Nap*_ + *I*_*Ks*_ models and their dynamics are described in Section 2.3 (see also Sections S1.1 and Fig. S4). The linearized *I*_*Nap*_ + *I*_*h*_ and *I*_*Nap*_ + *I*_*Ks*_ models have the form (1)-(2) with *g*_*L*_ and *g* (*g*_*L,2*_ and *g*_2_) defined by eqs. (S14) and (S15) in terms of the biophysical parameters of the corresponding nonlinear models. Our results are presented in Figs. 8 and 10 (*I*_*Nap*_ + *I*_*h*_ model) and Figs. 10 and 11 (*I*_*Nap*_ + *I*_*Ks*_ model). Fig. 7 shows the results for the *I*_*Nap*_ model, a reduced model lacking the slower negative feedback currents.

**Figure 8:**
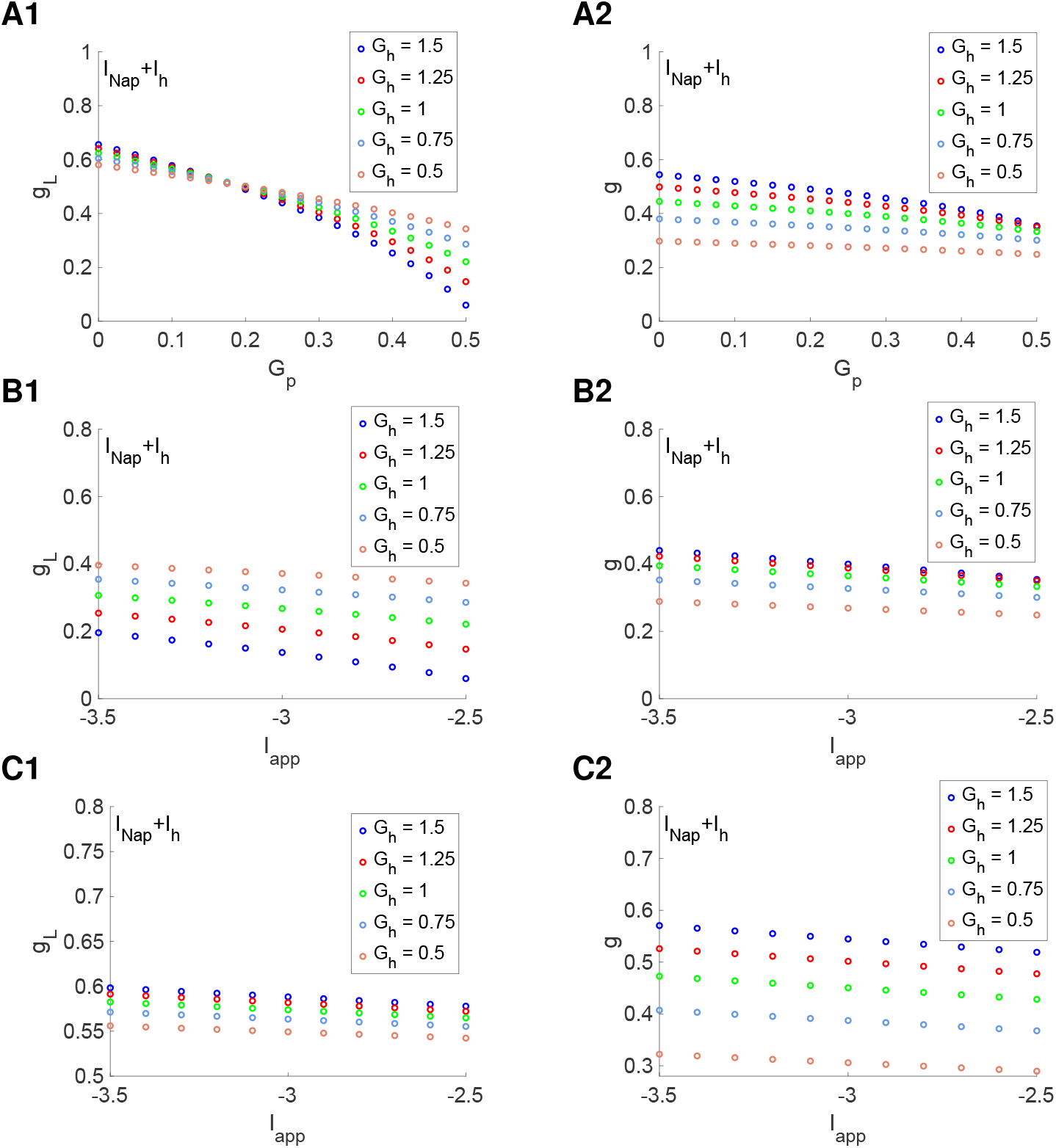
Dependence of the linearized conductances (g_L_ and g) on the biophysical maximal conductances (G_p_ and G_h_) and I_app_ for the I_Nap_+I_h_ model for representative parameter values. We used the biophysical model (S1)-(S2) with the parameter values presented in Section S1.2 and linear model (1)-(2). The linearization process is described in Section S1.3 **A.** *g*_*L*_ and *g* as a function of *G*_*p*_ and *G*_*h*_ for *G*_*L*_ = 0.5 and *I*_*app*_ = *−*2.5. **B.** *g*_*L*_ and *g* as a function of *I*_*app*_ for *G*_*L*_ = 0.5 and *G*_*p*_ = 0.5. **C.** *g*_*L*_ and *g* as a function of *I*_*app*_ for *G*_*L*_ = 0.5 and *G*_*p*_ = 0.1.

**Figure 9:**
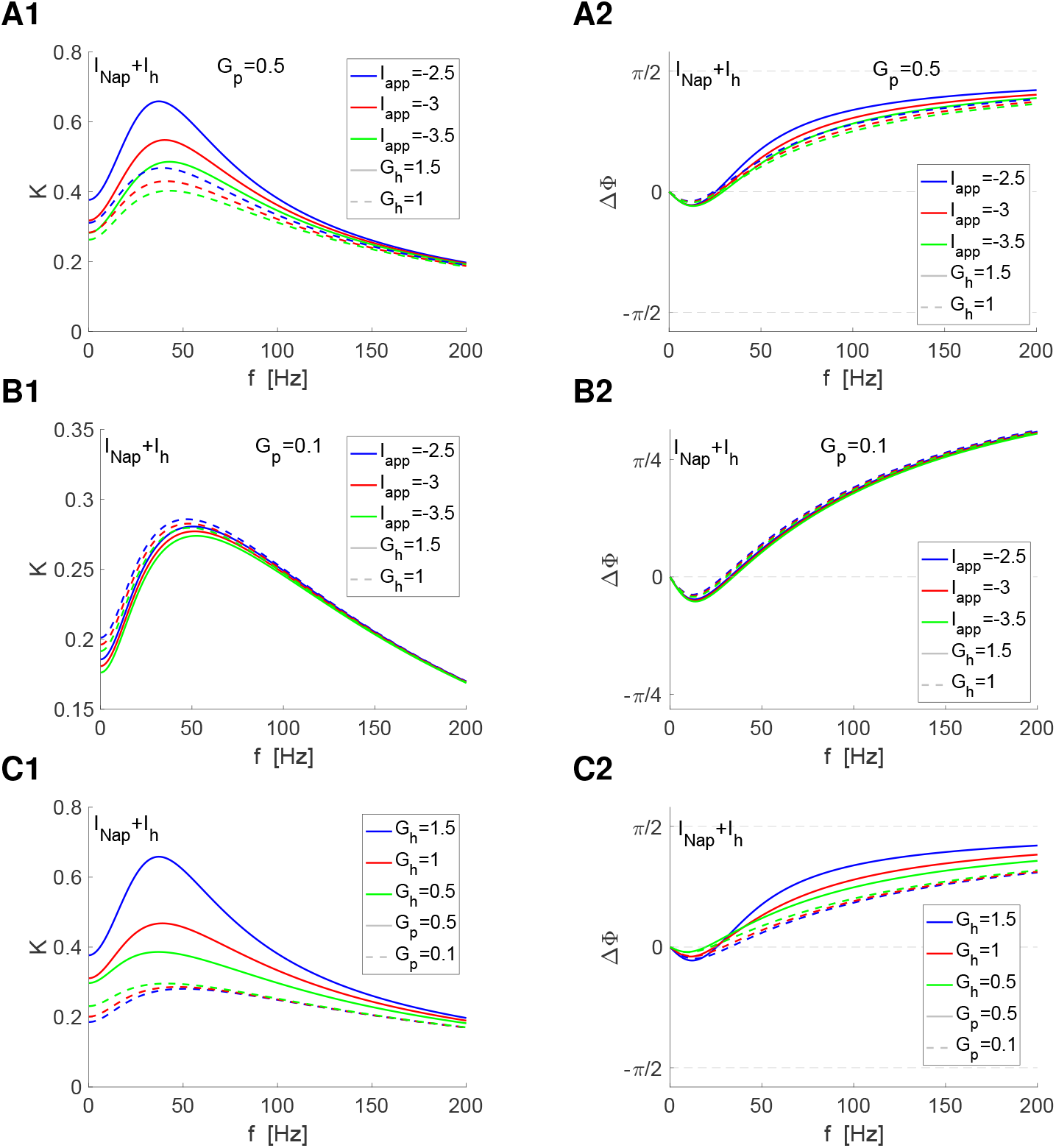
K- and ΔΦ-profiles for the I_Nap_ + I_h_ model for representative parameter values. We used eqs. (28)-(29) for the biophysical *I*_*Nap*_ + *I*_*h*_ model (S1)-(S2) with the parameter values presented in Section S1.2. The linearization process leading to the linearized model (1)-(2) is described in Section S1.3. **Left column.** *K*-profiles. **Right column.** ΔΦ*|*-profiles. **A.** *G*_*p*_ = 0.5, *G*_*L*_ = 0.5. **B.** *G*_*p*_ = 0.1, *G*_*L*_ = 0.5. **C.** *I*_*app*_ = *−*2.5, *G*_*L*_ = 0.5.

**Figure 10:**
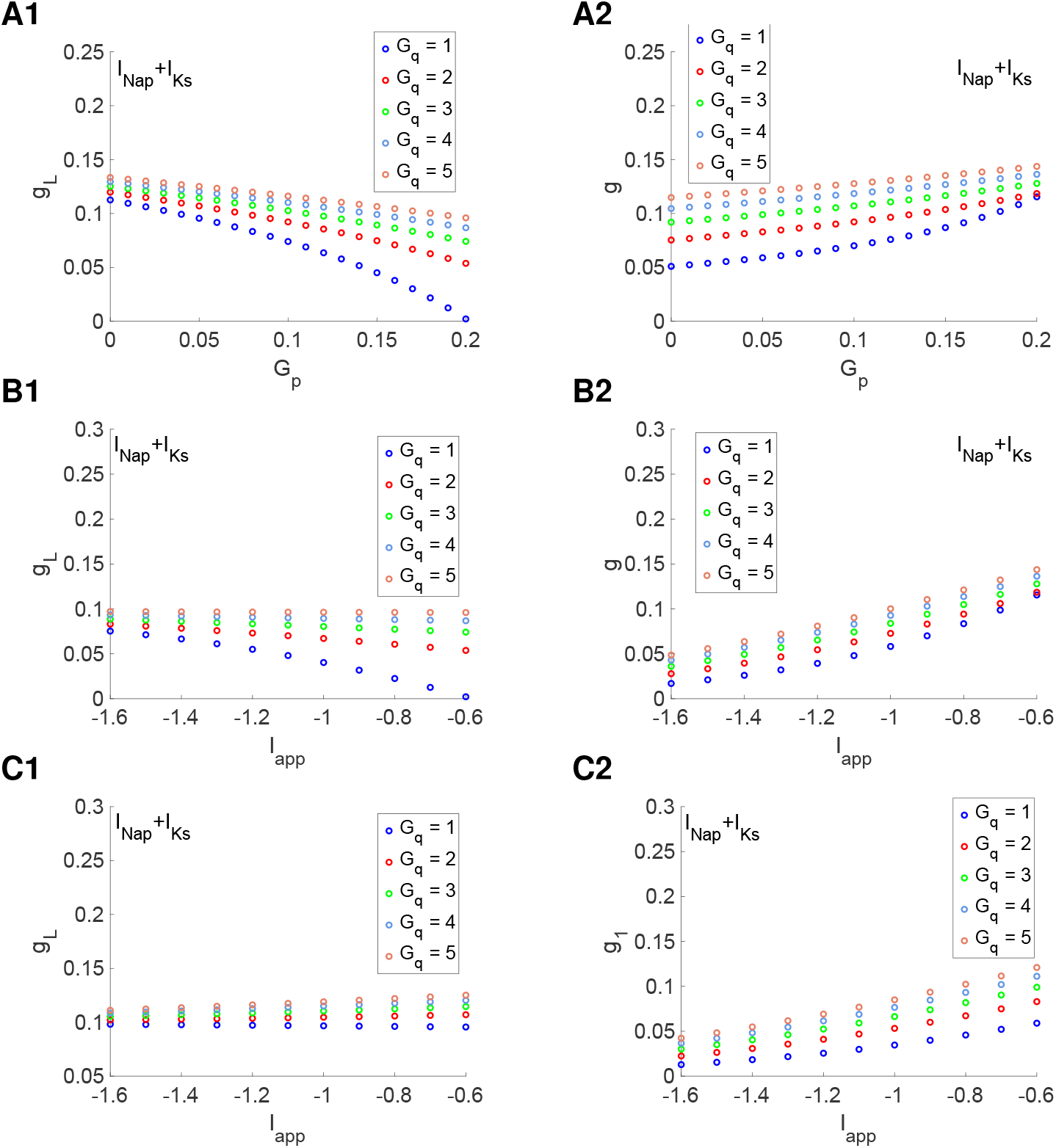
Dependence of the linearized conductances (g_L_ and g) on the biophysical maximal conductances (G_p_ and G_q_) and I_app_ for the I_Nap_+I_Ks_ model for representative parameter values. We used the biophysical model (S1)-(S2) with the parameter values presented in Section S1.2 and linear model (1)-(2). The linearization process is described in Section S1.3 **A.** *g*_*L*_ and *g* as a function of *G*_*p*_ and *G*_*q*_ for *G*_*L*_ = 0.1 and *I*_*app*_ = *−*0.6. **B.** *g*_*L*_ and *g* as a function of *I*_*app*_ for *G*_*L*_ = 0.1 and *G*_*p*_ = 0.2. **C.** *g*_*L*_ and *g* as a function of *I*_*app*_ for *G*_*L*_ = 0.1 and *G*_*p*_ = 0.05.

**Figure 11:**
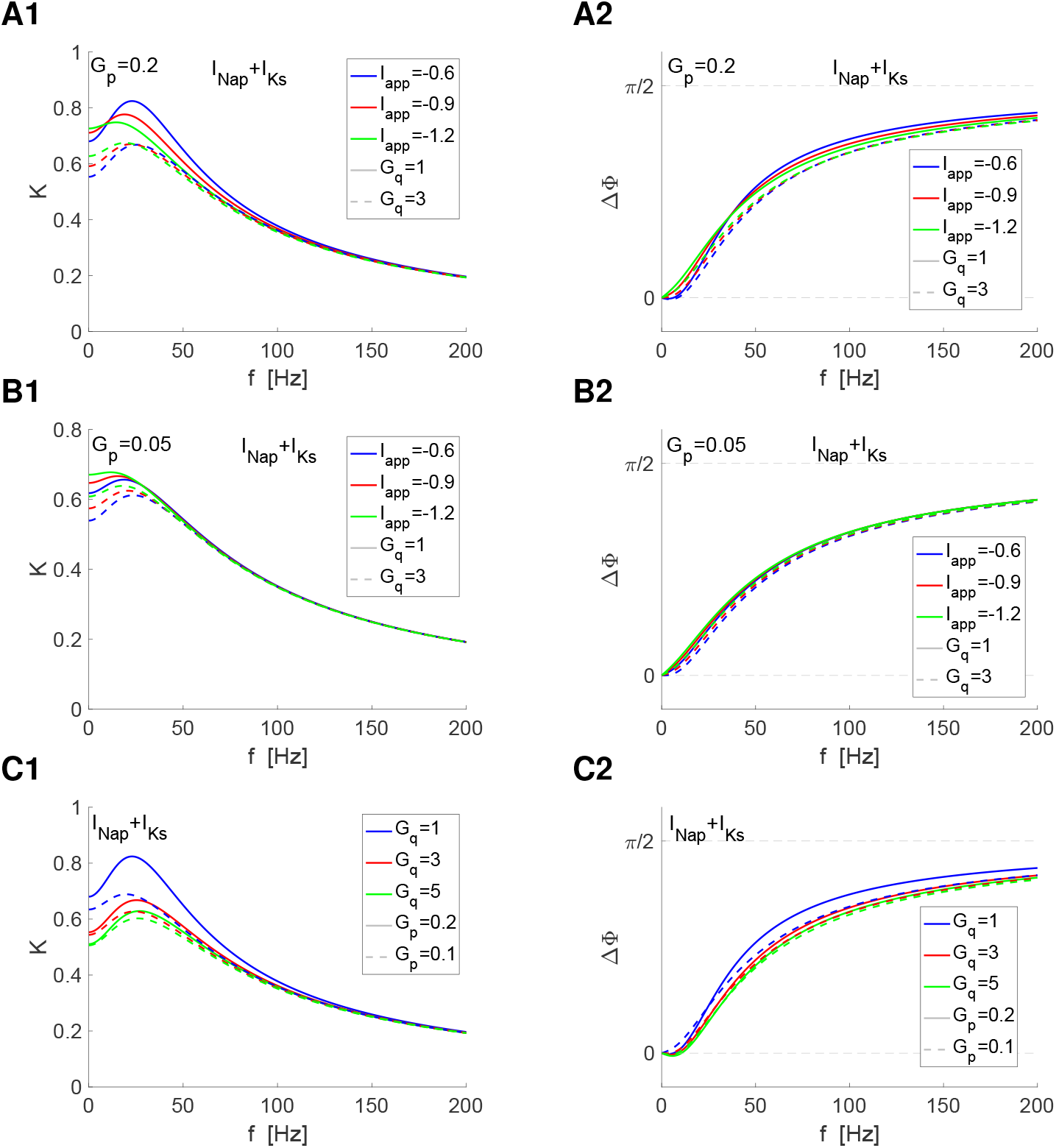
K- and ΔΦ-profiles for the I_Nap_ + I_Ks_ model for representative parameter values. We used eqs. (28)-(29) for the biophysical *I*_*Nap*_ + *I*_*Ks*_ model (S1)-(S2) with the parameter values presented in Section S1.2. The linearization process leading to the linearized model (1)-(2) is described in Section S1.3. **Left column.** *K*-profiles. **Right column.** ΔΦ*|*-profiles. **A.** *G*_*p*_ = 0.2, *G*_*L*_ = 0.1. **B.** *G*_*p*_ = 0.05, *G*_*L*_ = 0.1. **C.** *I*_*app*_ = *−*0.6, *G*_*L*_ = 0.1.

While both *I*_*h*_ and *I*_*Ks*_ are resonant, *I*_*h*_ is hyperpolarization-activated and depolarizing, while *I*_*Ks*_ is depolarization-activated and hyperpolarizing (Fig. S1). These differences are reflected in the geometry of the phase-plane diagram, which are qualitatively mirror images of each other (Figs. S4, S1.2.4 and S6-A1). The parabolic *V* -nullclines are concave up for the *I*_*Nap*_ + *I*_*h*_ model (e.g., Fig. S4-A3, Fig. S6-A1) and concave down for the *I*_*Nap*_ + *I*_*Ks*_ model (e.g., Fig. S1.2.4-A3, Fig. S6-B1). In previous work [21, 66], we showed that these differences translate to differences in some of the models’ oscillatory properties. Here we examine the *K*(*f*) and ΔΦ(*f*) profiles for the two models when electrically coupled and their dependence on the model parameters, and uncover the differences between the two models particularly in the parabolic-like nonlinear regimes.

The excitability level is determined by a combination of the model parameters. For relatively low excitability levels, the fixed-point is relatively far away from the knee of the *V* -nullcline and the system is in a quasi-linear regime (e.g., Figs. S4-A1 and -A2 and Figs. S1.2.4-A1 and -A2). As the excitability level increases, the fixed-point first moves towards the knee of the *V* -nullcline and the parabolic-like nonlinearities become increasingly dominant (e.g., Figs. S4-A3 and S6-A1 and Figs. S1.2.4-A3 and Fig. S6-B1), particularly so in a close vicinity of the bifurcation that produces the onset of spikes. This is reflected in the linearized conductances *g*_*L*_ and *g* (*g*_*L,2*_ and *g*_2_) both explicitly and implicitly through the equilibrium 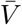. Importantly, changes in *G*_*h*_ or *G*_*q*_ (maximal conductances associated to the resonant currents) explicitly affect both linearized conductances, while changes in *G*_*p*_ (maximal conductance associated to the amplifying current) explicitly affects only the linearized leak conductance *g*_*L*_.

#### 3.3.1 Linearized I_Nap_ model

The *I*_*Nap*_ model (Sections 2.3 and S1.2) is a nonlinear 1D model extending the passive cell to include *I*_*Nap*_ (nonlinear, instantaneously fast). The dynamics of the *I*_*Nap*_ model are illustrated in Fig. S2.

The passive cell can be thought of as the linearization of the *I*_*Nap*_ model with

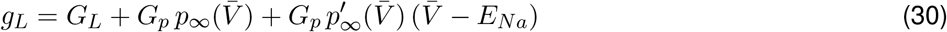

where *G*_*L*_ and *G*_*p*_ are the leak and persistent sodium maximal conductances of the original biophysical model and 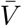 is the equilibrium voltage in the subthreshold regime (if it exists). The last term in eq. (30) is negative, characterizing the amplifying nature of the *I*_*Nap*_ gating variable.

We analyze the dependence of the *K*- and ΔΦ-profiles as well as cells’ response profiles on the model parameters *G*_*L*_ and *G*_*p*_ (to the linear level of approximation) by analyzing the dependence of *g*_*L*_ on these parameters both directly and indirectly through the changes they produced on the equilibrium 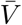.

The dependence of *g*_*L*_ on *G*_*L*_ is straightforward. The dependence of *g*_*L*_ on *G*_*p*_ involves two terms with opposite signs, positive (second) and negative (third). While *g*_*L*_ increases linearly with *G*_*L*_, it decreases with increasing values of *G*_*p*_ (Fig. 7) because of the dominance of the last term over the second one. Geometrically, the *dV/dt* vs. *V* curve becomes shallower as *G*_*p*_ increases. Therefore, increasing values of *G*_*p*_ has the same effect on the network response profiles as decreasing values of *g*_*L*_ in Fig. 3. In particular, increasing values of *G*_*p*_ produces the expected amplification of the *K*-profiles and cause the increase of the ΔΦ profiles to be more pronounced.

#### 3.3.2 Linearized I_Nap_ + I_h_ model

A summary of our findings is presented in Tables 2 and 3 (middle columns). Briefly, increasing levels of *I*_*app*_ cause the model to transition from the quasi-linear to the parabolic-like regimes as the fixed-point moves up along the left branch of the *V* -nullcline towards its knee. As this happens, both *g*_*L*_ and *g* decrease (Figs. 8-B and -C) for fixed values of *G*_*h*_ and *G*_*p*_. Increasing values of *I*_*app*_ cause an amplification of the *K*(*f*) profiles (Figs. 9-A1 and -B1), a decrease in the *K*-resonant frequencies, and a slight increase in the ΔΦ(*f*) profiles (Figs. 9-A2 and -B2). This is more pronounced for *G*_*p*_ = 0.5 than for *G*_*p*_ = 0.1 (compare Figs. 9-A1 and -B1). Increasing values of *G*_*p*_ also cause both *g*_*L*_ and *g* decrease (Figs. 8-A) and therefore the effects are similar to increasing values of *I*_*app*_ (compare Figs. 9-A and -B).

**Table 2:** Dependence of the linearized conductances g_L_ and g on the biophysical parameters I_*app*_, *G*_*p*_, and G_h_ / G_q_.

| As X increases |  |  |
| --- | --- | --- |
| | $I_{Nap} + I_h$ | $I_{Nap} + I_{Ks}$ |
| $X = I_{app}$ | - $g_L$ decreases (Fig. 8-B1, -C1) | - $g_L$ decreases for the higher values of $G_p$ (Fig. 10-B1)<br>- $g_L$ increases for the lower values of $G_p$ (Fig. 10-C1) |
| | - $g$ decreases (Fig. 8-B2, -C2) | - $g$ increases for all values of $G_p$ (Fig. 10-B2, -C2) |
| $X = G_h / G_q$ | - $g_L$ decreases for the higher values of $G_p$ (Fig. 8-B1)<br>- $g_L$ increases for the lower values of $G_p$ (Fig. 8-C1) | - $g_L$ increases for all values of $G_p$ (Fig. 10-B1, -C1) |
| | - $g$ increases for all values of $G_p$ (Fig. 8-B2, -C2) | - $g$ increases for all values of $G_p$ (Fig. 10-B2, -C2) |
| $X = G_p$ | - $g_L$ decreases for all values of $G_h$ (Figs. 8-A1) | - $g_L$ decreases for all values of $G_q$ (Fig. 10-A1) |
| | - $g$ decreases for all values of $G_h$ (Figs. 8-A2) | - $g$ increases for all values of $G_q$ (Fig. 10-A2) |

**Table 3:** Dependence of the properties of the *K* profile on the biophysical parameters I_*app*_, *G*_*p*_, and G_h_ / G_q_. When appropriate, the changes in ΔΦ are reported in absolute value. * When it exists (there are parameter regimes for which ΔΦ(*f*) is always positive).

| As X increases |  |  |
| --- | --- | --- |
| | $I_{Nap} + I_h$ | $I_{Nap} + I_{Ks}$ |
| $X = I_{app}$ | - $K(f)$ is amplified (Fig. 9-A1, -B1) | - $K(f)$ is amplified for the lower values of $G_q$ and the higher values of $G_p$ (Fig. 11-A1, solid)<br>- $K(f)$ is attenuated for the higher values of $G_q$ and the higher values of $G_p$ (Fig. 11-A1, dashed)<br>- $K(f)$ is attenuated for all values of $G_q$ and the lower values of $G_p$ (Fig. 11-B1) |
| | - $f_{res,K}$ decreases (Fig. 9-A1, -B1) | - $f_{res,K}$ increases for the higher values of $G_p$ (Fig. 11-A1, -B1) |
| | - $\Delta\Phi(f)$ increases (Fig. 9-A2, -B2) | - $\Delta\Phi(f)$ decreases for the lower values of $f$ and increases for the higher values of $f$ (Figs. 11-A2) |
| | - $f_{phas,K}$ decreases (Fig. 9-A2, -B2) | - $f_{phas,K}$ increases* (Fig. 11-A2, -B2) |
| $X = G_h / G_q$ | - $K(f)$ is amplified for the higher values of $G_p$ (Figs. 9-C1, solid)<br>- $K(f)$ is attenuated for the lower values of $G_p$ (Figs. 9-C1, dashed) | - $K(f)$ is attenuated (Fig. 11-C1) |
| | - $f_{res,K}$ first increases and then decreases for the higher values of $G_p$ (Figs. 9-C1, solid)<br>- $f_{res,K}$ increases for the lower values of $G_p$ (Figs. 9-C1, dashed) | - $f_{res,K}$ increases (Fig. 11-C1) |
| | - $\Delta\Phi(f)$ increases for the higher values of $G_p$ (Figs. 9-C2, solid)<br>- $\Delta\Phi(f)$ decreases for the lower values of $G_p$ (Figs. 9-C2, dashed) | - $\Delta\Phi(f)$ decreases (Fig. 11-C2) |
| | - $f_{phas,K}$ increases (Figs. 9-C2) | - $f_{phas,K}$ increases* (Figs. 11-C2) |
| $X = G_p$ | - $K(f)$ is amplified (Figs. 9-C1) | - $K(f)$ is amplified (Figs. 11-C1) |
| | - $f_{res,K}$ decreases (Figs. 9-C1) | - $f_{res,K}$ increases (Figs. 11-C1) |
| | - $\Delta\Phi(f)$ increases (Figs. 9-C2) | - $\Delta\Phi(f)$ decreases for the lower values of $f$ and increases for the higher values of $f$ (Figs. 11-C2) |
| | - $f_{phas,K}$ decreases (Figs. 9-C2) | - $f_{phas,K}$ increases* (Figs. 11-C2) |

The nonlinear effects become apparent for the largest values of *I*_*app*_ and *G*_*p*_ for which the fixed-point of the individual cells is located in a close vicinity of the knee of the parabolic-like nullcline, while for lower values of *I*_*app*_ and *G*_*p*_ the system is in a quasi-linear regime.

The dependence of the linearized conductances with *G*_*h*_ is more complex. Increasing values of *G*_*h*_ cause an increase in *g* for all values of *G*_*p*_ as expected from a resonant conductance (a maximal conductances associated to a resonant current) (Fig. 8-A2). However, the dependence of *g*_*L*_ with *G*_*h*_ switches for some intermediate value of *G*_*p*_. For the higher values of *G*_*p*_, *g*_*L*_ decreases with increasing values of *G*_*h*_ (from light-salmon to blue in Fig. 8-A1) for the higher values of *G*_*p*_, while for the lower values of *G*_*p*_, *g*_*L*_ increases with increasing values of *G*_*h*_. This behavior implies that the *K*-profile is nonlinearly amplified by increasing values of *G*_*h*_ for the higher values of *G*_*p*_ (Fig. 9-C1, solid), while it is attenuated by increasing values of *G*_*h*_ for the lower values of *G*_*p*_ where the model is quasi-linear (Fig. 9-C1, dashed). Similarly, as *G*_*h*_ increases, the *K*-resonant frequency decreases for the higher values of *G*_*p*_ and increases for the lower values of *G*_*p*_. A qualitatively switch in behavior occurs for the ΔΦ-profiles (Fig. 9-C2).

In other words, the presence of *I*_*Nap*_ in the nonlinear regime causes the resonant current *I*_*h*_ to amplify the *K*-profile, while it would otherwise attenuate it in the quasi-linear regime where the levels of *I*_*Nap*_ are lower.

#### 3.3.3 Linearized *I*_*Nap*_ + *I*_*Ks*_ model

A summary of our findings is presented in Tables 2 and 3 (right columns). Briefly, increasing levels of *I*_*app*_ cause the individual neurons to transition from quasi-linear to parabolic-like as the fixed-point moves down along the left branch of the *V* -nullcline towards its knee. As this happens, *g*_*L*_ decreases for the higher values of *G*_*p*_ (Fig. 10-B1) and increases for the lower values of *G*_*p*_ (Fig. 10)-C1), while *g* increases for all values of *G*_*p*_ (Fig. 10)-B2 and -C2). This behavior is qualitatively similar for fixed representative values of *G*_*q*_ and *G*_*p*_ although in some cases the dependencies with *I*_*app*_ are weak. As *I*_*app*_ increases for the higher values of *G*_*p*_, the *K*-profiles are amplified for the higher values of *G*_*q*_ (Fig. 11-A1, solid) and attenuated for the lower values of *G*_*q*_ (Fig. 11-A1, dashed). In contrast, for the lower values of *G*_*p*_, the *K*-profiles are attenuated for all values of *G*_*q*_ as *I*_*app*_ increases (Fig. 11-B1). In both cases, the *K*-resonant frequency increases with increasing values of *I*_*app*_. A similar type of switch can be observed in the ΔΦ-profiles although they are minimally affected and the differences are almost imperceptible (Figs. 11-A2 and -B2).

For fixed values of *I*_*app*_ and *G*_*p*_, both *g*_*L*_ and *g* increase with increasing values of *G*_*q*_ (Fig. 10-A to -C). The *K*-profiles are attenuated and the *K*-resonant frequency increases with increasing values of *G*_*q*_ (Fig. 11-C1). For fixed values of *I*_*app*_ and *G*_*q*_, both *g*_*L*_ and *g* increase with increasing values of *G*_*p*_ (Fig. 10-A1). As expected, the *K*-profiles are amplified by increasing values of *G*_*p*_ (Fig. 11-C1).

Note that for certain parameter regimes the ΔΦ(*f*) profile is always positive and therefore the network does not exhibit ΔΦ-phasonance (e.g, Fig. 11-B2).

#### 3.3.4 The electrically coupled I_Nap_ + I_h_ and I_Nap_ + I_Ks_ cells have different response properties that are captured by the linearized models

Together, the results of the previous two sections highlight the fact that although the two biophysical models involve the interplay of resonant and amplifying currents, there exist significant qualitative differences between the models’ *K*(*f*) and ΔΦ(*f*) profiles and their dependence with the model parameters and levels of excitability. As expected, the two biophysical models have a relatively similar dependence with the levels of the participating currents (model parameters) in the quasi-linear regimes where the excitability levels are relatively low and the behavior is dominated by the resonant currents (*I*_*h*_ and *I*_*Ks*_). However, this changes significantly when the excitability levels are higher and the models are in a nonlinear, parabolic-like regime in the presence of higher levels of *I*_*Nap*_. In these regimes, the synergistic interaction between the resonant and amplifying currents produce unexpected results that cannot be predicted from the quasilinear behavior nor from the nature of the resonant currents such as the amplifying effects of the resonant current *I*_*h*_. The similarities and differences between the two models are summarized in Tables 2 and 3.

### 3.4 Electrically coupled two-compartmental models of spatially extended neurons: Effects of asymmetry in the connectivity

Here, we extend our investigation to two-compartment neuron models having asymmetric connectivity and network heterogeneity. More specifically, we consider two-compartment models of linear or linearized cells where the oscillatory in-put arrives only to one of the compartments (indexed by 1). The coupling coefficient 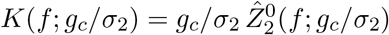 and the phase-difference coefficient 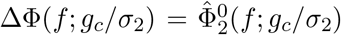 for two-compartment linear models are presented in Section 2.7 where 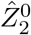 and 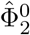 are the extended impedances of the individual (disconnected) compartments. Both depend on the ratio of *g*_*c*_/*σ*_2_ where *g*_*c*_ represents the factor in the coupling strength between the two compartments that is common to both and *σ*_2_ represents the proportion of the total cell are taken up by compartment 2. Specific expressions for these quantities can be computed from eqs. (26) and (27), respectively, when compartment 2 is passive, and by eqs. (28) and (29), respectively when compartment 2 is a 2D resonator.

Because these quantities depend on the quotient *g*_*c*_/*σ*_2_, it is enough to examine 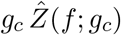 and 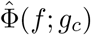 for varying values of *g*_*c*_ and arbitrary extended shapes of the impedance and phase profiles. As mentioned above, the dependence of these expressions on *g*_*c*_ consist of a deformation and an additional rescaling by the same quantity (*g*_*c*_) for the firsts one. The results of the previous sections for electrically coupled cells extend to two-compartment models in a straightforward way by performing these operations. They do not affect the relative values of the profiles nor the mechanisms by which they are generated.

As *g*_*c*_ increases, 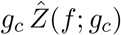 is amplified and becomes shallower (Fig. 12-A1 to -C1) and 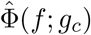 decreases in absolute value (Fig. 12-A2 to -C2). The dependence of these quantities with *σ*_2_ can be inferred from these results for fixed values of *g*_*c*_ as *σ*_2_ decreases. We note that the primary effect of increasing *g*_*c*_ on these extended quantities is to attenuate the impedance profile (Fig. S7-A1 to -C1) and make it shallower, and to decrease (in absolute value) the phase profile (Fig. S7-A1 to -C1).

**Figure 12:**
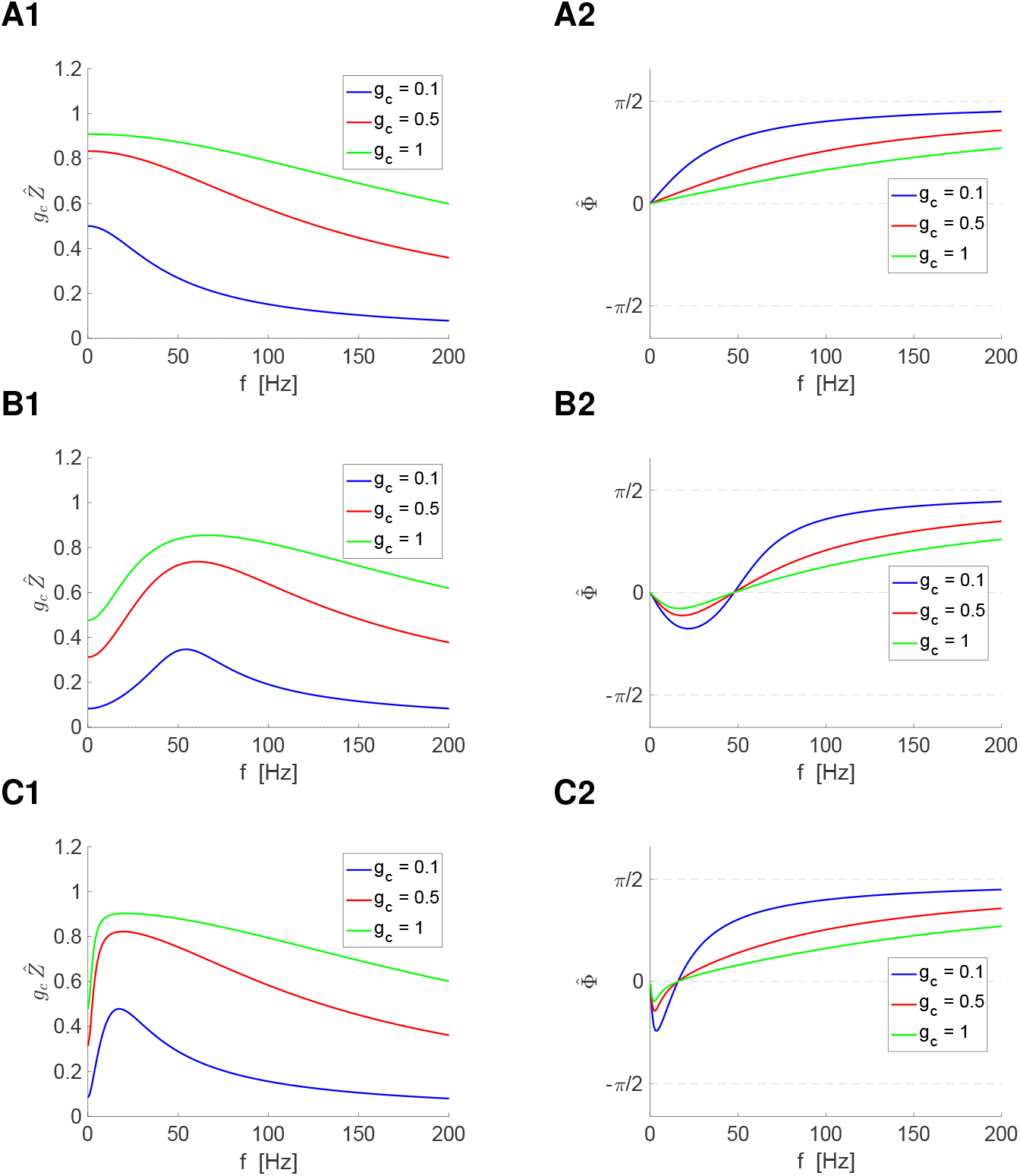
Extended impedance 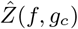 multiplied by *g*_*c*_ and phase 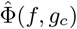 profiles. We used eqs. (S53)-(S54). For *g* = 0, these eqs. reduce to eqs. (S55)-(S56). These quantities are equivalent to the coupling coefficients *K*(*f*) and phase-difference coefficient ΔΦ. **Left columns.** Extended impedance profiles multiplied by *g*_*c*_. **Right columns.** Extended phase profiles. **A.** *g* = 0. **B.** *g* = 1, *τ* = 10. **C.** *g* = 1, *τ* = 100. We used the additional parameter value: *g*_*L*_ = 0.1.

### 3.5 Nonlinear models: The coupling and phase-difference coefficients depend on the interplay of both the pre- and post-J cells

For linear models, both the *K*(*f*) and ΔΦ(*f*) profiles depend on the properties of the post-J cell, and are independent of the properties of the pre-J cell and the input amplitude. Analytical results for electrically coupled weakly nonlinear cells show that both the pre- and post-J cell and the input amplitude contribute to shape these two profiles (see Section S4). Here we test how the *K*(*f*) and ΔΦ(*f*) profiles are shaped by the intrinsic nonlinearities of quadratic type (Section 2.2; Fig. 3, Fig. S3) in the pre-J and post-J cells and the input amplitude.

We use the quadratic model (3)-(4) where the connectivity parameters are *g*_*c,k*_ = *g*_*c*_ for electrically coupled cells mediated by gap junctions and *g*_*c,k*_ = *g*_*c*_/σ_*k*_ for two-compartmental models. For the 1D reduced model (no negative feedback), we use a slightly modified version of the model, which is described below. The description of the quadratization process linking the parameters of the quadratic models with these of biophysical models with nonlinearities of parabolic-type in the voltage equation is presented Section S1.4 [70–72].

#### 3.5.1 Electrically coupled 1D quadratic cells: interplay of nonlinear LPFs

For cells with 1D quadratic dynamics, the electrically coupled cell model reads

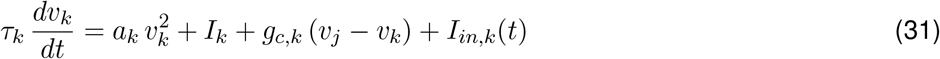

for *k, j* = 1, 2, *k≠ j* where *I*_*k*_ represents the level of excitability, and *a*_*k*_ captures the effect of an instantaneously fast amplifying current (e.g., *I*_*Nap*_). As before, the time-dependent oscillatory inputs are given by eq. (5) with *I*_*in,2*_ = 0.

In the presence of *I*_*Nap*_ the cell develops nonlinearities of parabolic-type (Fig. S2). For relatively low levels of *I*_*Nap*_, the nonlinearities are away from the equilibrium and therefore the systems is quasi-linear. As the levels of *I*_*Nap*_ increase, the equilibrium becomes closer to the knee of the parabolic nullcline and to the unstable fixed-point (who serves as a threshold for spike generation when the model is supplemented with a spiking mechanism). For a critical level of *I*_*Nap*_, the two fixed-points collide in a bifurcation and disappear. In the quadratic models, the excitability levels are represented by the DC current *I*_*k*_.

The quadratic 1D model is the subthreshold description of the well-known quadratic integrate-and-fire model [79, 80, 90, 91]. Fig. S3 illustrates the dynamics of an individual quadratic cell (*g*_*c*_ = 0 and *I* representing the DC current). As *I* increases, the speed (*dV/dt* vs. *V*) curve increases, the fixed-point moves to the right and the range of values of *v* for which the response to oscillatory inputs remains in the subthreshold regime shrinks. We choose values of *A*_*in*_ so that the response for all input frequencies remains in the subthreshold regime. The effects of the nonlinearities are apparent in the asymmetry, non-sinusoidal shape of the voltage response (compare the response waveforms in Figs. S3-A2 and -B2 and the *V*_*max*_ and *V*_*min*_ envelope responses in Figs. S3-A3 and -B3).

In Fig. 13 we compare the *K*(*f*) and ΔΦ(*f*) profiles for two identical connected cells (blue) and two different connected cells (red) where cell 2 is the same as for the identical pair and cell 1 has the same parameter values as cell 2, except for one that varies across panels. If the model were linear, we would expect the resulting graphs to be superimposed. The differences observed in Fig. 13 indicate that the *K*(*f*) and ΔΦ(*f*) profiles are determined by the interplay of both participating cells. These differences are stronger for the *K*(*f*) than for the ΔΦ(*f*) profiles. For the *K*(*f*) profiles, the differences are stronger for the lower frequencies and decrease as the frequency increases. Because the two cells are network LPFs (not shown) their voltage response range shrinks as the input frequency increases. For the higher frequencies, the response trajectory (e.g., blue line in Fig. S3) reaches the region where the parabolic nonlinearities are prominent. For the lower frequencies, the response trajectory remains in a close vicinity of the fixed point where the dynamics are quasi-linear. These results persists for two-compartment models (not shown).

**Figure 13:**
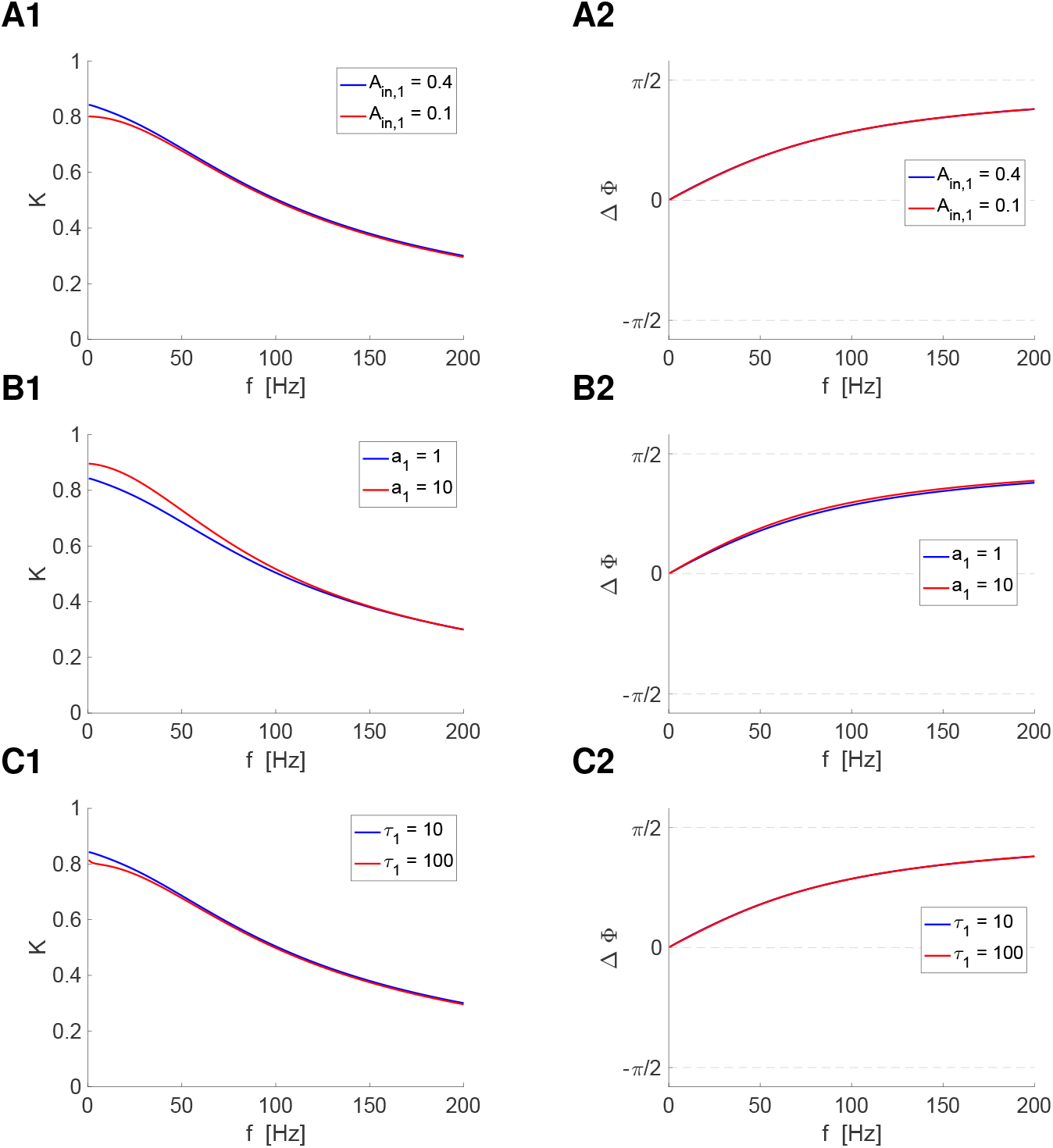
Electrically coupled quadratic 1D model: Effects of the pre-J cell’s properties on the response profiles. We used the model (31) for the electrically coupled pre- and post-J cells with quadratic nonlinearities (cell 1 is the pre-J cell; *A*_*in,2*_ = 0) mediated by gap junctions (*g*_*c*_ = 0.5). The parameter values for the two cells are identical, except for the ones indicated in the legends. **Left column.** *K* profiles **Right column.** ΔΦ profiles **A.** *a*_1_ = *a*_2_ = 1, *τ*_1_ = *τ*_2_ = 10. **B.** *τ*_1_ = *τ*_2_ = 10, *A*_*in,1*_ = 0.4. **C.** *a*_1_ = 1 = *a*_2_ = 1, *A*_*in,1*_ = 0.4. We used the following additional parameter values: *I*_1_ = *I*_2_ = *−*0.25 and *g*_*c*_ = 4

#### 3.5.2 Electrically coupled 2D quadratic cells with negative feedback effects: interplay of nonlinear BPFs, LPFs and time scales

Figs. 14 llustrates the nonlinear (amplitude) BPFs (panels A1, C1, D1) and LPFs (panel B1) for the quadratic model for an individual cell for representative parameter values. In addition to resonance, the BPF’s systems exhibit phasonance (panels A2, C2, D2), while the LPF’s system exhibits no phasonance (panel B2). The filters’ nonlinearity is apparent in the lack of symmetry of the peak (upper) and trough (lower) envelopes with respect to the equilibrium (gray dashed horizontal line) and the lack of proportionality with respect to the input amplitude *A*_*in*_ (e.g., Fig. 14-A1, compare the solid-blue response to *A*_*in*_ = 0.05 and the dashed-blue response for *A*_*in*_ = 0.01). The effects of the model’s quadratic nonlinearity are reflected in the shapes of the response limit cycles for the resonant frequency (Figs. 14-A3, solid) [71]. For smaller values of the input amplitude *A*_*in*_, the response limit cycle remains in a close vicinity of the fixed-point (Figs. 14-A3, dashed) and the peak and trough envelopes are symmetric with respect to the equilibrium (Figs. 14-A1, dashed).

**Figure 14:**
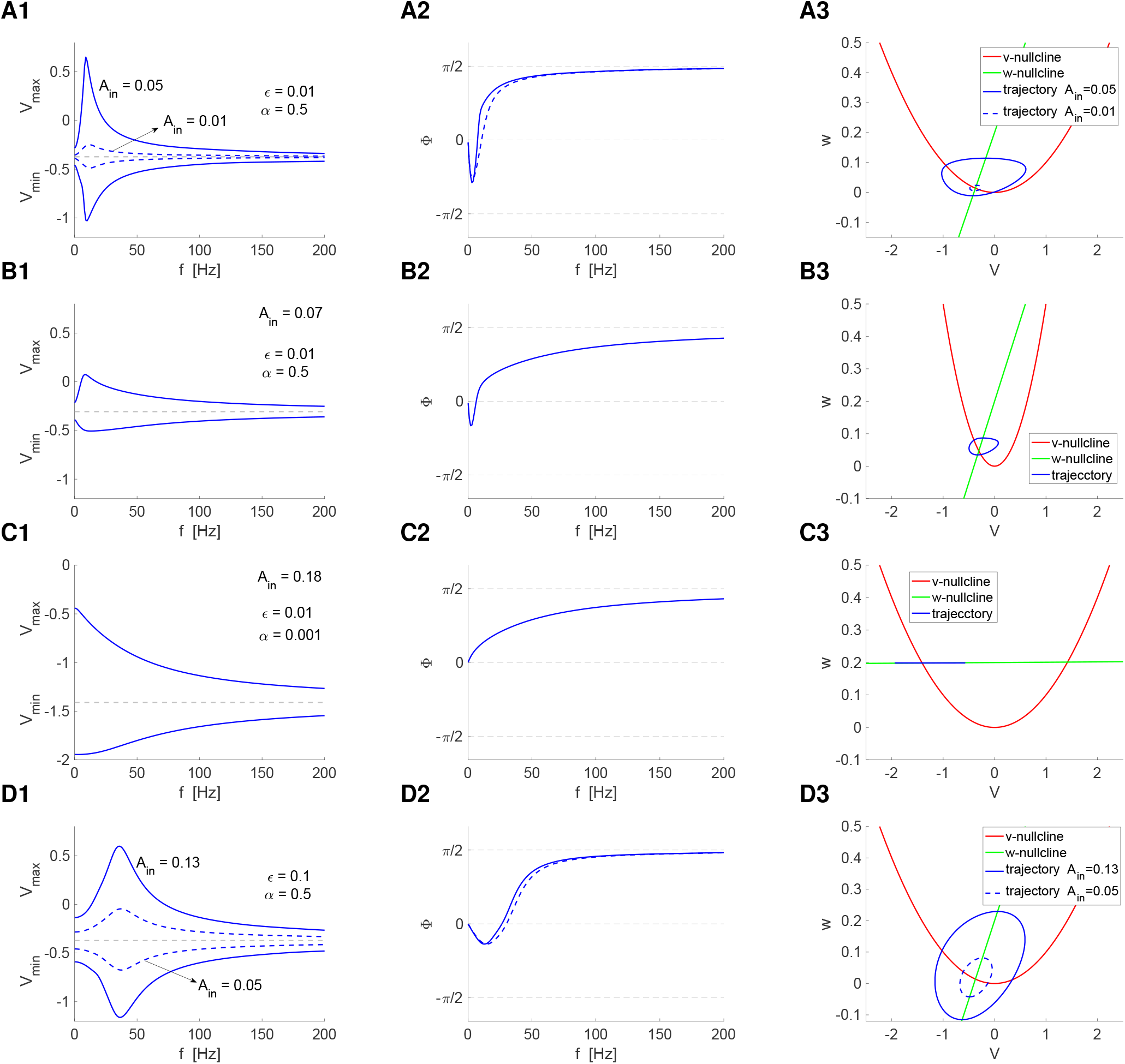
Response of the individual quadratic 2D cells to oscillatory inputs: Representative nonlinear BPFs and LPFs. We used the quadratic model for an individual cell (3)-(4) (*k* = 1, *g*_*c*_ = 0). **Left column.** Peak (*V*_*max*_) and trough (*V*_*min*_) envelope profiles **Middle column.** Phase (Φ) profiles **Right column.** Phase-space diagrams for the peak / resonant frequency. The *v*- and *w*-nullclines are given by *w* = *a v*^2^ and *w* = *α v − λ*, respectively. The response trajectory is a projection of the three-dimensional space onto the *v*-*w* plane. **A.** *α* = 0.5, *ϵ* = 0.01, *a* = 0.1, *A*_*in*_ = 0.05 (solid) and *A*_*in*_ = 0.01 (dashed). The dashed curve is presented for comparison between the responses of the nonlinear (solid) and quasi-linear (dashed) systems for low enough values of *A*_*in*_. The solid curve corresponds to approximately the highest value of *A*_*in*_ for which the response remains within the subthreshold regime for all input frequencies.. **B.** *α* = 0.5, *ϵ* = 0.01 *a* = 0.5 and *A*_*in*_ = 0.01. The parabolic nonlinearity is more pronounced and the response nonlinear. **C.** *α* = 0.001, *ϵ* = 0.01, *a* = 0.1 and *A*_*in*_ = 0.18. The system is quasi-1D as the result of the disruption of the activation of the negative feedback term. For lower values of *A*_*in*_ (e.g., *A*_*in*_ = 0.05 as the one in panel A), the LPF is quasi-symmetric reflecting the response of a quasi-linear system (not shown). **D.** *α* = 0.5, *ϵ* = 0.1, *a* = 0.1, *A*_*in*_ = 0.13 (solid) and *A*_*in*_ = 0.05 (dashed). The BPF is quasi-symmetric reflecting the underlying quasi-linear dynamics. The dashed curve is presented for comparison with the BPF in panel A. We use the following additional parameter values: *a* = 0.1, *λ* = *−*0.2.

The presence of the model’s nonlinearities do not necessarily imply a strongly nonlinear response such as the one in Fig. 14-A1 (solid) for *A*_*in*_ = 0.05. Not only because weakly nonlinear responses (quasi-linear filters) are obtained for lower values of *A*_*in*_ that are not necessarily small enough (e.g., Fig. 14-A1, dashed, for *A*_*in*_ = 0.01), but also because of other factors such as a small enough negative feedback time constant (higher value of *ϵ*) (e.g., Fig. 14-D1). Comparison between the BPFs in Figs. 14-A1 (solid) and 14-D1 (dashed), for the same value of *A*_*in*_ shows that the BPFs are more symmetric and more attenuated, the higher *ϵ* [71]. This is the result of the interplay of the oscillatory inputs and the underlying vector field [71] (compare Figs. 14-A3 and -D3). BPFs with a comparable peak value requires a higher value of *A*_*in*_ for *ϵ* = 0.1 than for *ϵ* = 0.01. An increase in the parabolic curvature *a* (Fig. 14-B3) produces a nonlinear response (Fig. 14-B1). However, the filter’s amplitude is smaller as compared to the one for lower values of *a* (Fig. 14-A1) since the range of voltage values for which the response remains within the subthreshold voltage regime shrinks as *a* increases.

In our study, we use the parameter values leading to the amplitude BPF in Fig. 14-A1 as the baseline parameter set (*a* = 0.1, *α* = 0.5 and *ϵ* = 0.01), and we compare a homogeneous network consisting of two identical baseline cells with heterogenous, mixed-cells networks where one cell is a baseline cell and the other has the same parameter values as the baseline cell, except for one. We use the representative parameter values in Figs. 14-B (increase in *a*, change in the parabolic nonlinearity), 14-C (decrease in *α*, LPF), and 14-D (increase in *ϵ*, change in the negative feedback time constant).

In Fig. 15 we test whether the *K*(*f*) and ΔΦ(*f*) profiles depend on the properties of the pre-J cell (Fig. 15-B-D) and the input amplitude *A*_*in*_ (Fig. 15-A) in addition to the post-J cell. To this end, we proceed as in the previous Section and compare the *K*(*f*) and ΔΦ(*f*) profiles of a homogeneous and heterogeneous networks as described above. Our results demonstrate that the *K*(*f*) and ΔΦ(*f*) profiles depend on *A*_*in*_ and on the properties of the pre-J cell (in addition to the properties of the post-J cell). In some cases, the differences in the profiles between the homogeneous and heterogenous networks is significant, more so than for the quadratic 1D models discussed in the previous section. These differences are more prominent for the lower frequencies and the frequencies in the resonant frequency band where the response amplitude is larger and therefore the response trajectories in the phase-space diagram are able to reach the region of parabolic nonlinearities. For the input frequencies for which the response amplitude is smaller, the trajectories evolve in a closer vicinity of the fixed-point and therefore the dynamics is quasi-linear. In Fig. 14-C we observe that the *K*(*f*) profile for the heterogenous network exhibits *K*-antiresonance in addition of *K*-resonance, and the ΔΦ(*f*) profile exhibits ΔΦ-antiphasonance in addition to the ΔΦ-phasonance. This is the result of a network effect, since neither antiresonance nor antiphasonance are present in any of the participating cells (these phenomena require 3D cellular dynamics).

**Figure 15:**
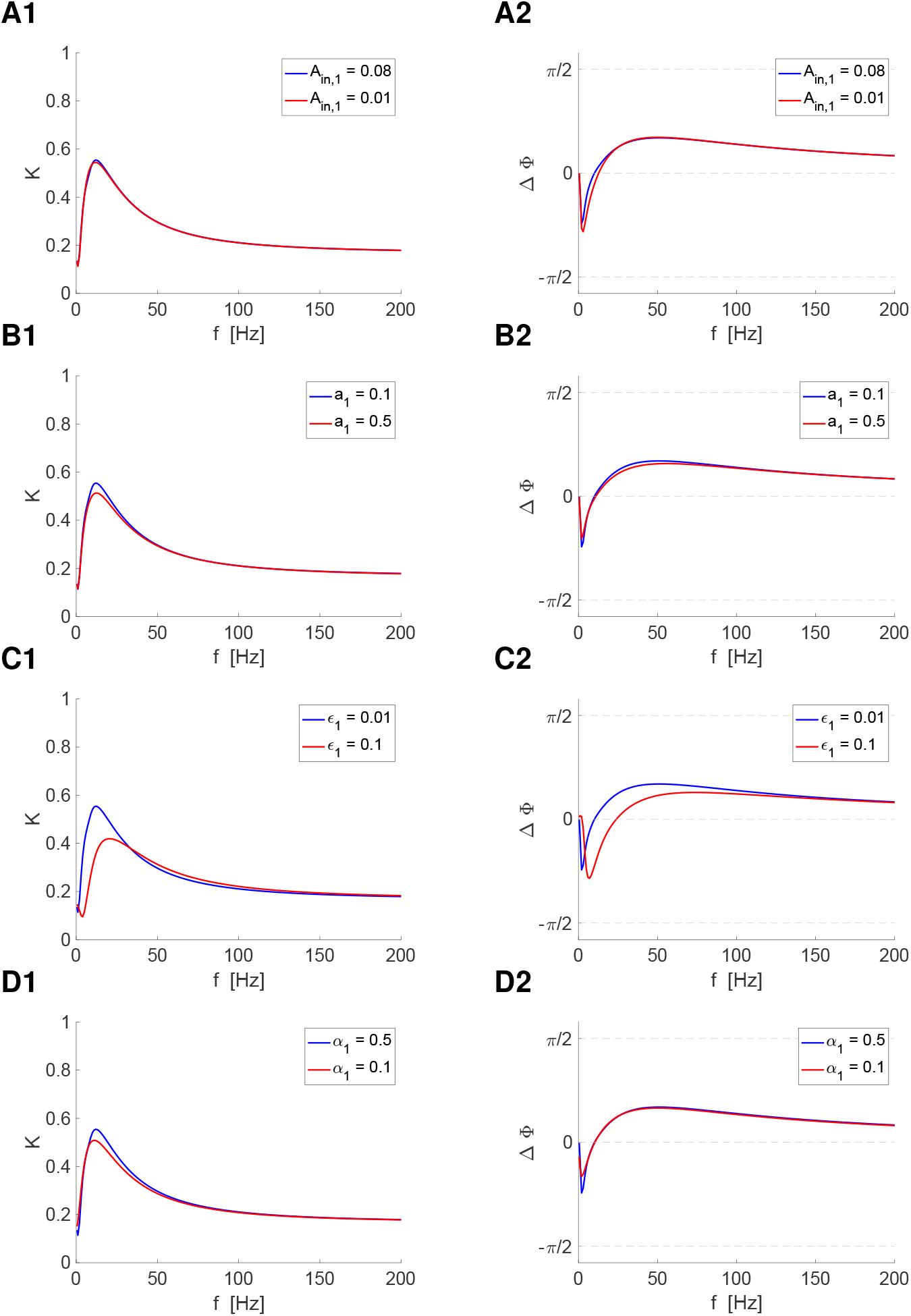
Electrically coupled quadratic 2D model: Effects of the pre-J cell’s properties on the response profiles. We used the quadratic model (3)-(4) for the electrically coupled pre- and post-J cells with quadratic nonlinearities mediated by gap junctions (cell 1 is the pre-J cell; *A*_*in,2*_ = 0). The parameter values for the two cells are identical, except for the ones indicated in the legends. **Left column.** *K* profiles **Right column.** ΔΦ profiles **A.** *a*_1_ = *a*_2_ = 0.1, *α*_1_ = *α*_2_ = 0.5, *ϵ*_1_ = *ϵ*_2_ = 0.01. **B.** *a*_2_ = 0.1, *α*_1_ = *α*_2_ = 0.5, *ϵ*_1_ = *ϵ*_2_ = 0.01, *A*_*in,1*_ = 0.08. **C.** *a*_1_ = *a*_2_ = 0.1, *α*_1_ = *α*_2_ = 0.5, *ϵ*_2_ = 0.01, *A*_*in,1*_ = 0.08. **D.** *a*_1_ = *a*_2_ = 0.1, *α*_2_ = 0.5, *ϵ*_1_ = *ϵ*_2_ = 0.01, *A*_*in,1*_ = 0.08. We used the following additional parameter values: *λ*_1_ = *λ*_2_ = *−*0.2, *g*_*c*_ = 0.1 and *A*_*in,2*_ = 0.

In Figs. 16 and 17 we extend our results and test the effects of the input location on the *K*(*f*) and ΔΦ(*f*) profiles for the heterogeneous networks investigated in Fig. 15. Because in our formulation, the input always arrives to the cell indexed by 1 (the pre-J cell), we switch the parameter values of cells 1 and 2. Specifically, the curves shown in these figures correspond to the homogeneous network of baseline cells (blue), and the two heterogeneous network where cell 2 (cell 1) is a baseline cell and cell 1 (cell 2) has the same parameter values as the baseline cell, except for one, which varies across panels (red and green, respectively). In all three cases in Fig. 16 the *K*(*f*) and ΔΦ(*f*) profiles depend on the properties of both the pre-J and post-J cell and are not necessarily dominated by the properties of the post-J cell. This persist for larger values of *g*_*c*_ (Figs. 17) and for two-compartment models where the effects of the compartments’ area renders the connectivity asymmetric (Figs. 18 and 19).

**Figure 16:**
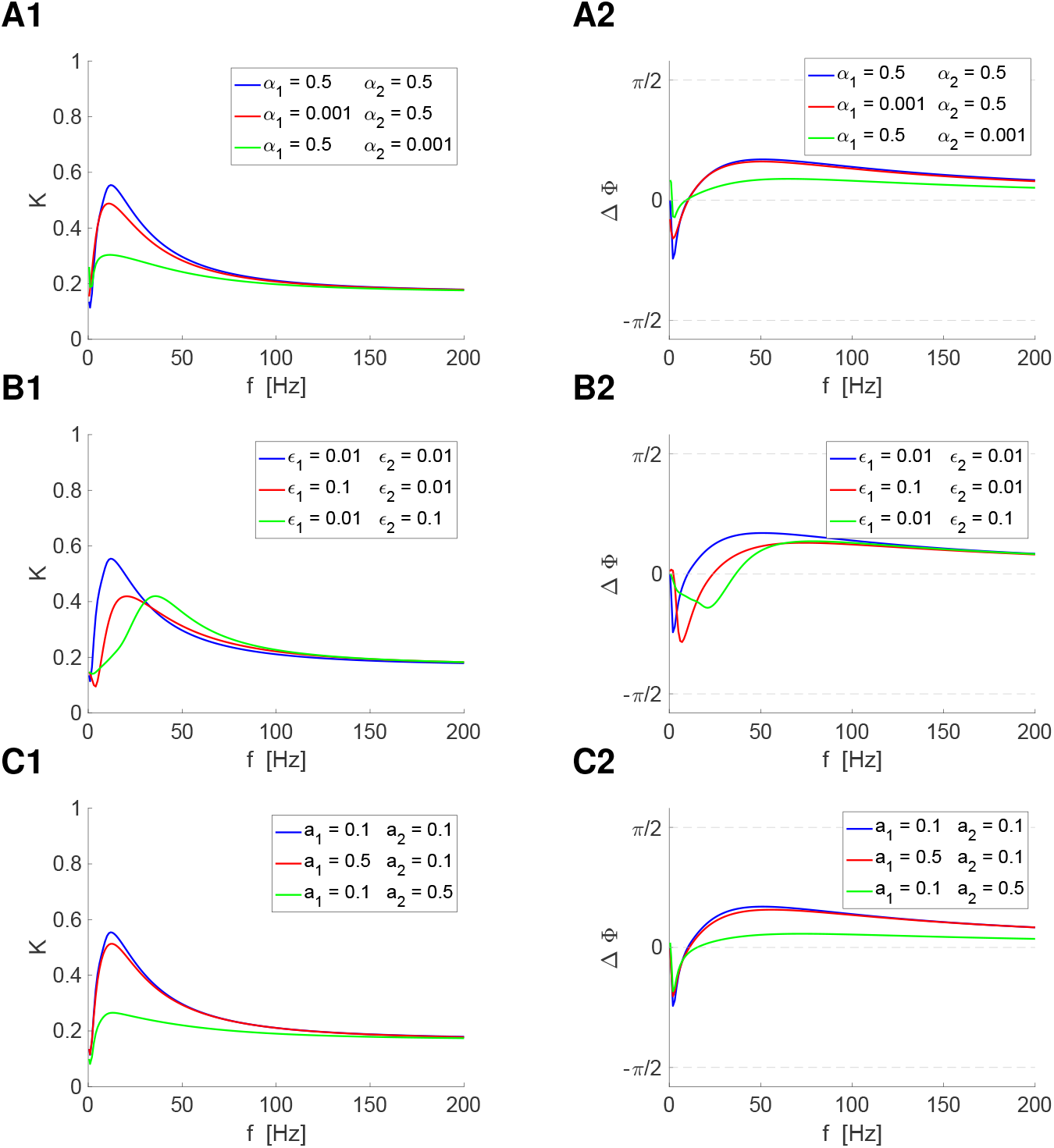
Electrically coupled quadratic 2D model: Effects of the pre-J and post-J cells’ properties on the response profiles. We used the quadratic model (3)-(4) for the electrically coupled pre- and post-J cells with quadratic nonlinearities mediated by gap junctions (cell 1 is the pre-J cell; *A*_*in,2*_ = 0). The parameter values for the two cells are identical, except for the ones indicated in the legends. **Left column.** *K* profiles **Right column.** ΔΦ profiles **A.** Effect of the *α* (negative feedback / resonant current strength). We used *a*_1_ = *a*_2_ = 0.1, *ϵ*_1_ = *ϵ*_2_ = 0.01. **A-blue.** The pre-J and post-J cells are identical BPFs (Fig. 14-A). **A-red.** The pre-J cell is a LPF (Fig. 14-C) and the post-J cell is a BPF (Fig. 14-A). **A-green.** The pre-J cell is a BPF (Fig. 14-A) and the post-J cell is a LPF (Fig. 14-C). **B.** Effect of *ϵ* (negative feedback / resonant current time constant). We used *a*_1_ = *a*_2_ = 0.1, *α*_1_ = *α*_2_ = 0.5. The pre-J and post-J cells are both BPFs (Figs. 14-A and -D). **B-blue.** The pre-J and post-J cells are identical BPFs (Fig. 14-A). **B-red.** The resonant frequency of the pre-J cell (Fig. 14-D) is higher than for the post-J cell (Fig. 14-A). **B-green.** The resonant frequency of the pre-J cell (Fig. 14-A) is lower than for the post-J cell (Fig. 14-D). **C.** Effect of *a* (curvature of the parabolic nonlinearity). We used *α*_1_ = *α*_2_ = 0.5, *ϵ*_1_ = *ϵ*_2_ = 0.01. **C-blue.** The *V* -nullcline for the pre-J and post-J cells have identical curvatures (Fig. 14-A). **C-red.** The curvature of the *V* -nullcline for the pre-J cell is larger (Fig. 14-B) than for the post-J cell (Fig. 14-A). **C-green.** The curvature of the *V* -nullcline for the pre-J cell is smaller (Fig. 14-A) than for the post-J cell (Fig. 14-B). We used the following additional parameter values:, *λ*_1_ = *λ*_2_ = *−*0.2, *g*_*c*_ = 0.1, *A*_*in,1*_ = 0.06 and *A*_*in,2*_ = 0.

**Figure 17:**
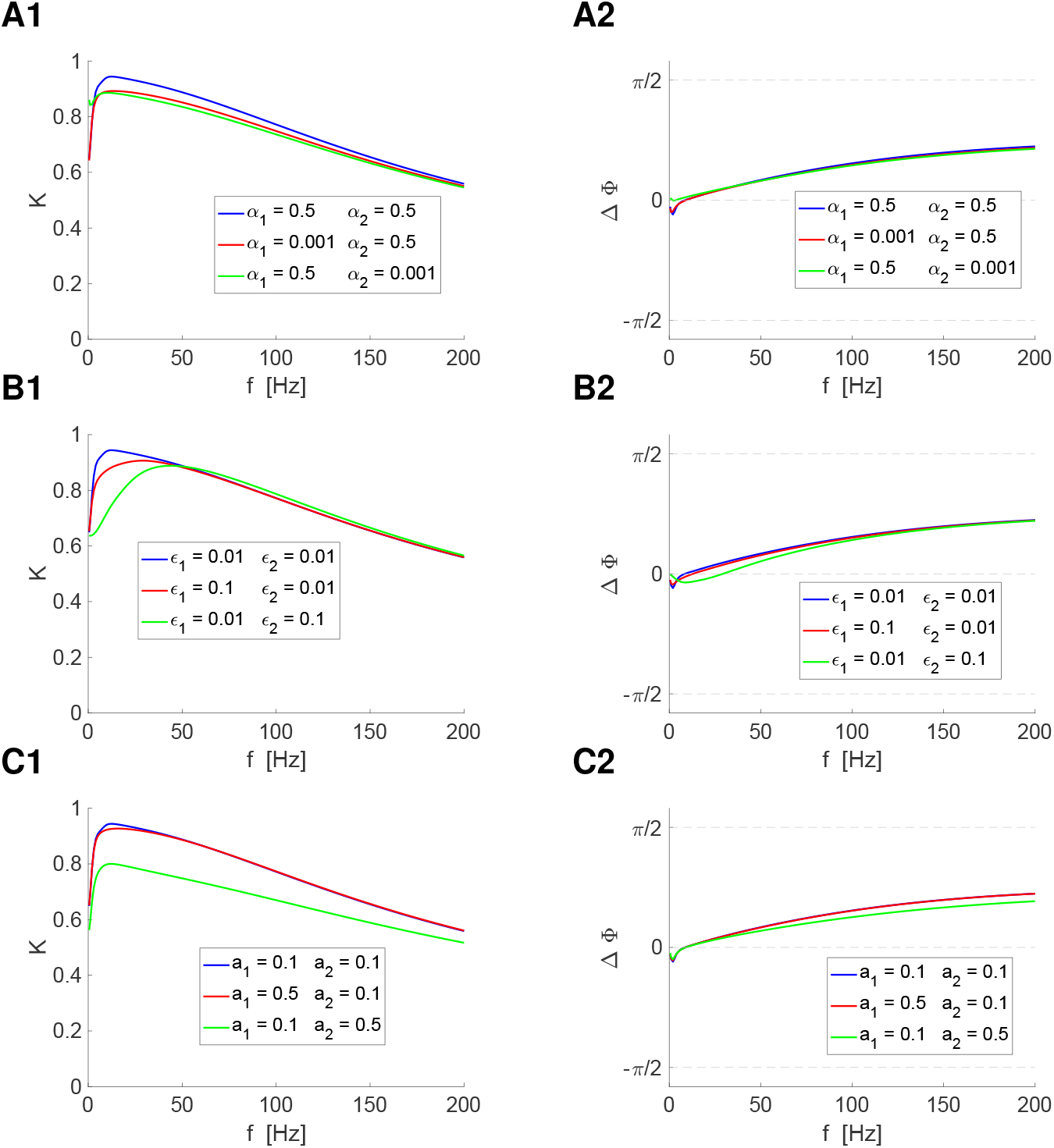
Electrically coupled quadratic 2D model: Effects of the pre-J and post-J cells’ properties on the response profiles. We used the quadratic model (3)-(4) for the electrically coupled pre- and post-J cells with quadratic nonlinearities mediated by gap junctions (cell 1 is the pre-J cell; *A*_*in,2*_ = 0). The parameter values for the two cells are identical, except for the ones indicated in the legends. **Left column.** *K* profiles **Right column.** ΔΦ profiles **A.** Effect of the *α* (negative feedback / resonant current strength). We used *a*_1_ = *a*_2_ = 0.1, *ϵ*_1_ = *ϵ*_2_ = 0.01. **A-blue.** The pre-J and post-J cells are identical BPFs (Fig. 14-A). **A-red.** The pre-J cell is a LPF (Fig. 14-C) and the post-J cell is a BPF (Fig. 14-A). **A-green.** The pre-J cell is a BPF (Fig. 14-A) and the post-J cell is a LPF (Fig. 14-C). **B.** Effect of *ϵ* (negative feedback / resonant current time constant). We used *a*_1_ = *a*_2_ = 0.1, *α*_1_ = *α*_2_ = 0.5. The pre-J and post-J cells are both BPFs (Figs. 14-A and -D). **B-blue.** The pre-J and post-J cells are identical BPFs (Fig. 14-A). **B-red.** The resonant frequency of the pre-J cell (Fig. 14-D) is higher than for the post-J cell (Fig. 14-A). **B-green.** The resonant frequency of the pre-J cell (Fig. 14-A) is lower than for the post-J cell (Fig. 14-D). **C.** Effect of *a* (curvature of the parabolic nonlinearity). We used *α*_1_ = *α*_2_ = 0.5, *ϵ*_1_ = *ϵ*_2_ = 0.01. **C-blue.** The *V* -nullcline for the pre-J and post-J cells have identical curvatures (Fig. 14-A). **C-red.** The curvature of the *V* -nullcline for the pre-J cell is larger (Fig. 14-B) than for the post-J cell (Fig. 14-A). **C-green.** The curvature of the *V* -nullcline for the pre-J cell is smaller (Fig. 14-A) than for the post-J cell (Fig. 14-B). We used the following additional parameter values:, *λ*_1_ = *λ*_2_ = *−*0.2, *g*_*c*_ = 1, *A*_*in,1*_ = 0.08 and *A*_*in,2*_ = 0.

**Figure 18:**
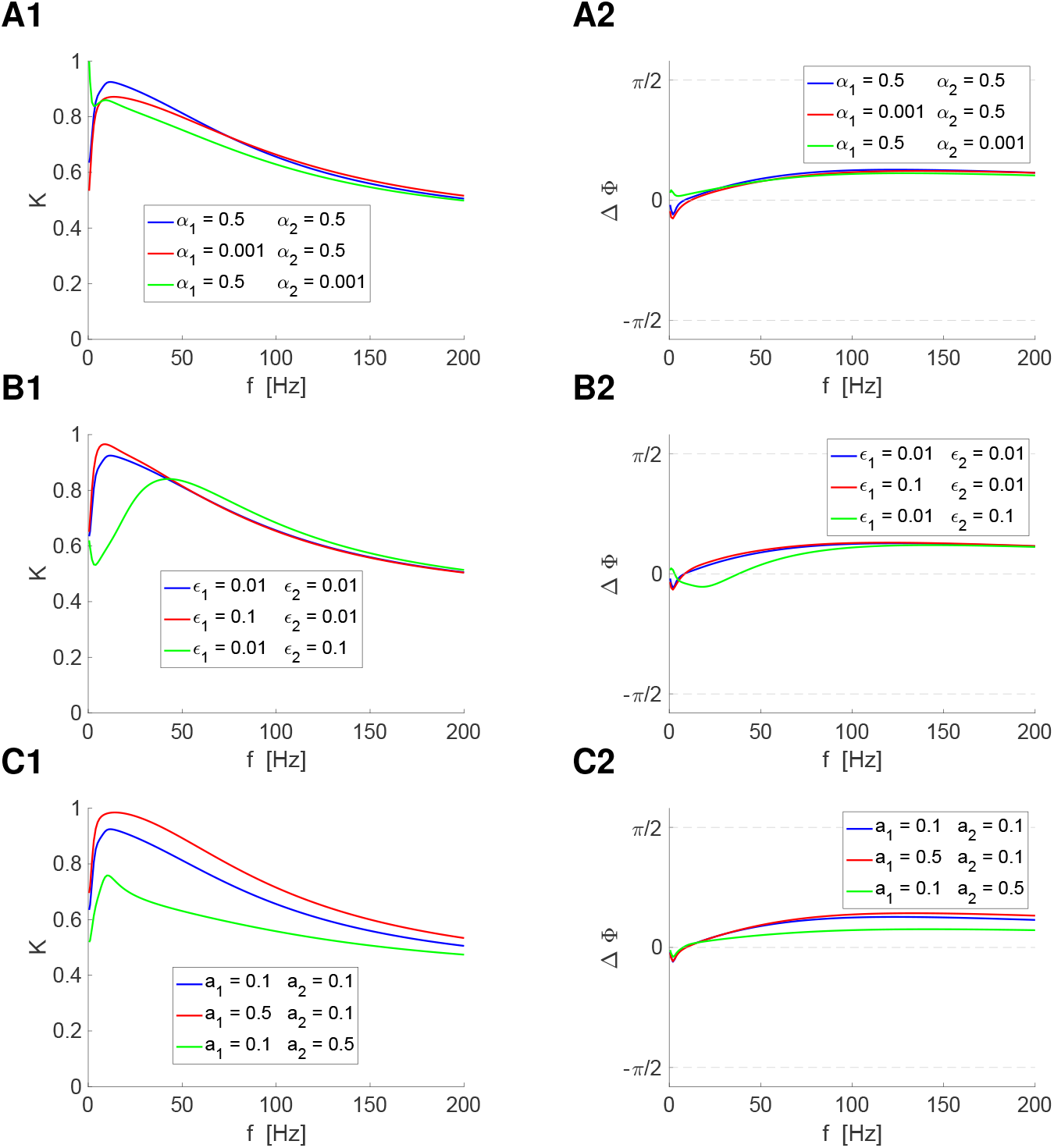
Two-compartment model with quadratic 2D dynamics : Effects of the pre-J and post-J compartments’ properties on the response profiles. We used the quadratic model (3)-(4) for the electrically coupled pre- and post-J compartments with quadratic nonlinearities where the coupling parameter *g*_*c,k*_ was substituted by *g*_*c*_/σ_*k*_ with *σ*_1_ = 0.8 and *σ*_2_ = 0.2 (compartment 1 is the pre-J compartment; *A*_*in,2*_ = 0). The parameter values for the two compartments are identical, except for the ones indicated in the legends. **Left column.** *K* profiles **Right column.** ΔΦ profiles **A.** Effect of the *α* (negative feedback / resonant current strength). We used *a*_1_ = *a*_2_ = 0.1, *ϵ*_1_ = *ϵ*_2_ = 0.01. **A-blue.** The pre-J and post-J compartments are identical BPFs (Fig. 14-A). **A-red.** The pre-J compartment is a LPF (Fig. 14-C) and the post-J compartment is a BPF (Fig. 14-A). **A-green.** The pre-J compartment is a BPF (Fig. 14-A) and the post-J compartment is a LPF (Fig. 14-C). **B.** Effect of *ϵ* (negative feedback / resonant current time constant). We used *a*_1_ = *a*_2_ = 0.1, *α*_1_ = *α*_2_ = 0.5. The pre-J and post-J compartments are both BPFs (Figs. 14-A and -D). **B-blue.** The pre-J and post-J compartments are identical BPFs (Fig. 14-A). **B-red.** The resonant frequency of the pre-J compartment (Fig. 14-D) is higher than for the post-J compartment (Fig. 14-A). **B-green.** The resonant frequency of the pre-J compartment (Fig. 14-A) is lower than for the post-J compartment (Fig. 14-D). **C.** Effect of *a* (curvature of the parabolic nonlinearity). We used *α*_1_ = *α*_2_ = 0.5, *ϵ*_1_ = *ϵ*_2_ = 0.01. **C-blue.** The *V* -nullcline for the pre-J and post-J cells have identical curvatures (Fig. 14-A). **C-red.** The curvature of the *V* -nullcline for the pre-J cell is larger (Fig. 14-B) than for the post-J cell (Fig. 14-A). **C-green.** The curvature of the *V* -nullcline for the pre-J cell is smaller (Fig. 14-A) than for the post-J cell (Fig. 14-B). We used the following additional parameter values:, *λ*_1_ = *λ*_2_ = *−*0.2, *g*_*c*_ = 0.05, *A*_*in,1*_ = 0.06 and *A*_*in,2*_ = 0.

**Figure 19:**
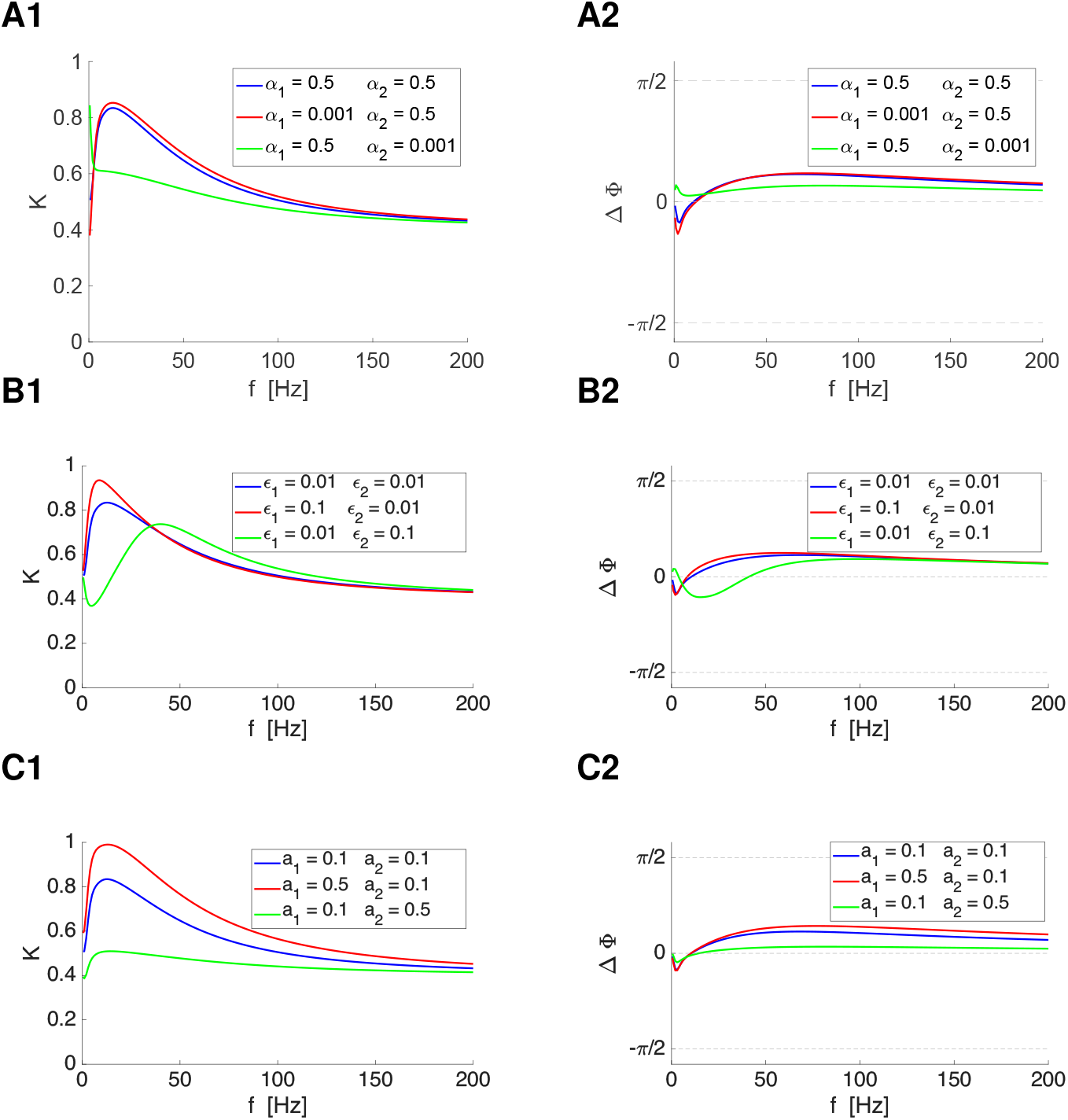
Two-compartment model with quadratic 2D dynamics : Effects of the pre-J and post-J compartments’ properties on the response profiles. We used the quadratic model (3)-(4) for the electrically coupled pre- and post-J compartments with quadratic nonlinearities where the coupling parameter *g*_*c,k*_ was substituted by *g*_*c*_/σ_*k*_ with *σ*_1_ = 0.2 and *σ*_2_ = 0.8 (compartment 1 is the pre-J compartment; *A*_*in,2*_ = 0). The parameter values for the two compartments are identical, except for the ones indicated in the legends. **Left column.** *K* profiles **Right column.** ΔΦ profiles **A.** Effect of the *α* (negative feedback / resonant current strength). We used *a*_1_ = *a*_2_ = 0.1, *ϵ*_1_ = *ϵ*_2_ = 0.01. **A-blue.** The pre-J and post-J compartments are identical BPFs (Fig. 14-A). **A-red.** The pre-J compartment is a LPF (Fig. 14-C) and the post-J compartment is a BPF (Fig. 14-A). **A-green.** The pre-J compartment is a BPF (Fig. 14-A) and the post-J compartment is a LPF (Fig. 14-C). **B.** Effect of *ϵ* (negative feedback / resonant current time constant). We used *a*_1_ = *a*_2_ = 0.1, *α*_1_ = *α*_2_ = 0.5. The pre-J and post-J compartments are both BPFs (Figs. 14-A and -D). **B-blue.** The pre-J and post-J compartments are identical BPFs (Fig. 14-A). **B-red.** The resonant frequency of the pre-J compartment (Fig. 14-D) is higher than for the post-J compartment (Fig. 14-A). **B-green.** The resonant frequency of the pre-J compartment (Fig. 14-A) is lower than for the post-J compartment (Fig. 14-D). **C.** Effect of *a* (curvature of the parabolic nonlinearity). We used *α*_1_ = *α*_2_ = 0.5, *ϵ*_1_ = *ϵ*_2_ = 0.01. **C-blue.** The *V* -nullcline for the pre-J and post-J cells have identical curvatures (Fig. 14-A). **C-red.** The curvature of the *V* -nullcline for the pre-J cell is larger (Fig. 14-B) than for the post-J cell (Fig. 14-A). **C-green.** The curvature of the *V* -nullcline for the pre-J cell is smaller (Fig. 14-A) than for the post-J cell (Fig. 14-B). We used the following additional parameter values:, *λ*_1_ = *λ*_2_ = *−*0.2, *g*_*c*_ = 0.05, *A*_*in,1*_ = 0.06 and *A*_*in,2*_ = 0.

For example, in Figs. 16-A, green, the post-J cell is a LPF and the Φ profile is monotonically increasing, but the *K*(*f*) and ΔΦ(*f*) profiles exhibit *K*-resonance and ΔΦ-phasonance. On the other hand, in Figs. 16-B (green) the *K*(*f*) and ΔΦ(*f*) profiles appear to be dominated by the amplitude and phase profiles of the post-J cell since they are shifted to the right as compared to the other two cases (blue and red). However, the presence of antiresonance and antiphasonance in Figs. 16-B (red) are generated, as mentioned above, by a network mechanism. Similarly, in Figs. 16-C, green, the *K*(*f*) and ΔΦ(*f*) profiles are significantly different than the other two cases (blue and red).

Comparison among Figs. 16, 18 and 19 provide a picture of how the interplay of the neuronal biophysical properties and dendritic geometry shape the *K*(*f*) and ΔΦ(*f*) profiles. To aid in the comparison, Figs. S8, S9 and S10 shows a rearrangement of the panels shown in Figs. 16, 18 and 19. Each row in Figs. S8, S9 and S10 correspond to a value of *σ* for the same fixed set of biophysical parameter values.

## 4 Discussion

The coupling strength between nodes in electrically coupled networks receiving time-dependent inputs is frequency-dependent. The *K*(*f*) and ΔΦ(*f*) coupling coefficient (CC) profiles measure the degree to which the pre-J and post-J cells are coupled in terms of their response amplitude and phase profiles, which are shaped by the complex interaction between the biophysical and geometric properties of the participating nodes and the connectivity. The biophysical and dynamic mechanisms that govern the generation of the *K*(*f)* and ΔΦ(*f*) profiles were largely unknown beyond electrically coupled passive cells (but see [32, 43, 49, 51–53]), which are 1D and linear, and for which the *K*(*f)* profile is a LPF and ΔΦ profile is always positive and monotonically increasing [23–29, 36, 40, 41, 43, 44, 46–50]. In addition, for linear systems, the CC profiles are independent of the biophysical properties of the pre-J cell and the input amplitude. Whether and under what conditions the CC profiles exhibit preferred frequencies in response to oscillatory inputs (*K*-resonance and ΔΦ phasonance) and how these profiles depend on the biophysical parameters of the pre-J and post-J cells remained elusive.

In this paper we set out to understand how CC profiles with more intricate frequency-filtering properties than these for passive cells emerge in the more complex electrically coupled networks where the pre-J and post-J nodes are higher-dimensional and nonlinear, and how these CC profiles are shaped by the biophysical properties of these nodes. 2D models incorporate the presence of cellular resonance and phasonance and the associated BPFs and mixed leading-lagging phase responses. Nonlinear models incorporate the dependence of the *K*(*f)* and ΔΦ(*f)* profiles on the properties of the pre-J cell and the input amplitude in addition to the properties of the post-J cell and the connectivity.

We used biophysical modeling, numerical simulations, analytical calculations and dynamical systems tools to address these issues. We focused on gap junction connected networks and compartmental models of spatially extended neurons, which belong to the same family of electrically coupled network. The connectivity for gap junction networks is symmetric. Asymmetric connectivity arises in two-compartment models with heterogeneous spatial geometry.

We considered minimal network architectures consisting of two electrically connected nodes receiving oscillatory inputs (Fig. 2). The individual nodes (when disconnected) were either LPFs (e.g., passive cells; Fig. 1-A1) or BPFs (e.g., resonators; Fig. 1-A2) and their phase responses were positive (e.g., passive cells; Fig. 1-C1) or mixed negative-positive (e.g., phasonators; Fig. 1-C2). These choices allowed us to systematically investigate the biophysical and dynamic mechanisms that shape the CC profiles and determine their preferred coupling frequencies (*f*_*res,K*_ and *f*_*phas,K*_) in several representative scenarios where the same or different types of cellular filters interact. While gap junction connected networks typically involve the same type of pre-J and post-J neuron with the same biophysical properties, compartmental models may involve compartments with heterogeneity in their intrinsic properties [55, 92].

Linear models are amenable to analytical calculations, which we used to understand how the *K*(*f)* and ΔΦ(*f)* profiles are shaped by the participating building blocks (e.g., ionic conductances controlling the negative and positive feedback effects) and cellular filtering properties. The *K*(*f)* and ΔΦ(*f)* profiles for passive cells transition to BPFs and mixed negative-positive, respectively, in the presence of an added negative feedback term, which transform the nodes into resonators (2D). The shape of these CC profiles are modulated by the linearized leak and negative feedback conductances and the negative feedback time constant, which control the corresponding filtering properties of the post-J cell.

Armed with these results and knowledge, we investigated the dynamics of nonlinear neurons with biophysically realistic ionic currents to understand, to the linear level of approximation, how the presence and different types of amplifying and resonant ionic processes shape the CC profiles. Motivated by experimental results and previous theoretical studies, we used the *I*_*Nap*_ + *I*_*h*_ and *I*_*Nap*_ + *I*_*Ks*_ models [21, 55, 66, 67] that exhibit both cellular resonance and phasonance [63–65]. These two models are biophysically different, because they involve different resonant currents, but dynamically similar since their phase-plane diagrams are almost a mirror image of each other [93].

Each parameter in the linearized models is a function of a set of parameters in the (original) biophysical model. The response for each set of parameter values in the linearized model is, obviously, linear. However, the notion of linearity has different meanings and implications depending on the context. On the one hand, linear models are reduced models. In this sense, the response of a linearized models are approximation to the original corresponding biophysical model. On the other hand, linear models have certain properties that become apparent only upon changes in the model parameters (e.g., the amplitudes of the response is proportional to the input for all frequencies). Here, we were interested in processes such as amplification and the modulation of resonance, which materialize when one changes the amplifying and resonant ionic conductances.

In the “linear study” discussed above, we investigated how the *K*(*f)* and ΔΦ(*f)* profiles change as we changed the linear parameters independently of any other consideration. In this “linearized” part of the study, in contrast, we changed the biophysical parameters (e.g., ionic conductances) and recalculated the linear parameters for each one of the choices of biophysical parameter values. This process captured the nonlinear effects “to the linear level of approximation” and the differences between the way the two models (*I*_*Nap*_ + *I*_*h*_ and *I*_*Nap*_ + *I*_*Ks*_) shape the *K*(*f)* and Δ(*f)* profiles and determine their preferred frequencies. This is summarized in Tables 2 and 3. One salient difference between the results for the two models is that, in the presence of high enough levels of *I*_*Nap*_, increasing levels of *I*_*h*_ amplify the *K*(*f)* profile, while increasing values of *I*_*Ks*_ attenuate the *K*(*f)* profile, while in the presence of low enough levels of *I*_*Nap*_, increasing values of both *I*_*h*_ and *I*_*Ks*_ attenuate the *K*(*f)* profile. A similar effect has been observed for the impedance profile in single cells [21]. The effects of asymmetry in the connectivity (e.g., for two-compartment models) was inferred from these results by a set of algebraic operations.

While the linearized models capture certain nonlinear properties of the behavior of the biophysical models, the linearized models’ CC profiles remain independent of the properties of the pre-J cells and depend only on the properties of the post-J cell and the connectivity. A set of analytical results using regular perturbation theory for electrically coupled weakly nonlinear cells showed that the both the pre-J and post-J cells contribute to shaping the CC profiles. However, while weakly nonlinear models capture the dynamics of the *I*_*Nap*_ + *I*_*h*_ and *I*_*Nap*_ + *I*_*Ks*_ models for at most low levels of *I*_*Nap*_, they fail to do so for the more realistic scenarios with higher levels of *I*_*Nap*_ causing the development of voltage nonlinearities of parabolic type in a vicinity of the resting potential. To understand how these nonlinearities affect the properties of the CC profiles, we extended our investigation to networks of electrically coupled quadratized cells, which have a parabolic voltage nonlinearities and linear dynamics for the recovery variable. Similarly to linearization, quadratization of biophysical models produces a quadratic model whose parameters are functions of the biophysical parameters of the original, parabolic-like model. In this case, the *I*_*Nap*_ + *I*_*h*_ and *I*_*Nap*_ + *I*_*Ks*_ models.

We computationally analyzed the biophysical mechanisms of generation of the *K*(*f)* and Δ(*f)* in the quadratic models and demonstrated computationally that they depend on the properties of both the pre-J and post-J cells and the network connectivity. In some cases, the contribution of the pre-J cells is relatively small, but in others is significant, which is missed by using linearized models. Finally, we showed that these results persists for two-compartment models with the additional dependence on the pre-J and post-J geometric properties and for inputs effectively arriving in different compartments.

Our results advance our knowledge on the response of electrically coupled networks to oscillatory inputs and the frequency-dependent properties of the CC profiles. These results highlight the emergence of preferred CC frequencies at which *K*(*f)* peaks (the coupling strength is maximal at the *K*-resonant frequency *f*_*res,K*_) and for which the pre-J and post-J cell peak synchronously in phase (ΔΦ = 0 at the ΔΦ-phasonant frequency *f*_*phas,K*_). These results have direct implications for spike transmission [32, 52] in electrically coupled networks in response to oscillatory inputs, for the propagation of signals along the somato-dendritic axis in the presence of resonance and phasonance along the dendritic tree [54, 55, 94], for the rhythmic coding of hippocampal pyramidal cells [95] and, more generally, for dendritic computation [96–101].

The ideas and results developed in this paper can be extended to electrically coupled networks with a larger number of nodes and possibly multiple inputs, to spatially extended multicompartment neuronal models possibly receiving inputs at different locations (e.g., distal and proximal inhibition by oriens lacunosum-moleculare and fast spiking interneurons, respectively, in hippocampal pyramidal cells), to models having combined amplifying and resonant processes such as *I*_*Ca,T*_ activation and inactivation, respectively, and to *I*_*Nap*_ + *I*_*h*_ and *I*_*Nap*_ + *I*_*Ks*_ models having nonlinearities of cubic type, instead of parabolic type, in the subthreshold voltage regime [93]. More research is needed to address these issues. Research is also needed to test experimentally the predictions of our study.

## Acknowledgments

AB and UC acknowledge support from the Universidad Nacional del Sur grant PGI 24/L131 and CONICET, Argentina. HGR acknowledges support from the National Science Foundation grants DMS-1608077 and IOS-2002863. HGR is a Corresponding Researcher in CONICET, Argentina and a Graduate Faculty Member in the Graduate Program in Neuroscience (GPN) in the Center for Molecular and Behavioral Neuroscience (CMBN) at Rutgers University.

## Supplementary Material

**S1.** Linearization and quadratization of conductance-based (biophysically plausible) models of Hodgkin-Huxley type

**S2.** Impedance amplitude and phase profiles for 2D linear systems: individual cells

**S3.** Network impedance profiles for linear systems: networks with linear nodes and connectivity

**S4.** Networks of electrically coupled weakly nonlinear (quasi-linear) cells: Stationary and frequency-dependent coupling coefficient

**S5.** Impedance amplitude and phase profiles for 2D linear systems: general calculation

**S6.** Additional Figures.

Part of the supplementary material has been used in other publications (e.g., [102, 103])

### S1 Linearization and quadratization of conductance - based (biophysically plausible) models of Hodgkin-Huxley type

#### S1.1 Conductance-based models of single cells

We use the following biophysical (conductance-based) models of Hodgkin-Huxley type [69,73] to describe the neuronal subthreshold dynamics

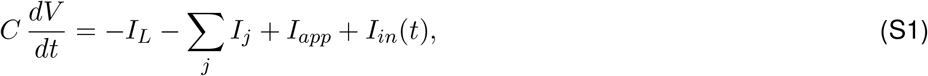

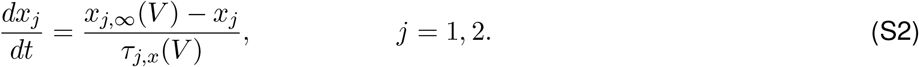

In the current-balance equation (S1), *V* is the membrane potential (mV), *t* is time (ms), *C* is the membrane capacitance (*µ*F/cm^2^), *I*_*app*_ is the applied bias (DC) current (*µ*A/cm^2^), *I*_*L*_ = *G*_*L*_ (*V ™ E*_*L*_) is the leak current, and *I*_*j*_ = *G*_*j*_ *x*_j_, (*V ™ E*_*j*_) are generic expressions for ionic currents (with *j* an index) with maximal conductance *G*_*j*_ (mS/cm^2^) and reversal potentials *E*_*j*_ (mV) respectively. The dynamics of the gating variables *x*_*j*_ obey the kinetic equations (S2) where *x*_*j,∞*_(*V)* and *τ*_*j,x*_(*V)* are the voltage-dependent activation/inactivation curves and time constants respectively. For simplicity, the generic ionic currents *I*_*j*_ we consider here are restricted to have a single gating variable *x*_*j*_ and to be linear in *x*_*j*_. This is typically the case for persistent sodium (*I*_*Nap*_), *h*-(hyperpolarization-activated, mixed-cation, inward; *I*_*h*_), and slow-potassium (M-type) (*I*_*Ks*_) currents (see Section S1.2). Our discussion and results can be easily adapted and generalized to include ionic currents having two gating variables [20, 87–89] raised to powers different from one such as T-type calcium and A-type potassium currents [1]. The input current *I*_*in*_(*t*) (*µ*A/cm^2^) in eq. (S1) has the form (5), *I*_*in*_(*t*) = *A*_*in*_ *sin*(Ω *t*) with Ω = 2 *π f /* 1000 (*f* is the input frequency in Hz).

Here we focus on 2D models describing the dynamics of *V* and the gating variable (*x*_1_) associated to the ionic current *I*_1_. Additional currents whose gating variables evolve on a very fast time scale (as compared to the other variables) can be included by using the adiabatic approximation *x*_*j*_ = *x*_*j,∞*_(*V)*. Here we include one such fast current *I*_2_ = *G*_2_ *x*_2,*∞*_(*V)* (*V ™ E*_2_). Additional fast currents can be included without significantly changing the formalism used here.

#### S1.2 Two examples: *I*_*Nap*_ + *I*_*h*_ and *I*_*Nap*_ + *I*_*Ks*_ models in the subthreshold regime

The *I*_*Nap*_ + *I*_*h*_ and *I*_*Nap*_ + *I*_*Ks*_ models [66, 71] are adapted versions of the models presented in [75, 78] (see also [66,67,70–72]). The *I*_*Nap*_ + *I*_*h*_ involves the interaction between a persistent sodium (*I*_*Nap*_ = *I*_2_) and a hyperpolarization -activated (or h-) (*I*_*h*_ = *I*_1_) currents. The *I*_*Nap*_ + *I*_*Ks*_ model involves the interaction between a persistent sodium (*I*_*Nap*_ = *I*_2_) and an M-type slow potassium (*I*_*Ks*_ = *I*_1_) currents. The formulation of these currents is

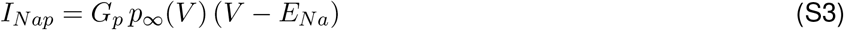

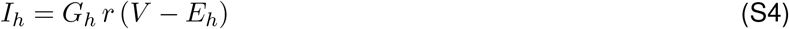

and

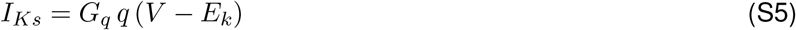

The activation and inactivation curves we used for these currents are given, respectively, by the following functions

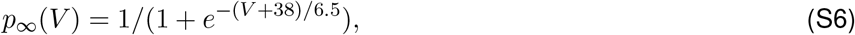

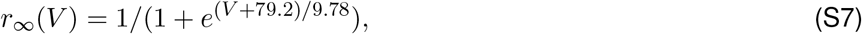

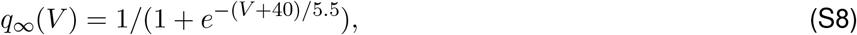

that have the general form

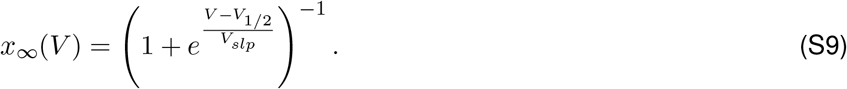

Graphs of these functions are shown in Fig. S1 (see also [70], Fig. 1)).

**Figure S1:**
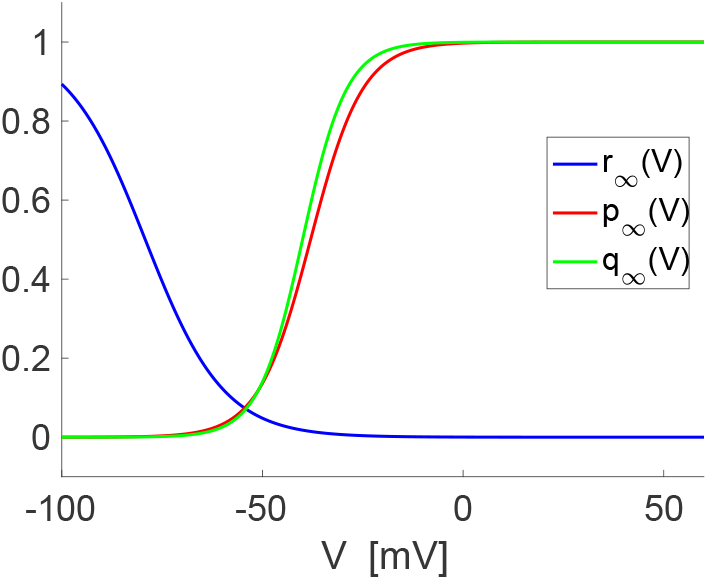
**Activation and inactivation curves for the *I*_*Nap*_ + *I*_*h*_ and *I*_*Nap*_ + *I*_*Ks*_ models. We used eqs. (S6)-(S8) in Section S1.2**.

#### S1.2.1 Parameter values for the I_Nap_ + I_h_ model

We used the following parameter values unless stated otherwise: *C* = 1, *E*_*L*_ = *−*65, *E*_*h*_ = *−*20, *E*_*Na*_ = 55, *G*_*L*_ = 0.5, *V*_*p,1/2*_ = *−*38, *V*_*p,slp*_ = *−*65, *V*_*r,1/2*_ = *−*79.2, *V*_*r,slp*_ = 9.78, *τ*_*r*_ = 100, *G*_*p*_ = 0.5, *G*_*h*_ = 1.5, *I*_*app*_ = *−*2.5. The values of *G*_*L*_, *G*_*p*_, *G*_*h*_ and *I*_*app*_ are the baseline parameter values and were varied in our simulations.

For our calculations using the generic model (S1)-(S2), we used the following notation: *E*_1_ = *E*_*h*_, *E*_2_ = *E*_*Na*_, *G*_1_ = *G*_*h*_, *G*_2_ = *G*_*p*_, *V*_1,1*/*2_ = *V*_*h,1/2*_, *V*_1,*slp*_ = *V*_*h,slp*_, *V*_2,1*/*2_ = *V*_*p,1/2*_, *V*_2,*slp*_ = *V*_*p,slp*_, *τ*_1_ = *τ*_*r*_.

#### S1.2.2 Parameter values for the I_Nap_ + I_Ks_ model

We used the following parameter values: *C* = 1, *E*_*L*_ = *−*54, *E*_*K*_ = *−*90, *E*_*Na*_ = 55, *G*_*L*_ = 0.1, *V*_*p,1/2*_ = *−*38, *V*_*p,slp*_ = −65, *V*_*q,1/2*_ = −40, *V*_*q,slp*_ = −5.5, *τ*_*q*_ = 100, *G*_*p*_ = 0.21, *G*_*q*_ = 1, *I*_*app*_ = −0.6. The values of *G*_*L*_, *G*_*p*_, *G*_*q*_ and *I*_*app*_ are the baseline parameter values and were varied in our simulations.

For our calculations using the generic model (S1)-(S2), we used the following notation: *E*_1_ = *E*_*K*_, *E*_2_ = *E*_*Na*_, *G*_1_ = *G*_*q*_, *G*_2_ = *G*_*p*_, *V*_1,1*/*2_ = *V*_*q,1/2*_, *V*_1,*slp*_ = *V*_*q,slp*_, *V*_2,1*/*2_ = *V*_*p,1/2*_, *V*_2,*slp*_ = *V*_*p,slp*_, *τ*_1_ = *τ*_*q*_ .

#### S1.2.3 Parameter values for the I_Nap_ model

We used the following parameter values: *C* = 1, *E*_*L*_ = −65, *E*_*Na*_ = 55, *G*_*L*_ = 0.1, *V*_*p,1/2*_ = −38, *V*_*p,slp*_ = −65, *G*_*p*_ = 0.1, *I*_*app*_ = 0. The values of *G*_*L*_, *G*_*p*_, and *I*_*app*_ are the baseline parameter values and were varied in our simulations. Fig. S2 illustrates the dynamics for representative scenarios.

For our calculations using the generic model (S1)-(S2), we used the following notation: *E*_2_ = *E*_*Na*_, *G*_2_ = *G*_*p*_, *V*_2,1*/*2_ = *V*_*p,1/2*_ and *V*_2,*slp*_ = *V*_*p,slp*_.

**Figure S2:**
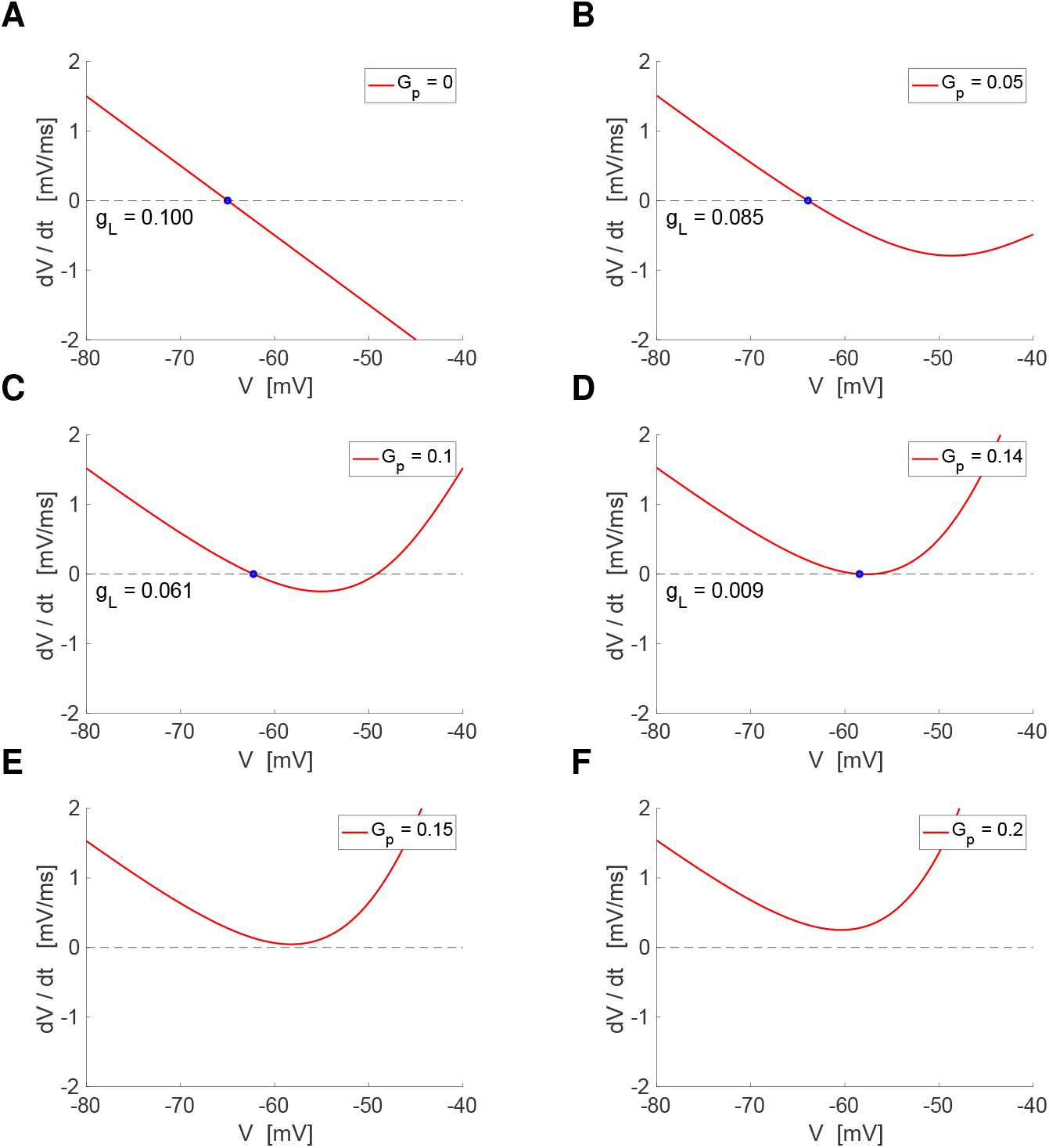
Phase-space diagrams for the *I*_*Nap*_ model (subthreshold regime) for representative parameter values. *dV/dt* vs. *V* curves for representative parameter values of *G*_*p*_ and *G*_*L*_ and *I*_*app*_ = 0. We used the parameter values presented in Section 4. This model can be thought of as a 1D reduced version of the 2D models illustrated in Fig. S6. See additional details there. The trajectories (not shown) move along the *V* axis and converge to the stable fixed points (blue dots) (A - D) or escape the subthreshold regime (E, F). For *G*_*p*_ = 0 the modes is a passive cell, which is linear (A). As *G*_*p*_ increases, nonlinearities of parabolic type develop and become more pronounced in a vicinity of the stable fixed-point (B-D). The unstable fixed-point serves as a threshold for spike generation. If the model is supplemented with a return mechanism to the subthreshold regime, it becomes a model of quadratic integrate-and-fire type [72]. For the saddle-node bifurcation value of *G*_*p*_, the fixed-points cease to exist (E, F). The associated quadratic integrate-and-fire model produces periodic spiking.

**Figure S3:**
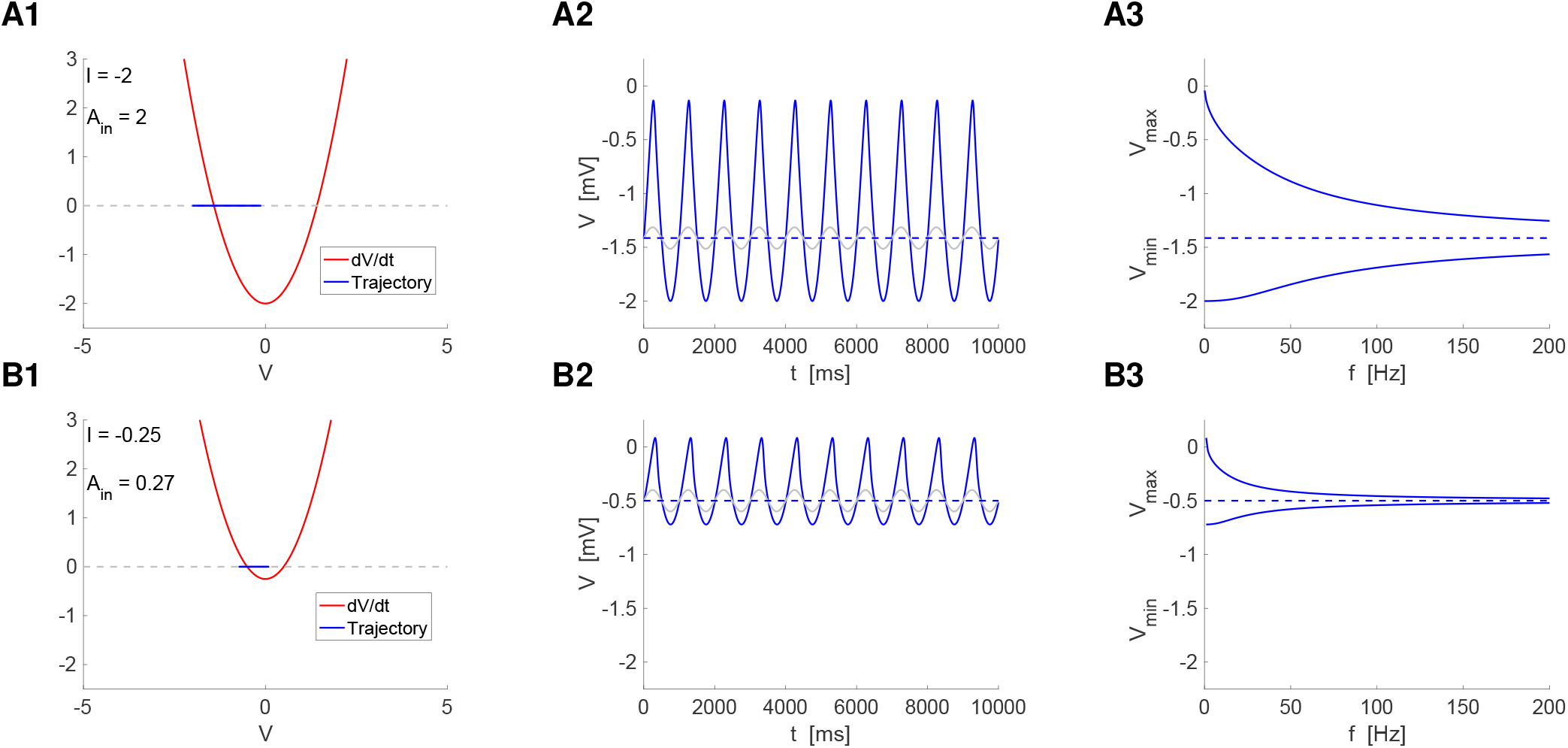
Quadratic 1D model (subthreshold regime) for representative parameter values. We used the model given by eq. (31) with *a* = 1, *τ* = 10 and *g*_*c*_ = 0 (isolated cell). **Left column.** Phase-space diagrams: *dV/dt* vs. *V* curves (red) and voltage trajectory corresponding to the voltage trace in the middle column (blue). **Middle column.** Voltage trace (blue) in response to an oscillatory input of frequency equal to 1 and amplitude *A*_*in*_ (gray; the input signal is rescaled). The value of *A*_*in*_ is the threshold value above which *v* increases unboundedly (escapes the subthreshold regime). **Right column.** Voltage envelope profile in response to oscillatory inputs with the same amplitude as in the left and middle panels.

**Figure S4:**
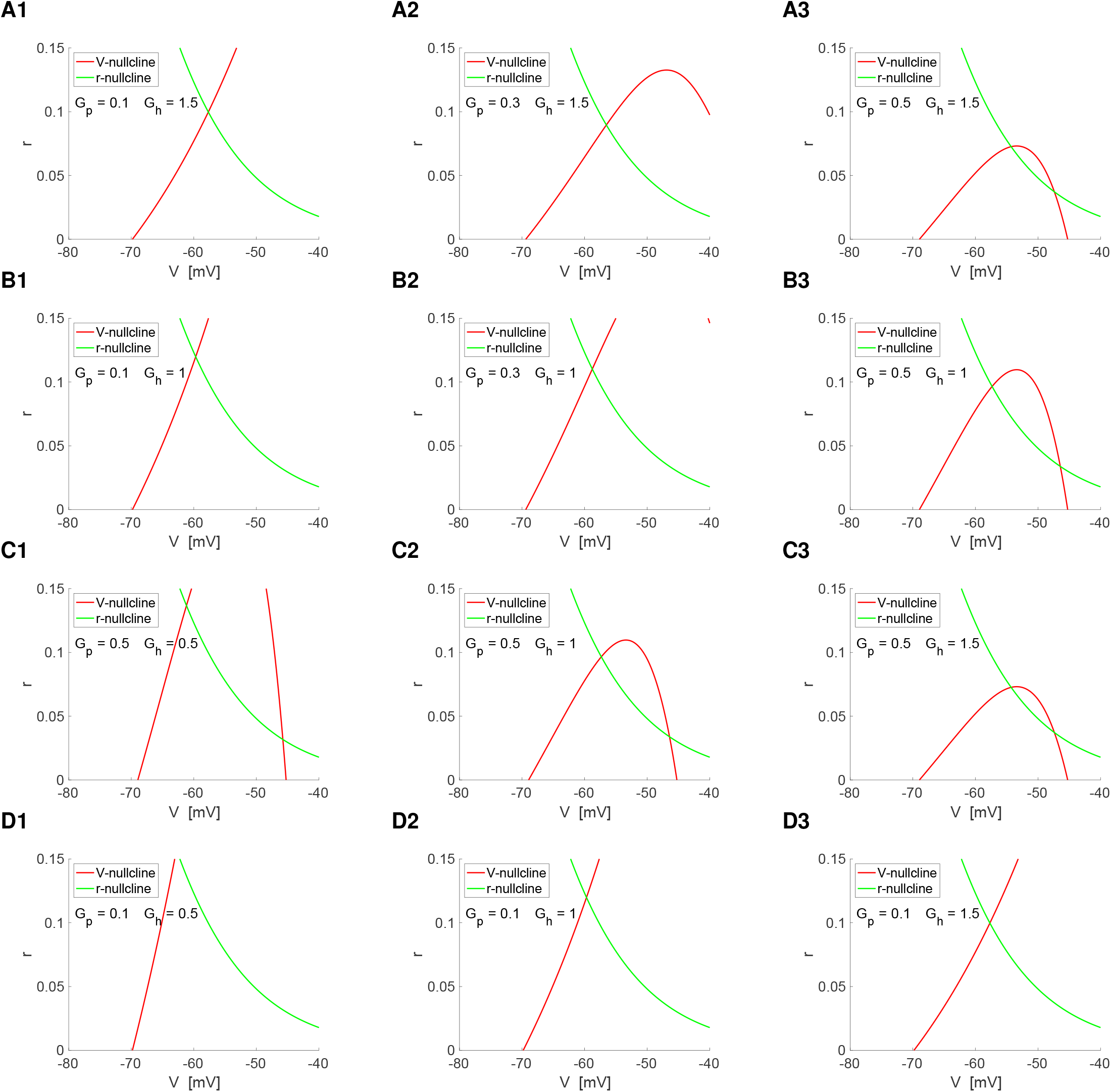
Phase-space diagram for the I_Nap_+I_h_ model (subthreshold regime) for representative parameter values of the I_Nap_ and I_h_ maximal conductances. The *V* - and *r*-nullclines were computed from eqs. (S1)-(S2) for the parameter values showed in each panel, *G*_*L*_ = 0.5, *I*_*app*_ = *−*2.5 and the remaining parameter values presented in Section S1.2. When there is a single fixed-point, it is stable. When there are two fixed-points, the one with the smaller value of *V* (left) is stable and the one with the larger values of *V* (right) is unstable. Trajectories (not shown) evolve towards the stable fixed-point (e.g., Fig. 3). **A.** Increasing values of *G*_*p*_ and *G*_*h*_ = 1.5. **B.** Increasing values of *G*_*p*_ and *G*_*h*_ = 1. **C.** Increasing values of *G*_*h*_ and *G*_*p*_ = 0.5. **C.** Increasing values of *G*_*h*_ and *G*_*p*_ = 0.1. **A, B, C.** The system transitions from quasi-linear (in a vicinity of the stable fixed-point) to parabolic-like. Increasing values of certain parameters (e.g., *I*_*app*_, *G*_*h*_, *G*_*p*_) cause Hopf bifurcation where the fixed-point loses stability and moves to the “right branch” of the parabolic-like nullcline. Trajectories leaving the subthreshold regime is interpreted as the onset of a spike and if the model is supplemented with a return mechanism to the subthreshold regime, it becomes a model of quadratic integrate-and-fire type with adaptation [72, 74, 76]. **D.** The system remains quasi-linear.

**Figure S5:**
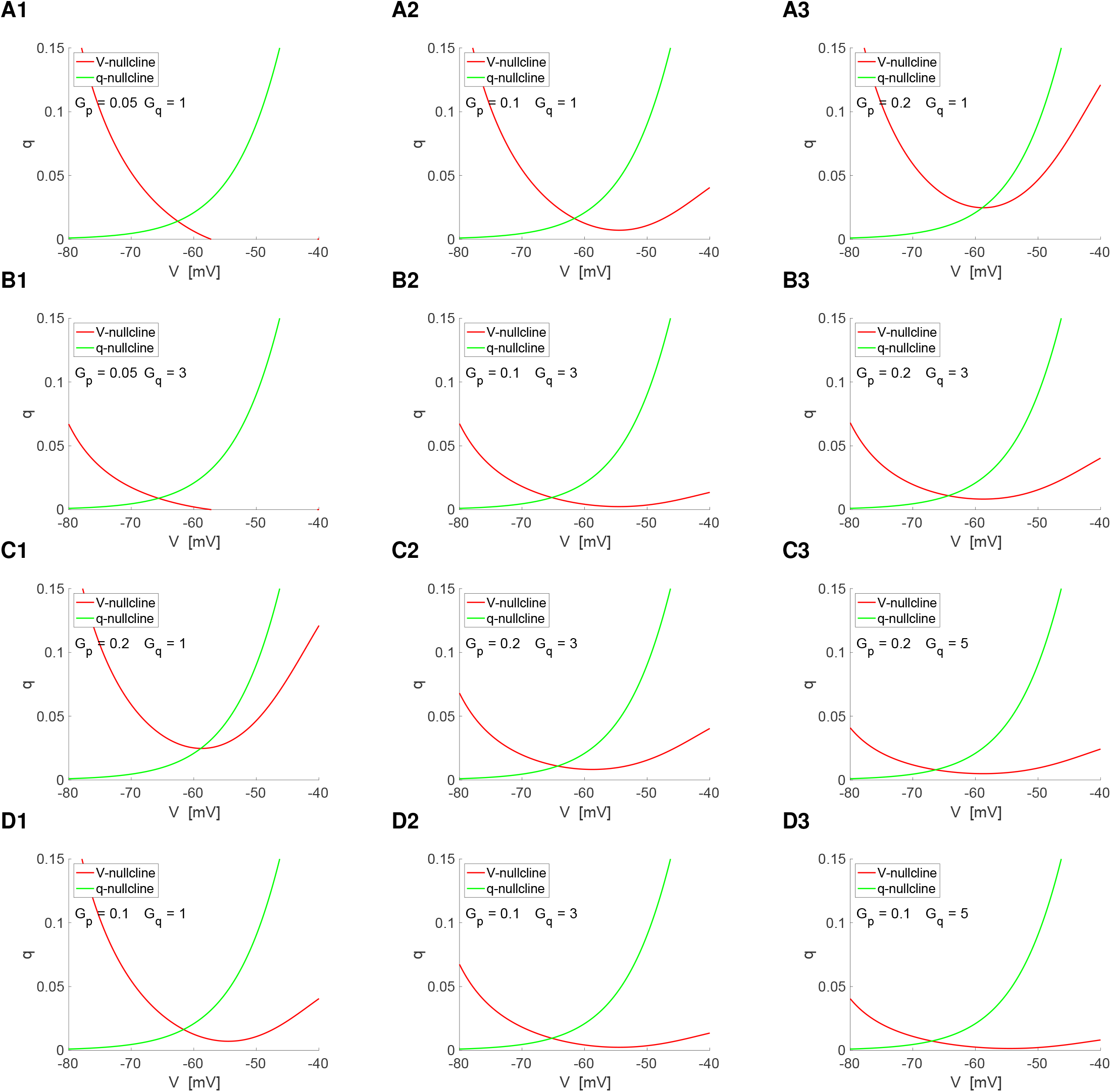
Phase-space diagram for the I_Nap_+I_Ks_ model (subthreshold regime) for representative parameter values of the I_Nap_ and I_Ks_ maximal conductances. The *V* - and *q*-nullclines were computed from eqs. (S1)-(S2) for the parameter values showed in each panel, *G*_*L*_ = 0.5, *I*_*app*_ = *−*2.5 and the remaining parameter values presented in Section S1.2. When there is a single fixed-point, it is stable. When there are two fixed-points, the one with the smaller value of *V* (left) is stable and the one with the larger values of *V* (right) is unstable. Trajectories (not shown) evolve towards the stable fixed-point (e.g., Fig. 3). **A.** Increasing values of *G*_*p*_ and *G*_*q*_ = 1. **B.** Increasing values of *G*_*p*_ and *G*_*q*_ = 3. **C.** Increasing values of *G*_*q*_ and *G*_*p*_ = 0.2. **C.** Increasing values of *G*_*q*_ and *G*_*p*_ = 0.1. **A, B.** The system transitions from quasi-linear (in a vicinity of the stable fixed-point) to parabolic-like. Increasing values of certain parameters (e.g., *I*_*app*_, *G*_*q*_, *G*_*p*_) causes Hopf bifurcation where the fixed-point loses stability and moves to the “right branch” of the parabolic-like nullcline. Trajectories leaving the subthreshold regime is interpreted as the onset of a spike and if the model is supplemented with a return mechanism to the subthreshold regime, it becomes a model of quadratic integrate-and-fire type with adaptation [72, 74, 76]. **C, D.** The system transitions from parabolic-like to quasi-linear (in a vicinity of the stable fixed-point).

#### S1.2.4 Dynamic structure of the I_Nap_ + I_h_ and I_Nap_ + I_Ks_ models in the subthreshold regime with nonlinearities of quadratic type

The two models have *V* -nullclines of quadratic type in the subthreshold regime or are quasi-linear for the parameter sets considered (Figs. S4, and S6-A1 and -B1) (see also [70], Fig. 2, and [72], Fig. 7). However, while this is a rather general property of models of HH type having regenerative (amplifying) ionic currents (e.g., *I*_*Nap*_ or inward-rectifying potassium *I*_*Kir*_) [66, 67, 74]), we note that in certain, also rather general parameter regimes, both models can have *V* -nullclines of cubic-type [66, 67, 104] around which, not around the parabolic component of the cubic-like nullcline relevant subthreshold behavior such as subthreshold oscillations are generated.

The *I*_*Nap*_ model, which can be thought of as a 1D reduced version of the 2D models discussed above has a parabolic-like *dV/dt* vs. *V* curve.

**Figure S6:**
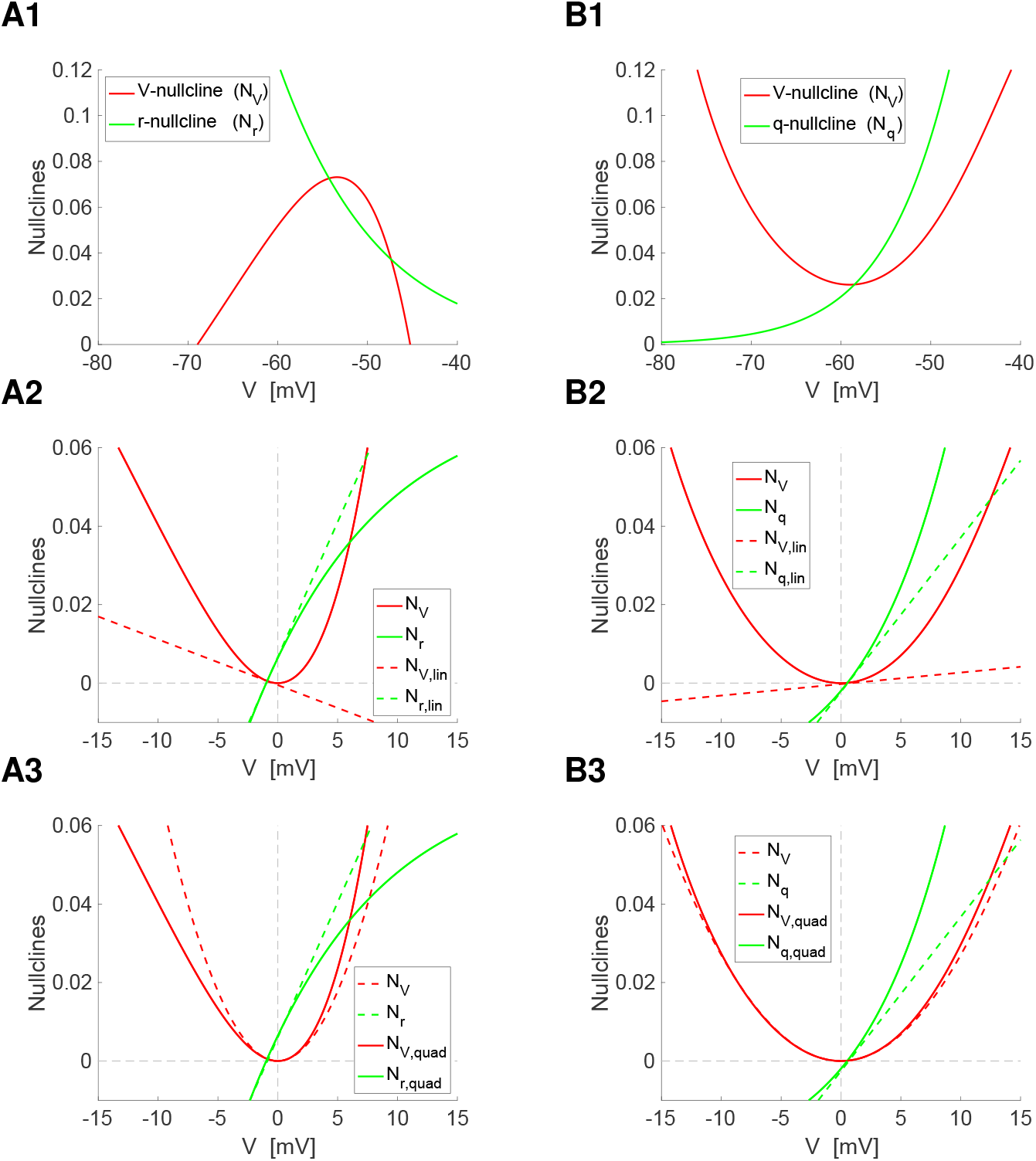
Linearization and quadratization of the *I*_*Nap*_+*I*_*h*_ and *I*_*Nap*_+*I*_*Ks*_ models with voltage nonlinearities of parabolic type in the subthreshold regime. **A.** *I*_*Nap*_+*I*_*h*_ model. We used the parameter values presented in Section S1.2 **B.** *I*_*Nap*_+*I*_*Ks*_. We used the parameter values presented in Section 4. A detailed description of the process for 3D models of HH type including time-dependent external currents and synaptic inputs is presented in [70] and also discussed in [72] in the context of reduced neuronal models. We note that while the presence of voltage nonlinearities of parabolic type are a rather general property of models of HH type having regenerative (amplifying) ionic currents (e.g., *I*_*Nap*_ or inward-rectifying potassium *I*_*Kir)*_ [66, 67, 74]), in other, also rather general parameter regimes, both models can have *V* -nullclines of cubic-type [66, 67, 104] around which, not around the parabolic component of the cubic-like nullcline relevant subthreshold behavior such as subthreshold oscillations are generated.

### S1.3 Linearization: conductance-based linearized models

Linearization of the autonomous part (*I*_*in*_(*t*) = 0) of system (S1)-(S2) around the fixed-point 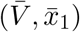 yields [20, 21, 88]

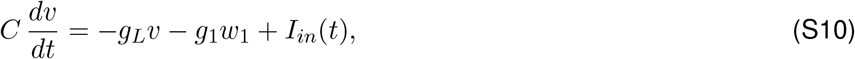

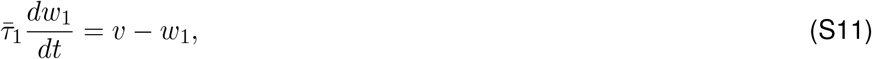

where

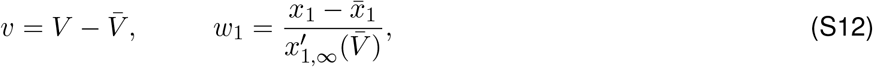

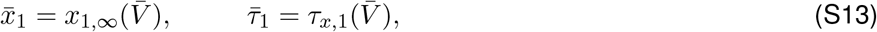

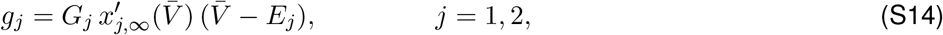

and

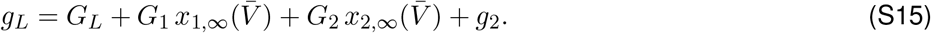

Note that the gating variables *w*_1_ and *w*_2_ in (S12) have units of voltage ([*v*] = [*w*_1_] =V). A Comparison between the original (biophysical) *I*_*Nap*_ + *I*_*h*_ and *I*_*Nap*_ + *I*_*Ks*_ models and their linearization are presented in Figs. S6-A2 and -B2 (see also [72], Fig. 7).

The effective leak conductance *g*_*L*_ (S15) contains information about the biophysical leak conductance *G*_*L*_, the ionic conductances, and their associated voltage-dependent activation / inactivation curves. The fast ionic current *I*_2_ contributes to *g*_*L*_ with an additional term *g*_2_. The signs of the effective ionic conductances *g*_*j*_ determine whether the associated gating variables are resonant (*g*_*j*_ *>* 0) or amplifying (*g*_*j*_ *<* 0) [1, 20]. Specific examples are the gating variables associated to *I*_*h*_ (resonant), *I*_*Ks*_ (resonant), *I*_*Nap*_ (amplifying) and *I*_*Kir*_ (amplifying). All terms in *g*_*L*_ are positive except for the last one that can be either positive or negative. Specifically, *g*_*L*_ can become negative for negative enough values of *g*_2_.

All the linearized conductances are affected not only by the respective biophysical conductances, but also by the magnitudes and signs of the activation (*σ <* 0) and inactivation (*σ >* 0) curves and their derivatives, which are typically given by expressions of the form

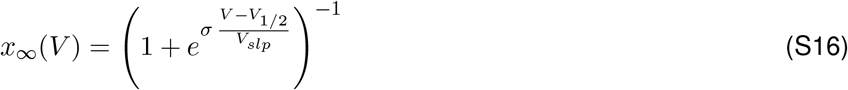

and

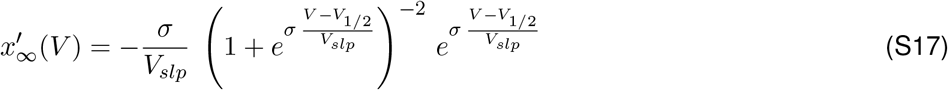

where *V*_1/2_ and *V*_*slp*_ > 0 are constants (see the examples shown in Section S1.2).

### S1.4 Quadratization: conductance-based linearized models

Quadratization of biophysically plausible models of HH type (S1)-(S2) extends the notion of linearization to include the parabolic-like properties of the *V* -nullcline in the subthreshold regime and therefore capture more realistic aspects of the dynamics of these models, which can be missed by the corresponding linearization [70–72]. In contrast to linearization, which is carried out around the model’s fixed point (determined by the interSection of the model’s nullclines), quadratization is done around the extremum (minimum or maximum) of the *V* -nullcline of quadratic type.

One important assumption is that the *V* -nullcline is parabolic-like in the subthreshold regime (e.g., Fig. 2 in [70] and Fig. 7 in [72]). This is a rather general property of neuronal models of HH type having regenerative (amplifying) ionic currents [66, 67, 74].

The process of quadratization [70, 71] consists of expanding the right-side of the model differential equations into Taylor series around the minimum/maximum (*V*_*e*_, *x*_1,*e*_) of the parabolic-like *V* -nullcline in the subthreshold regime, neglecting all the terms with power bigger than two in the equation for *V* and bigger than one in the equation for *x*_1_, and translating the minimum/maximum of the *V* -nullcline to the origin. Details are provided in [70–72] for 2D and 3D models.

The quadratized 2D model (S1)-(S2) around (*V*_*e*_,*x*_1,*e*_) is given by

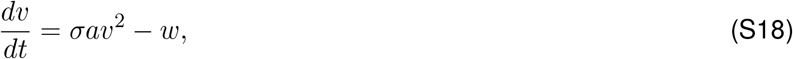

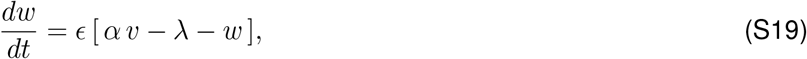

where

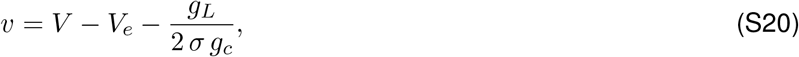

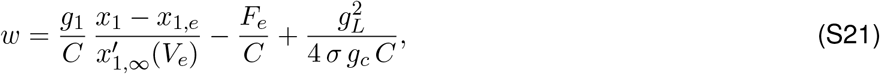

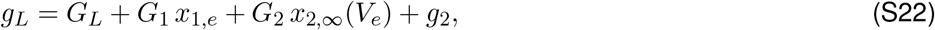

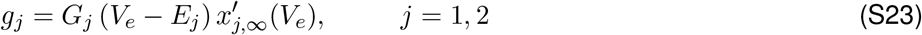

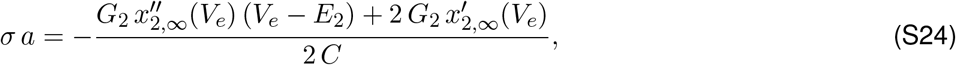

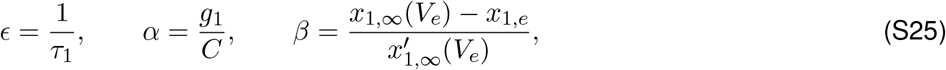

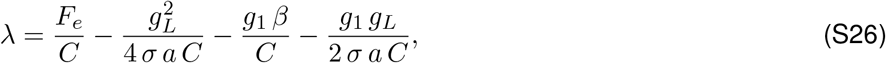

and

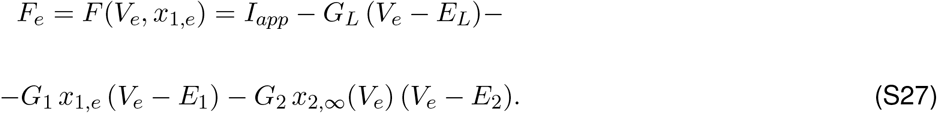

A Comparison between the original (biophysical) *I*_*Nap*_ + *I*_*h*_ and *I*_*Nap*_ + *I*_*Ks*_ models and their quadratization is presented in Figs. S6-A3 and -B3 (see also [72], Figs. 3 and 4). A detailed description of the process for 3D models of HH type including time-dependent external currents and synaptic inputs is presented in [70] and also discussed in [72] in the context of reduced neuronal models.

## S2 Impedance amplitude and phase profiles for 2D linear systems: individual cells

For a linear system

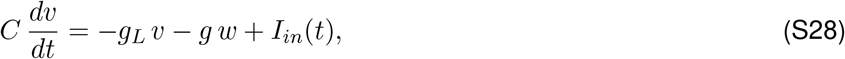

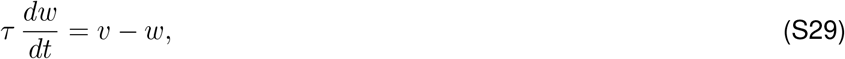

receiving a sinusoidal input current of the form (5)

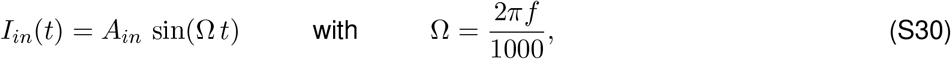

the voltage response is given by

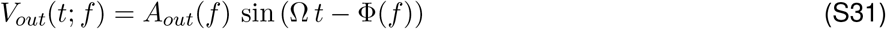

where *A*_*out*_(*f)* is the amplitude and Φ(*f)* is the phase-shift (or phase), which captures the difference between the peaks of *I*_*in*_(*t*) and *V*_*out*_(*t*; *f)* normalized by the period. For the linear system (S28)-(S29), the impedance is given by [20, 21, 88])

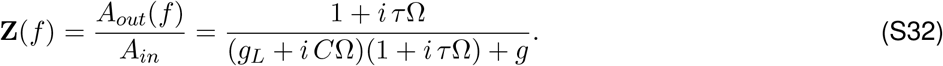

The *Z*- and Φ-profiles are given, respectively by

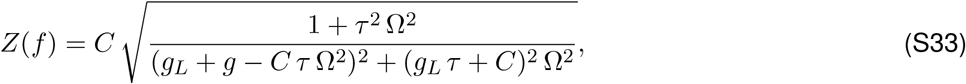

and

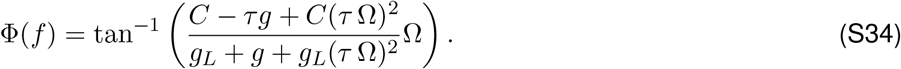

System (S28)-(S29) exhibits *resonance* (BPF) if *Z*(*f)* peaks at a non-zero (resonant) frequency *f*_*res*_ (Figs. 1-A2) and *phasonance* if Φ(*f)* vanishes at a non-zero (phasonant) frequency *f*_*phas*_ (Figs. 1-C2).

From (S33), the resonant frequency is given by

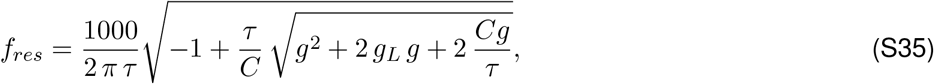

and the impedance peak *Z*_*max*_ is given by

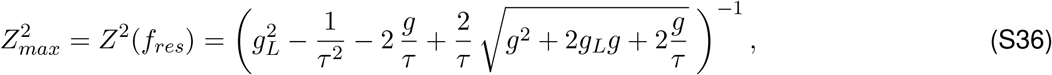

where for simplicity we used *C* = 1. From (S34), the phasonant frequency is given by

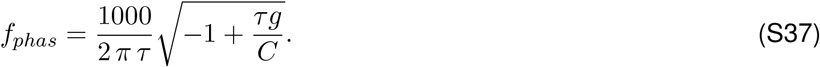

For passive (one-dimensional) cells (*g* = 0), the *Z*-profile reduces to a low-pass filter

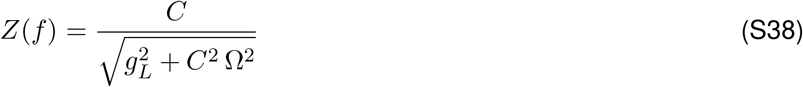

and the Φ-profile reduces to the positive increasing function

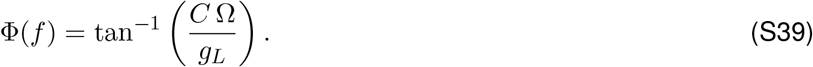

The eigenvalues for the autonomous part of system (S28)-(S29) (no inputs) are given by

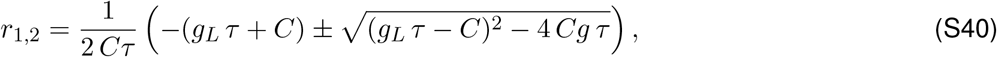

and the natural frequency, if it exists, is given by

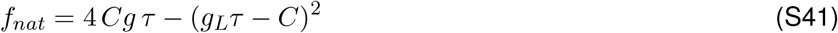

provided this quantity is positive.

## S3 Network impedance profiles for linear systems: networks with linear nodes and connectivity

We consider networks consisting of *N* nodes with 2D linear dynamics and linear connectivity. The dynamics are described by

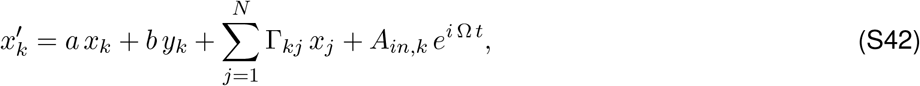

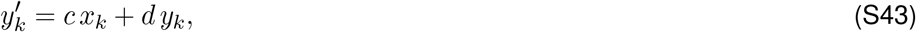

for *k* = 1, …, *N*. In (S42)-(S43), *a, b, c* and *d* are constant, Ω *>* 0 is the input frequency and *A*_*in,k*_ *≥* 0 are the input amplitudes. The connectivity coefficients Γ_*kj*_ indicate a connection from cell *j* to cell *k*. The coefficients *a, b, c* and *d* may be different in different nodes in the network. For simplicity in the notation we omit the subindex, which will be reintroduced later in the notation for the impedance profiles.

We use the notation **Z_k_**(Ω), *Z*_*k*_(Ω) and Φ_*k*_(Ω) (*k* = 1, …, *N)* to refer to the (complex) impedance, impedance amplitude (or simply, impedance) and phase profiles, respectively, of the individual nodes. The analytical expressions for 2D and 1D (*b* = 0) models are given in the Appendix S2.

The equation (S42) can be rewritten as

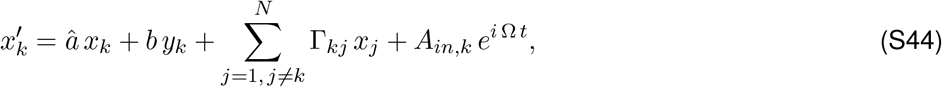

for *k* = 1, …, *N* where *â* = *a* + Γ_*kk*_ includes the coefficient (Γ_*kk*_) of the autonomous component of the coupling term (Γ_*kk*_ *x*_*k*_). We use the notation 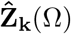 to refer to the (extended) impedance profile of the individual nodes including this term; i.e., the impedance profiles of the system consisting of the autonomous component in (S44) and equation (S43). Similarly, 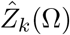 and 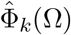 refer to the extended impedance amplitude and phase profiles of these systems, respectively.

The particular solution to system (S42)-(S43) has the form

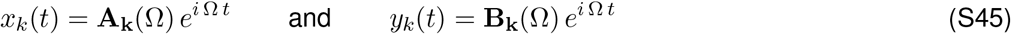

for *k* = 1, …, *N*. Substituting (S45) into (S43)-(S44) and rearranging terms we obtain

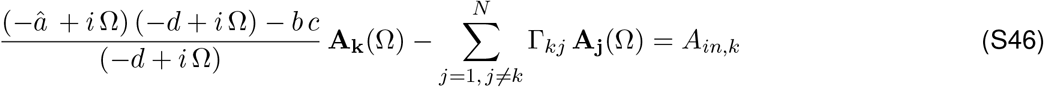

for *k* = 1, …, *N*. From (S82) in the Appendix S2 with *a* substituted by 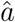 we obtain

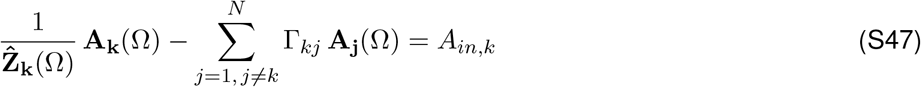

for *k* = 1, …, *N* .

This is a complex linear system of algebraic equations that can be written in compact form as

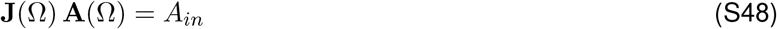

where

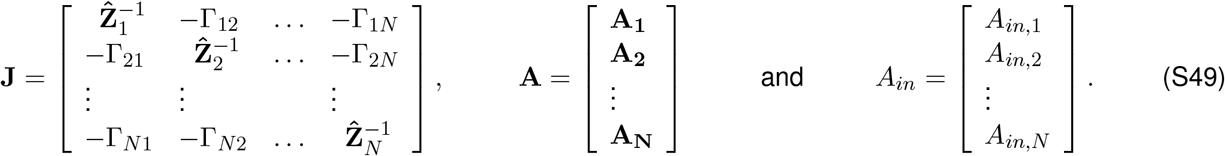

Note that for simplicity the explicit dependence of **J** and **A** on Ω has been omitted in (S49).

The solution to (S48) is formally given by

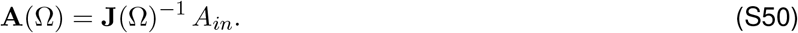

In order to invert the complex matrix **J**(Ω) one needs to express 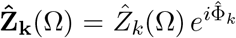 as 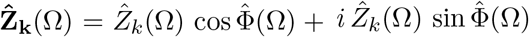 and **A_k_**(Ω) = *A*_*k,real*_(Ω) + *i A*_*k,imag*_(Ω). Alternatively, **A_k_** = *|A*_*k*_(Ω)*| e^iΨ(Ω)^* = *|A*_*k*_(Ω)*|* cos Ψ(Ω) + *i A*_*k*_(Ω) sin Ψ(Ω).

The vector **A**(Ω) represents the response of the network to the oscillatory inputs. For a two-cell network (*N* = 2) we obtain

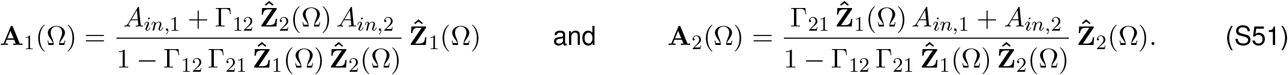

In order to extend the concept of impedance to networks we need to find an appropriate normalization. In the impedance for the individual neurons, this normalization is given by the input amplitude. If the input arrives only to one node in the network, it is natural to choose this input amplitude as the normalization constant. Otherwise, one has to choose an appropriate one among the various input amplitudes. Here we choose the largest of all and we index the sequence of input amplitudes in decreasing order of magnitude. Thus, eq. (S50) reads

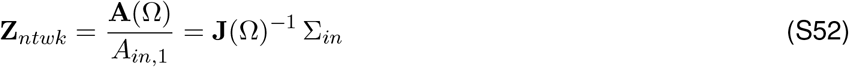

where Σ_*in*_ = [1, *σ*_2_, …, *σ*_*N*_ ]^*T*^ with *σ*_*k*_ = *A*_*in,k*_*/A*_*in,1*_ (*k* = 2, …, *N)*. For a two-cell network, we obtain eqs. (7)-(8).

Assuming we have chosen an appropriate normalization, we call the *Z*_*ntwk,k*_(Ω) and Φ_*ntwk,k*_(Ω) (*k* = 1, …, *N)* the network impedance amplitude and phases, respectively. While one could conceive other, global measures of the voltage response, such as the addition of the responses of the two cells or the quotient between the two, the one we choose is the most informative for the purpose of this paper.

### S3.1 Extended impedance

The extended 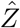 and 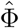 profiles are given, respectively by

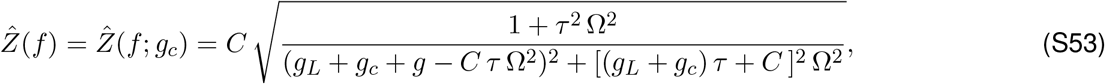

and

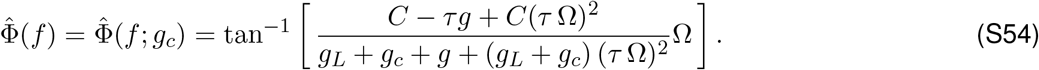

For passive (one-dimensional) cells (*g* = 0), the extended 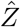 and 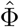 profiles reduce to

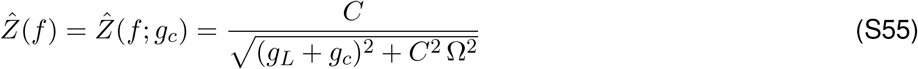

and the Φ-profile reduces to the positive increasing function

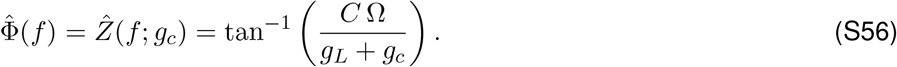

**Figure S7:**
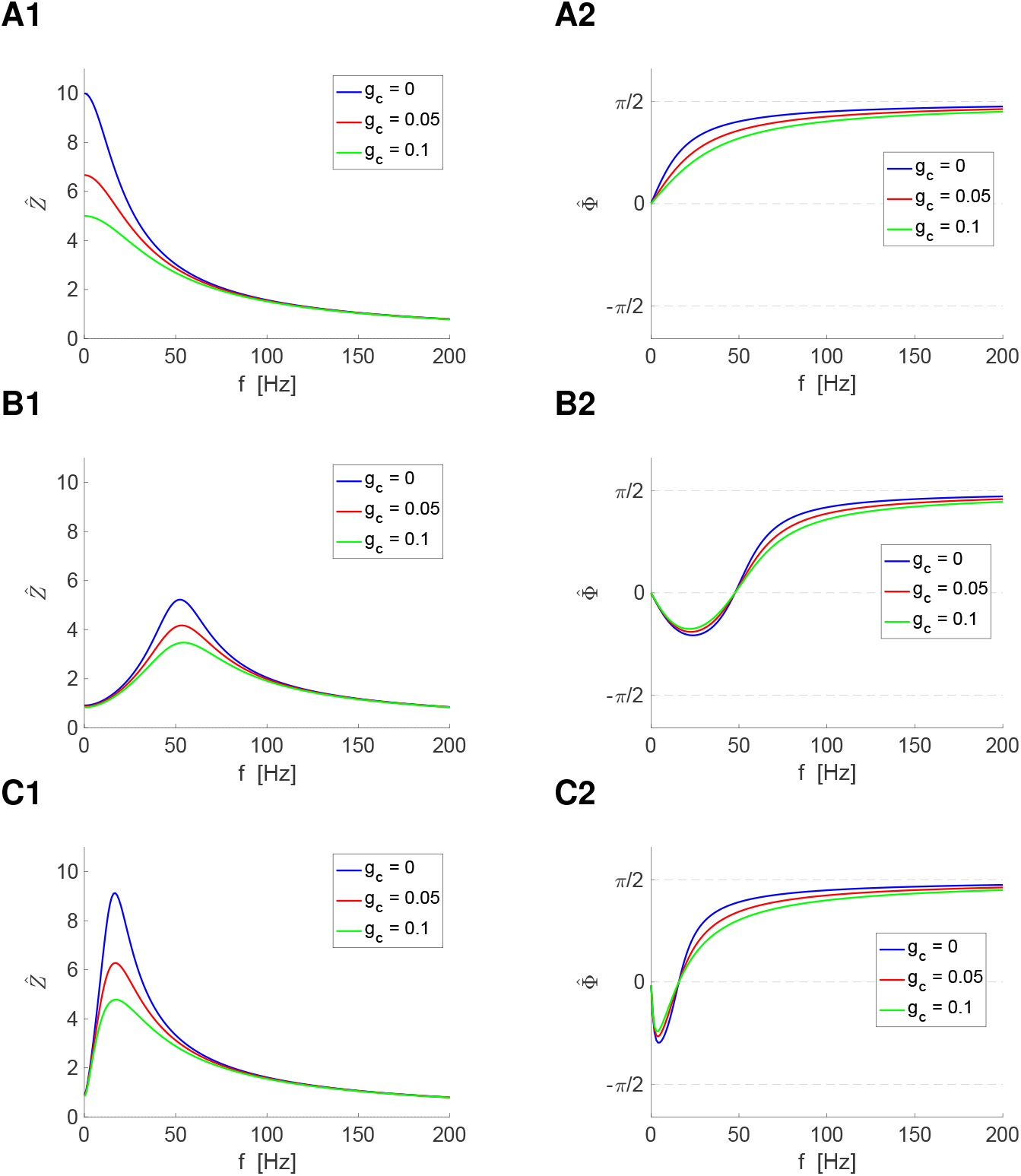
Extended impedance 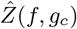 and phase 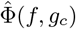 profiles. We used eqs. (S53)-(S54). For *g* = 0, these eqs. reduce to eqs. (S55)-(S56). **Left columns.** Extended impedance profiles. **Right columns.** Extended phase profiles. **A.** *g* = 0. **B.** *g* = 1, *τ* = 10. **C.** *g* = 1, *τ* = 100. We used the additional parameter value: *g*_*L*_ = 0.1.

### S3.2 Characteristic polynomial for a two-cell network

The characteristic polynomial for a two-cell network of the form (S42)-(S43) is given by

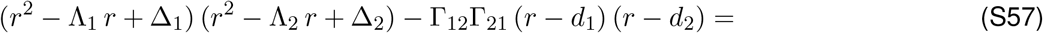

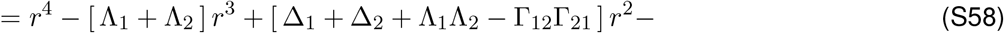

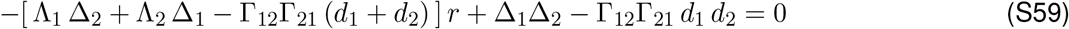

where

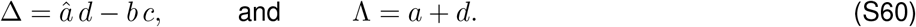

For two-cell networks of 1D cells, the characteristic polynomial reduces to

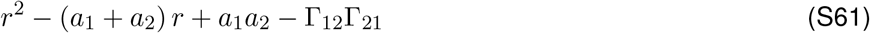

## S4 Networks of electrically coupled weakly nonlinear (quasi-linear) cells: Stationary and frequency-dependent coupling coefficient

The weakly nonlinear networks are an extension of the linear model (1)-(2) to include weak cellular nonlinearities in the current balance equation

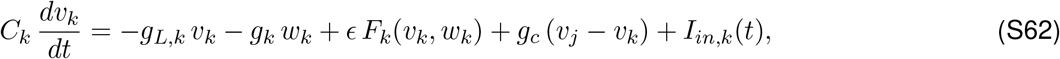

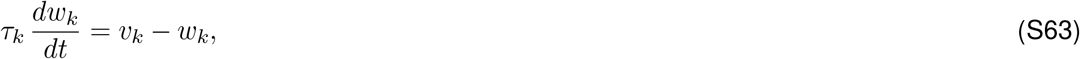

for *k, j* = 1, 2 where *ϵ* ≪ 1. The functions *F*_*k*_ are arbitrary well-behaved functions.

### S4.1 Response to constant inputs

The steady-state response of the quasi-linear system (S62)-(S63) to a constant (DC) inputs *I*_1_ (*I*_2_ = 0) can be written as

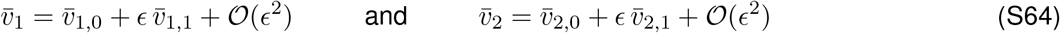

where 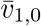 and 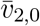 are given by (S66) and (S67), respectively. The 𝒪(*ϵ*) approximations are the solutions to

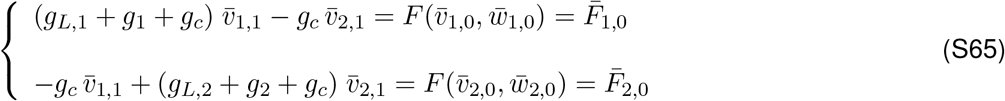

given by

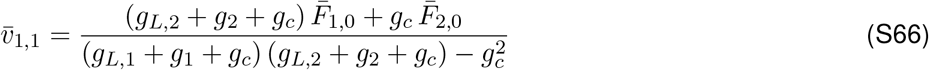

and

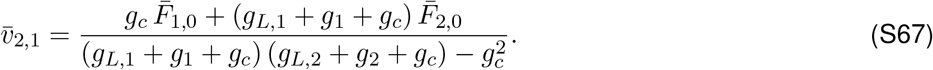

By substituting into (S64),

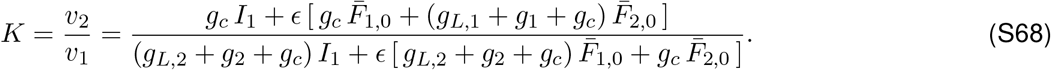

For linear models, *K* is independent of the input current and the intrinsic properties of the pre-J cell [43, 44] (see also Section 3.1.1). For networks of 2D linear cells, *K* is affected by *g*_*L,2*_ and also by *g*_2_, but it remains independent of the intrinsic properties of the pre-J cell (see Section 3.1.1). As the result of the presence of the nonlinearities (*ϵ >* 0), *K* is no longer independent of *I*_1_ and is affected by the intrinsic properties of both the pre- and post-J cells.

#### S4.2 Response to oscillatory inputs

The stationary solution of the quasi-linear system (S62)-(S63) to an oscillatory input current *I*_1_(*t*) = *A*_*in*_ *e*^*i*^ ^Ω^ ^*t*^ to cell 1 (the pre-J cell) and no input to cell 2 (the post-J cell; *I*_2_ = 0) can be written as

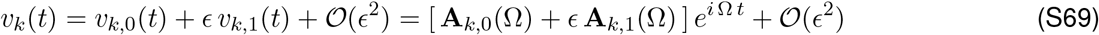

and

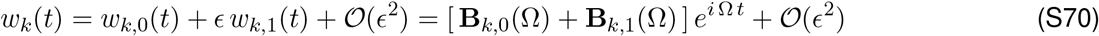

for *k* = 1, 2, where

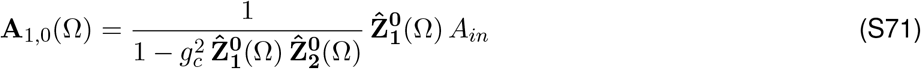

and

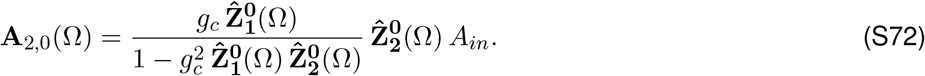

and 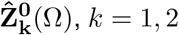 are given by (9)-(11).

For simplicity, we assume that *F*_*k*_ is independent of *w*_*k*_ (*k* = 1, 2) and we expand

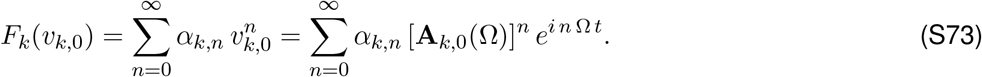

The 𝒪(*ϵ*) approximations are the stationary solutions of the following system

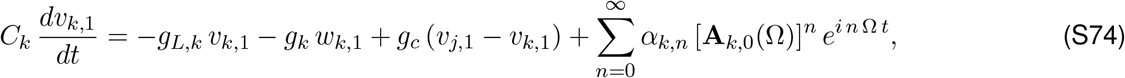

and

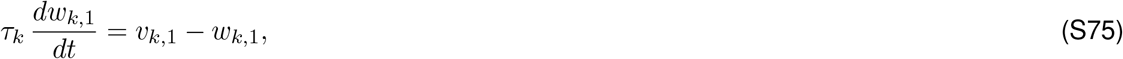

for *k, j* = 1, 2. By solving this system, one obtains

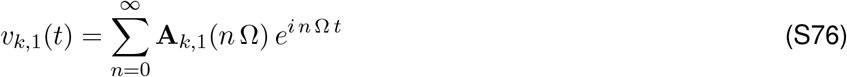

where

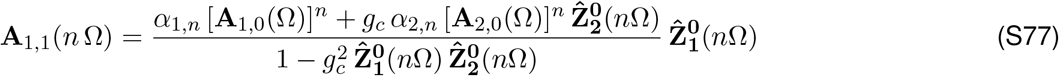

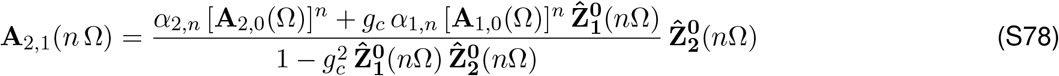

for *n* = 1, 2, The solution to the constant input term (*n* = 0) is provided in the previous section.

For linear networks, the frequency-dependent coupling coefficient is independent of the input current amplitude (*A*_*in*_) and the intrinsic properties of the pre-J cell and only depends on *g*_*c*_ and the intrinsic properties of the post-J cell via its extended impedance. As the result of the presence of cellular nonlinearities, the coupling coefficient is no longer independent of the input amplitude *A*_*in*_ and is affected by the intrinsic properties of both the pre-J and post-J cells.

## S5 Impedance amplitude and phase profiles for 2D linear systems: general calculation

In order to analytically compute the impedance amplitude and phase profiles for 2D linear systems motivated by neuronal models we use

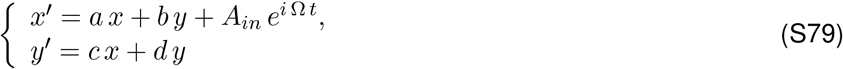

where *a, b, c* and *d* are constant, Ω *>* 0 and *A*_*in*_ *≥* 0 is the input amplitude.

### Impedance and phase profiles

The particular solution to system (S79) has the form

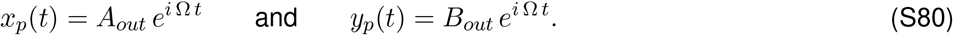

Substituting (S80) into (S79) and rearranging terms we obtain

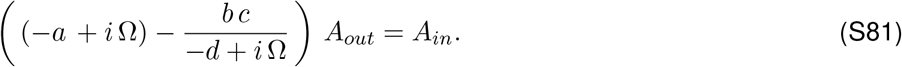

The impedance results

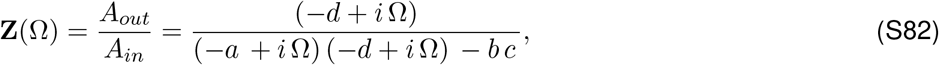

which is a complex quantity with amplitude *|***Z**(Ω)*|* and phase Φ(*f)*. For simplicity in the notation we use *Z*(Ω) for the impedance amplitude.

The impedance amplitude and phase are given, respectively, by

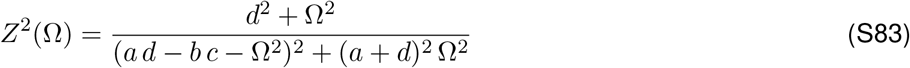

and

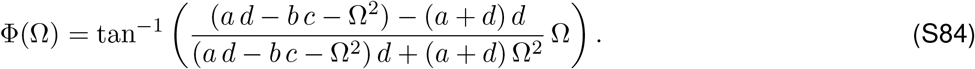

The resonance frequency, if it exists, is given by

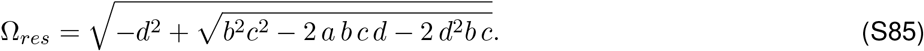

### Impedance and phase profiles: 1D linear systems

For 1D linear systems, by making *b* = 0 in (S82) the impedance **Z** reduces to

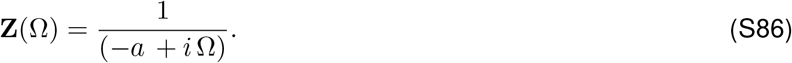

The impedance amplitude and phase are given, respectively, by

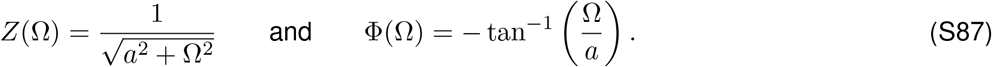

### Characteristic polynomial for the autonomous system

The characteristic polynomial for the corresponding homogeneous 2D system (*A*_*in*_ = 0) is given by

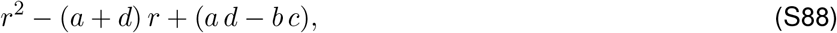

and the roots are

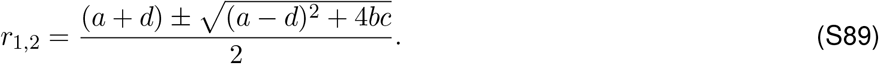

From the above equation, if 4*bc* + (*a − d*)^2^ *<* 0, the natural frequency of system (S79) is given by

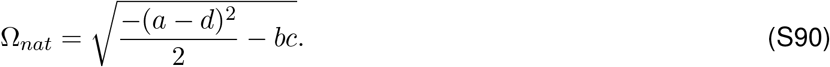

## S6 Additional Figures

**Figure S8:**
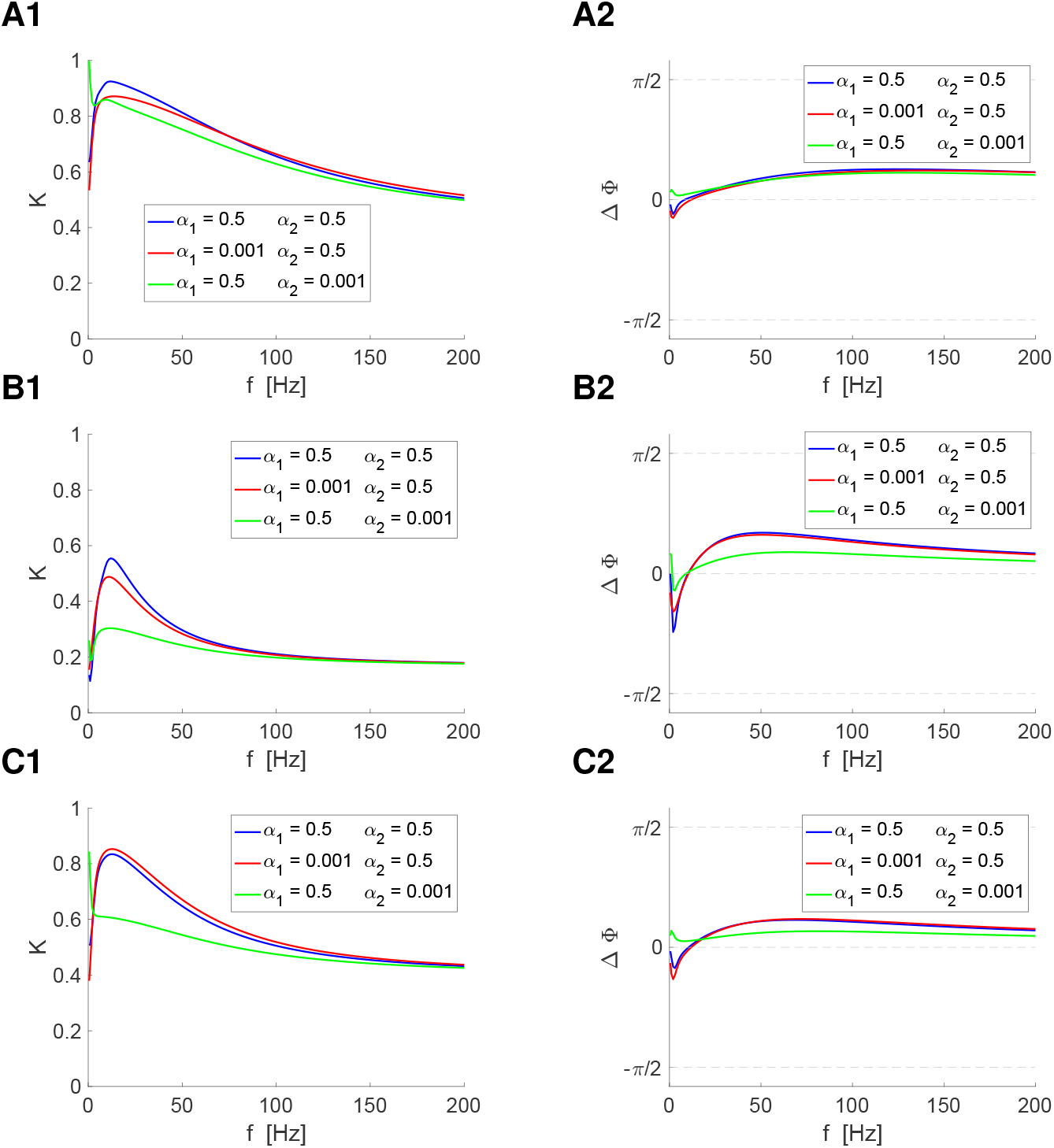
Electrically coupled quadratic 2D model: Effects of the interplay of the biophysical properties of the pre-J and post-J cells and the dendritic geometry on the response profiles. This figure is a rearrangement of Figs. 18-A, 16-A and 19-A to capture the effect of the dendritic geometry for fixed choices of the biophysical parameters. The description of the model is provided in the captions of these figures. We used *a*_1_ = *a*_2_ = 0.1, *ϵ*_1_ = *ϵ*_2_ = 0.01. **A.** Fig. 18-A (*σ* = 0.2). **B.** Fig. 16-A (*σ* = 0.5). **C.** Fig. 19-A (*σ* = 0.8).

**Figure S9:**
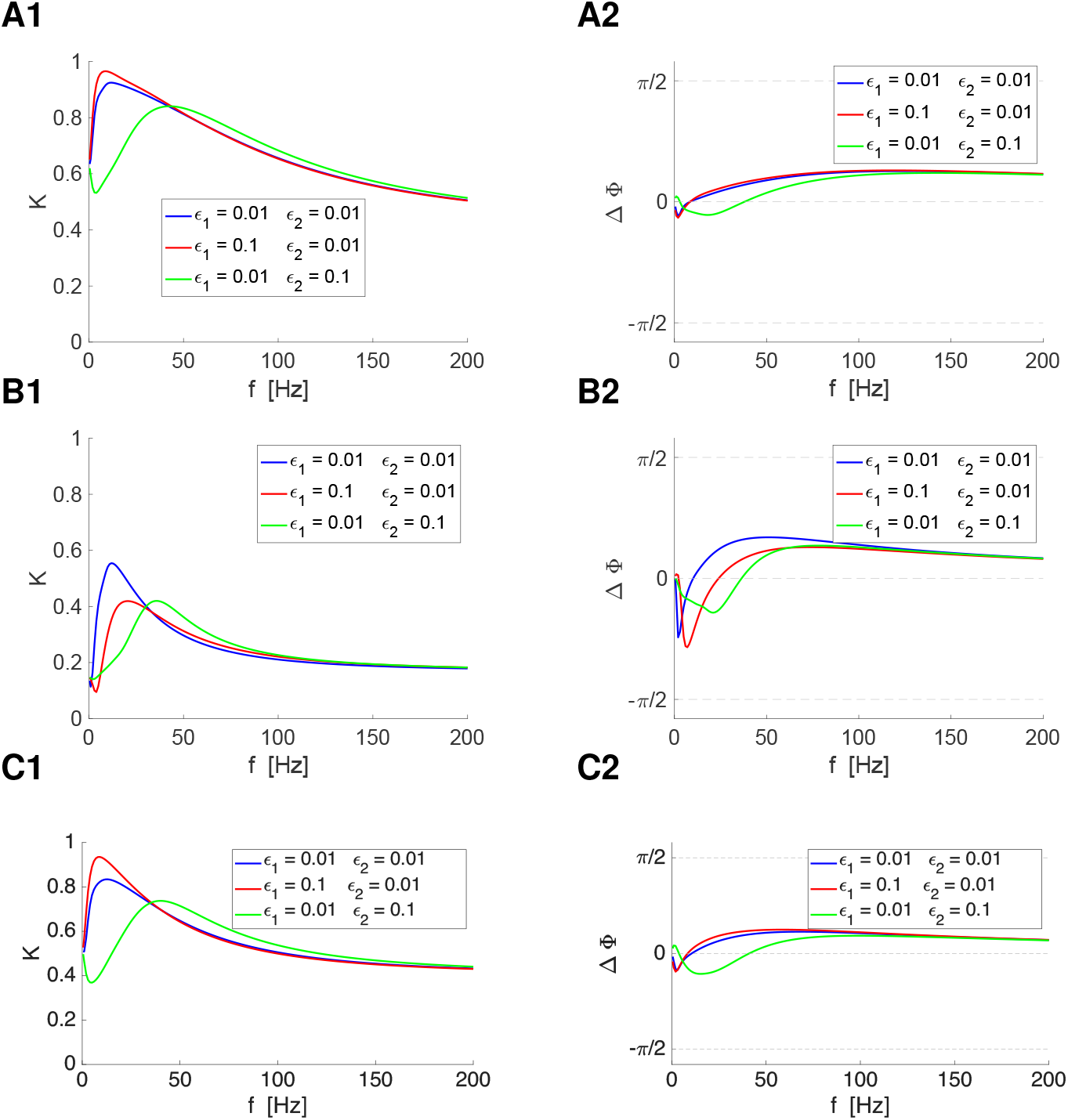
Electrically coupled quadratic 2D model: Effects of the interplay of the biophysical properties of the pre-J and post-J cells and the dendritic geometry on the response profiles. This figure is a rearrangement of Figs. 18-B, 16-B and 19-B to capture the effect of the dendritic geometry for fixed choices of the biophysical parameters. The description of the model is provided in the captions of these figures. We used *a*_1_ = *a*_2_ = 0.1, *α*_1_ = *α*_2_ = 0.5. **A.** Fig. 18-B (*σ* = 0.2). **B.** Fig. 16-B (*σ* = 0.5). **C.** Fig. 19-B (*σ* = 0.8).

**Figure S10:**
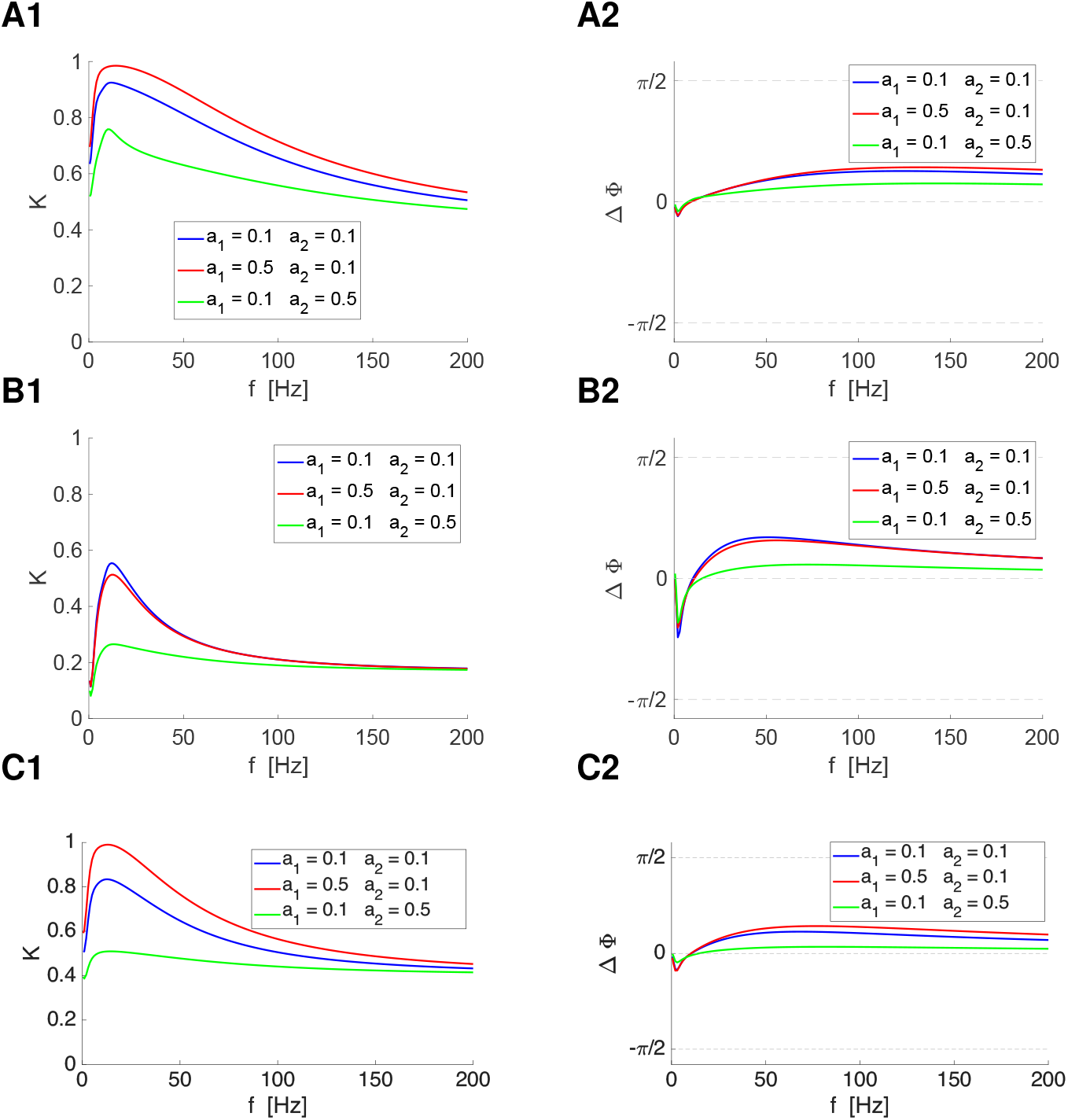
Electrically coupled quadratic 2D model: Effects of the interplay of the biophysical properties of the pre-J and post-J cells and the dendritic geometry on the response profiles. This figure is a rearrangement of Figs. 18-C, 16-C and 19-C to capture the effect of the dendritic geometry for fixed choices of the biophysical parameters. The description of the model is provided in the captions of these figures. We used *α*_1_ = *α*_2_ = 0.5, *ϵ*_1_ = *ϵ*_2_ = 0.01. **A.** Fig. 18-C (*σ* = 0.2). **B.** Fig. 16-C (*σ* = 0.5). **C.** Fig. 19-C (*σ* = 0.8). We used the following additional parameter values:, *λ*_1_ = *λ*_2_ = *−*0.2, *g*_*c*_ = 0.05, *A*_*in,1*_ = 0.06 and *A*_*in,2*_ = 0.

